# The bHLH121–CAT2 module maintains root tip redox homeostasis to confer salt tolerance additively with the SOS pathway in *Arabidopsis*

**DOI:** 10.64898/2026.11.02.763515

**Authors:** Peijun Zhou, Chuanfa Liu, Yilin Pan, Ruonan Wang, Yuchen Fei, Bingyan Liu, Xinran Du, Md. Isrfil Hossen, Yinglong Chen, Linzhou Liang, Renfang Shen, Ping Lan

**Author notes:** Correspondence (Ping Lan), (Ruonan Wang).

## Abstract

Soil salinity inhibits plant growth, in part, by restricting the availability of essential mineral nutrients such as iron (Fe). Although Fe deficiency frequently coincides with salt stress in the field, the molecular mechanisms integrating Fe homeostasis with salt tolerance remain largely unknown. Here, we identify the Fe deficiency response regulator bHLH121 as a key coordinator of salt tolerance in Arabidopsis. Loss-of-function mutants of bHLH121 exhibit pronounced hypersensitivity to salt stress, with severely impaired primary root elongation and reduced seedling biomass. Transcriptomic and histochemical analyses reveal that bHLH121 is required to maintain reactive oxygen species (ROS) homeostasis specifically in root tips under salt stress. We further demonstrate that *CATALASE 2* (*CAT2*) is a direct downstream target of bHLH121, and that *CAT2* overexpression fully rescues the salt hypersensitivity of the *bhlh121* mutant. Importantly, this bHLH121–CAT2 module operates independently of the classical SOS pathway, as *sos2 bhlh121* double mutants display additive salt sensitivity. Together, our findings uncover a novel regulatory module in which bHLH121 governs root tip redox homeostasis through CAT2 to confer salt tolerance, providing a direct mechanistic link between the Fe regulatory network and abiotic stress resilience in plants.

## 1 Introduction

Soil salinization is one of the major abiotic constraints limiting crop productivity worldwide. The damage imposed by salt stress on plants is multifaceted, and disruption of cellular ionic homeostasis is particularly critical (Mittler et al., 2022; Yang et al., 2018; Zhou et al., 2024). In the model plant *Arabidopsis thaliana*, one of the central mechanisms underlying salt stress responses is the SOS (Salt Overly Sensitive) signaling pathway (Ma et al., 2026). This pathway comprises the calcium sensor SOS3, the protein kinase SOS2, and the plasma membrane Na^+^/H^+^ antiporter SOS1. Upon salt stress, cytosolic Ca^2+^ signals are perceived by SOS3, which recruits and activates SOS2; the SOS3–SOS2 complex then phosphorylates and activates SOS1, driving Na^+^ efflux and maintaining cellular Na^+^/K^+^ balance. The discovery of the SOS pathway laid the foundation for understanding the molecular basis of plant salt tolerance, and the functional conservation of its components has been documented in a wide range of crop species.

However, the deleterious effects of salt stress are not restricted to ionic imbalance (van Zelm et al., 2020; Yang et al., 2018). A common metabolic consequence is the overaccumulation of reactive oxygen species (ROS), which cause oxidative damage to proteins, DNA, and membrane lipids (Mittler et al., 2022; Yang et al., 2018; Zhou et al., 2024). Under salt stress, stomatal closure restricts CO_2_ availability, and photosystems are directly damaged, leading to over-reduction of the photosynthetic electron transport chain. This promotes substantial electron leakage to O_2_, generating superoxide anions (O_2_^•−^) within chloroplasts (Castro et al., 2021; Miller et al., 2010; van Zelm et al., 2020). Parallel disruption of mitochondrial respiration further contributes to O_2_^•−^ production via electron leakage under ionic imbalance. Apoplastic ROS generation is simultaneously enhanced through the activation of NADPH oxidases (RBOHs). Critically, this elevated ROS production coincides with a compromised antioxidant system, in which suppression of key enzymes such as catalase (CAT) dampens hydrogen peroxide (H_2_O_2_) scavenging capacity. Together, these disruptions overwhelm cellular redox homeostasis, leading to oxidative damage and the progression of stress injury. Therefore, the establishment of salt tolerance depends not only on the maintenance of ionic homeostasis but also on the precise control of redox homeostasis.

Iron (Fe) occupies a particularly important position within this redox regulatory network. As a cofactor for numerous enzymes involved in photosynthesis, respiration, and redox reactions, Fe is essential for plant growth and development (Briat et al., 2015; Li et al., 2022). Consequently, misregulation of Fe metabolism under stress conditions can exacerbate ROS production and lead to oxidative damage (Gupta et al., 2024; Hansch et al., 2009). In saline soils, high pH and competitive effects of Na^+^ often impair Fe acquisition, creating a combined “salt stress–Fe deficiency” scenario. Notably, multiple lines of evidence demonstrate that Fe deficiency-responsive genes can simultaneously influence Fe nutrition and salt tolerance. The Fe deficiency-responsive gene *Femu2* was reported to be induced by both iron deficiency and salt stress in *Chlamydomonas reinhardtii*; its overexpression significantly enhanced salt tolerance, whereas silencing increased salt sensitivity, accompanied by decreased proline and soluble sugar contents and elevated MDA levels (Li et al., 2017). The apple bHLH transcription factor MxbHLH18 was shown to be induced by both salt and Fe stresses, and its overexpression in *Arabidopsis* enhanced tolerance to salt, high Fe, and low Fe, with increased SOD, POD, and CAT activities, reduced MDA, H_2_O_2_, and O_2_^•−^ contents, and upregulated expression of both salt-responsive and iron-responsive genes (Liang et al., 2022). Most recently, the soybean ferritin gene *GmFER1* was demonstrated to enhance salt tolerance through a coordinated “Fe–antioxidant–ion homeostasis” regulatory network: overexpression of *GmFER1* promoted Fe^3+^ accumulation, enhanced SOD and CAT activities, and upregulated *GmSOS1*-mediated Na^+^ efflux, while simultaneously improving yield through enhanced photosynthetic efficiency and Rubisco activity (Zhang et al., 2025). Collectively, these findings demonstrate that Fe homeostasis components can coordinately regulate both Fe nutrition and salt tolerance across diverse plant species, suggesting an intrinsic link between the Fe homeostasis regulatory network and salt stress responses. However, the molecular mechanism by which the Fe regulatory machinery participates in the maintenance of redox homeostasis under salt stress remains largely unclear.

In *Arabidopsis*, the core transcriptional network governing Fe homeostasis is orchestrated by basic helix-loop-helix (bHLH) transcription factors (Gao et al., 2024). FIT, the master regulator of Fe acquisition, forms heterodimers with subgroup Ib bHLH proteins including bHLH38/39/100/101 to orchestrate the Fe deficiency response (Colangelo et al., 2004). This module is positively regulated by upstream subgroup IVc bHLH factors, including bHLH34, bHLH104, ILR3/bHLH105, and bHLH115 (Li et al., 2016; Liang et al., 2017; Wang et al., 2013; Yuan et al., 2008; Zhang et al., 2015). Notably, the Upstream Regulator of IRT1 (URI)/bHLH121 emerges as a central, bridging factor that physically interacts with bHLH IVc proteins and facilitates the expression of *FIT* and *bHLH38/39/100/101* by bHLH IVc transcription factors (Gao et al., 2024; Gao et al., 2020; Kim et al., 2019; Lei et al., 2020). This regulatory circuit is negatively controlled by E3 ubiquitin ligases; BTS targets bHLH IVc factors for degradation (Selote et al., 2015), while BTSL1 and BTSL2 mediate FIT turnover (Rodríguez-Celma et al., 2019).

The choice of bHLH121 as a focal point of this study was based on several lines of evidence. First, bHLH121 occupies a central position in the Fe regulatory network, serving as a key node connecting upstream signals to downstream responses. Second, in peanut (*Arachis hypogaea*), *AhbHLH121* overexpression enhances antioxidant enzyme activity and salt tolerance (Zhao et al., 2024). Similarly, bHLH121 was identified as a transcription factor regulating the high-affinity K^+^ transporter HAK5, with its expression induced by both low K^+^ and salt stress (Hong et al., 2013). Moreover, bHLH121 (URI) acts as an essential component of the Fe deficiency signaling pathway (Kim et al., 2019). Through re-analysis of the ChIP-seq data from Kim et al. (2019), we found that the direct binding targets of bHLH121 include ROS-related genes, suggesting a potential link between bHLH121 and redox regulation. Overall, these findings suggest that bHLH121 may act as a critical hub coordinating Fe homeostasis and salt stress responses. However, the function of bHLH121 in *Arabidopsis* salt stress responses has not been reported, and whether and how it contributes to salt tolerance through the regulation of redox homeostasis remains to be elucidated.

To address this, we assessed the expression profile of *bHLH121* and confirmed its salt responsiveness. Consistently, *bhlh121* loss-of-function mutants displayed hypersensitivity, with substantially reduced root elongation and lower seedling fresh weight. Transcriptome profiling revealed that *bHLH121* is required for the proper expression of a suite of genes involved in redox homeostasis, and ROS staining combined with pharmacological assays confirmed that bHLH121 regulates redox homeostasis, specifically within root tips. Notably, overexpression of *CAT2*, a representative H_2_O_2_-scavenging gene, rescued the salt hypersensitivity of the *bhlh121* loss-of-function mutant, and bHLH121 was shown to bind the *CAT2* promoter. Collectively, these findings establish the bHLH121–CAT2 module as a crucial regulator of redox homeostasis in plant resilience to salt stress.

## 2 Materials and methods

### 2.1 Plant materials and growth conditions

*Arabidopsis thaliana* ecotype Columbia-0 (Col-0) plants were used as the wild-type (WT). The *sos2-T1* mutant was described previously (Ma et al., 2025). The *bhlh121-7* and *bhlh121-8* mutants, the *pro35S::bHLH121:GFP/WT* overexpression lines, and the complemented lines including *probHLH121::gbHLH121:GFP/bhlh121-7, pro35S::PtbHLH121:GFP/bhlh121-7, pro35S::GsbHLH121:GFP/bhlh121-7, pro35S::CcbHLH121:GFP/bhlh121-7,* and *pro35S::CsbHLH121:GFP/bhlh121-7*) were generated as described (Zhou et al., 2026). The *sos2-T1* and *bhlh121-7* mutants were crossed to generate the *sos2 bhlh121* double mutant.

*Arabidopsis* seeds were sterilized by washing first with 75% ethanol for 3 min, followed by 0.5% (v/v) NaClO solution containing 0.5% (v/v) Tween 20 for 10 min, and finally rinsing five times with sterile deionized water. After stratification at 4 °C for 2 d, sterilized seeds were sown on ES medium (Estelle et al., 1987) and grown in a greenhouse under a 22 °C under a 16-h light (*c.* 100 μmol·m^-2^·s^-1^ over the waveband 400–700 nm) /8-h dark photoperiod at 20 °C.

The ES medium contained 5 mM KNO_3_, 2 mM Ca(NO_3_)_2_·4H_2_O, 2 mM MgSO_4_·7H_2_O, 2.5 mM KH_2_PO_4_, 40 μM Fe(III)-EDTA, 14 μM MnCl_2_·4H_2_O, 70 μM H_3_BO_3_, 1 μM ZnSO_4_·7H_2_O, 0.5 μM CuSO_4_·5H_2_O, 0.2 μM NaMoO_4_·2H_2_O, and 10 μM NaCl, and was used as the control treatment. For salt stress treatments, NaCl was added at 75, 100, or 150 mM. For pharmacological treatments, diethyldithiocarbamic acid (DIECA) at 25, 50, and 100 μM was used to inhibit jasmonic acid (JA) biosynthesis, while indole-3-acetic acid (IAA) at 10, 50, and 100 nM was applied to simulate auxin effects. Additionally, 500 μM potassium iodide (KI) and 200 μM glutathione (GSH) were supplemented as ROS scavengers to alleviate oxidative stress. All media contained 1% sucrose (w/v), 0.1% MES (w/v), and 0.8% agar (w/v), with the pH adjusted to 5.5.

### 2.2 Plasmid construction and generation of transgenic plants

The full-length coding sequence of *CAT2* was amplified by PCR and cloned into the modified binary vector *pCAMBIA1300-GFP*, driven by the cauliflower mosaic virus (CaMV) 35S promoter. The resulting recombinant plasmid *pro35S::CAT2:GFP* was transformed into *Agrobacterium* strain GV3101. Subsequently, *Arabidopsis* plants were transformed using the floral dip method. T0 transgenic seeds were screened on half-strength Murashige & Skoog medium containing 25 mg L^−1^ hygromycin. Three independent stable lines were selected for further analyses. Primers are listed in Supplementary Table S1.

### 2.3 Root length and fresh weight analyses

For the root elongation and fresh weight assays, *Arabidopsis* seeds were germinated and grown vertically on ES medium with or without NaCl supplementation. After 10 d of cultivation in a greenhouse, seedlings were photographed to document their phenotypes. Primary root lengths were quantified from at least 20 seedlings per biological replicate using ImageJ software (National Institutes of Health, USA). Fresh weight was determined from pools of 10 seedlings. All experiments included three biological replicates, and representative data from one replicate are shown in the figures.

### 2.4 Measurement of chlorophyll content

Chlorophyll was extracted from *Arabidopsis* shoots with 80% acetone in the dark until tissues were fully bleached. The absorbance of the supernatant was read at 645 and 663 nm on a SpectraMax Plus 384 spectrophotometer (Molecular Devices). Total chlorophyll was quantified as (20.2 × A645 + 8.02 × A663) × V/FW and expressed in mg·g^-1^ FW, as described previously (Arnon, 1949).

### 2.5 Proline and malondialdehyde (MDA) assays

Measurements were performed as described (Hodges et al., 1999), with minor modifications. Briefly, seeds were sown on medium with or without 75 mM NaCl and grown for 10 d, after which shoot and root tissues were harvested separately. Crude protein extracts were prepared using 100 mM PBS (pH 7.8).

For proline determination, 150 µL of crude protein extract was mixed with 1 mL of reaction solution containing 0.75% (w/v) sulfosalicylic acid, 25% (v/v) glacial acetic acid, and 1.25% (w/v) acid ninhydrin, prepared in 100 mM PBS (pH 7.0). The mixture was incubated in a boiling water bath for 15 min, and absorbance was measured at 520 nm using a spectrophotometer. Proline content was calculated based on a proline standard curve.

For MDA measurement, 100 µL of crude protein extract was added to 1 mL of 0.25% thiobarbituric acid solution prepared in 10% trichloroacetic acid. After boiling for 15 min, absorbance was measured at 532 nm and 600 nm. MDA content was calculated using the formula: MDA (nmol·g^−1^ FW) = (A532-A600) × Vr × (V/Vt)/155 × 1000/ FW, where Vr is the total volume of the reaction mixture, V is the total volume of the protein extract, Vt is the volume of protein extract used in the assay, and FW is the sample fresh weight.

### 2.6 Electrolyte leakage assay

The electrolyte leakage (EL) assay was performed as described previously (Panter et al., 2019) with modifications. Nine-day-old seedlings grown on ES medium were transferred to fresh medium with or without 100 mM NaCl and grown for an additional 5 d. Shoots and roots were harvested separately for analysis. Briefly, 0.1 g of fresh tissue was placed in a tube containing 5 mL of ultrapure water and shaken at room temperature for 3 h. Initial conductivity (C1) was measured using a conductivity meter. Samples were then boiled for 20 min, cooled to room temperature, and final conductivity (C2) was measured. Relative electrolyte leakage was calculated as (C1/C2) × 100%.

### 2.7 Determination of Na^+^ and K^+^ content

Nine-day-old seedlings grown on ES medium were transferred to medium supplemented with 100 mM NaCl and treated for 5 d. Shoot and root tissues were then harvested separately and dried in an oven at 65 °C. The dried tissues were digested with concentrated nitric acid at 150 °C. Na^+^ and K^+^ contents were determined using a flame photometer (FP6431, YOKE) as described previously (Karimi et al., 2021).

### 2.8 Measurement of H_2_O_2_ content and antioxidant enzyme activities

The activities of catalase (CAT), peroxidase (POD) and superoxide dismutase (SOD), as well as the content of hydrogen peroxide (H_2_O_2_), were measured using commercially available kits (Geruisi-bio, G0105W, G0107W, G0101W, and G0168W, respectively). Seven-day-old seedlings grown on ES medium were transferred to fresh medium with or without 100 mM NaCl and grown for an additional 7 d. Roots were harvested for analysis.

### 2.9 Quantitative real-time PCR (RT-qPCR) analysis

TRIzol reagent (Vazyme) was employed for total RNA extraction in accordance with the manufacturer’s recommendations. Reverse transcription of 1 µg of RNA into cDNA was carried out using the PrimeScript™ RT reagent Kit with gDNA Eraser (TaKaRa). Following a 12-fold dilution, the cDNA products were used as templates for RT-qPCR on a PIKOREAL 96 platform (Thermo Fisher Scientific) with the SYBR® Premix Ex Taq™ II (Tli RNaseH Plus) kit (TaKaRa). The amplification procedure was initiated at 95°C for 7 min, then cycled 40 times at 95 °C for 5 s and 60 °C for 30 s. The *TUA3* gene (At5g19770) served as the normalization standard, and relative transcript abundances were calculated via the 2^−ΔΔCT^ method. Primers used are listed in Supplementary Table S1.

### 2.10 Messenger RNA (mRNA)-seq and transcriptome analyses

Seedlings were grown on ES medium with or without 100 mM NaCl. Total RNA was extracted from the roots of 10-day-old seedlings. mRNA-seq was performed by Novogene Bioinformatics Technology Co., Ltd. (Beijing, China) and sequenced on a NovaSeq 6000 platform (Illumina) to generate 150-bp paired-end reads. Data analysis was performed as described previously with minor modifications (Xue et al., 2021). Briefly, clean data were obtained by removing reads containing adapters, reads with a high proportion of uncertain bases (≥10%) and low-quality reads from the raw data. The clean reads were then mapped to the *Arabidopsis* Araport11 reference genome (Cheng et al., 2017) using HISAT2, and differentially expressed genes (DEGs) were identified using Cufflinks. Significantly up- and down-regulated DEGs were filtered by fold change > 1.5 and q value < 0.05. Gene Ontology (GO) enrichment analyses and heatmaps of gene expression profiles were generated using the OmicShare tools platform (http://www.omicshare.com/tools).

### 2.11 DAB and NBT staining

Diaminobenzidine (DAB) and nitroblue tetrazolium (NBT) were used for in situ staining of hydrogen peroxide (H_2_O_2_) and superoxide anions (O_2_^•**−**^) in seedlings, respectively (Dai et al., 2022). Seedlings were incubated in DAB staining solution (1 mg·mL^−1^, pH 5.8) or NBT staining solution (0.5 mg·mL^−1^, pH 7.6) for 60 min or 30 min, respectively, followed by the removal of the staining solution and three rinses with deionized water. Then, 1 mL of anhydrous ethanol was added for decolorization. After decolorization was complete, seedlings were observed and photographed under a Zeiss Stemi 305 microscope. DAB and NBT intensities were acquired in ImageJ using the segmented line tool, with line thickness sufficient to cover half the width of the root. Staining was quantified based on the gray values of linear ROIs.

### 2.12 Measurement of root apical meristem size

The root apical meristem was determined as previously described (Dello Ioio et al., 2007). In brief, seedlings were grown vertically on the indicated medium, and roots of 10-day-old seedlings were imaged using a confocal laser scanning microscope (STELLARIS 5, Leica). Before imaging, roots were stained with propidium iodide (PI) for 10 min. PI fluorescence was detected using excitation with an argon laser at 488 nm, with an emission filter set between 633 and 695 nm. Cortex cells were counted from the QC to the first elongated cell to determine meristem cell number and length.

### 2.13 Electrophoretic mobility shift assay (EMSA)

EMSA was carried out using the Chemiluminescent EMSA Kit (Beyotime, China) according to the manufacturer’s protocol. The recombinant bHLH121 protein was expressed in *E. coli* and purified as described previously (Zhou et al., 2026). The DNA fragments of the *CAT2* promoter were synthesized and labeled with biotin at their 5′ ends. Unlabeled probes, either identical or mutated in sequence, were included as specific competitors, and a 6xHis-tagged trigger factor (His-TF) protein preparation from *E. coli* transformed with the empty vector was used as the negative control. Probes used in these assays are listed in Supplementary Table S1.

### 2.14 Statistical analysis

All data are presented as means ± standard deviation (SD). Statistical analyses and graph generation were performed using GraphPad Prism 9.5. Differences between two groups were assessed using Student’s t-test, with asterisks indicating significance (\**P* < 0.05, \*\**P* < 0.01, \*\*\**P* < 0.001, \*\*\*\**P* < 0.0001). For multiple comparisons, one-way ANOVA followed by Tukey’s post hoc test was performed, with different letters indicating significant differences (*P* < 0.05). Each experiment included at least three independent biological replicates.

### 2.15 Accession numbers

Sequence data from this article can be found in the Arabidopsis Genome Initiative or GenBank/EMBL databases under the following accession numbers: bHLH121 (AT3G19860), SOS1 (AT2G01980), SOS2 (AT5G35410), IRT1 (AT4G19690), FRO2 (AT1G01580), PLT1 (AT3G20840), PLT2 (AT1G51190), RBOHD (AT5G47910), PRX8 (AT1G34510), CAT2 (AT4G35090), JAZ1 (AT1G19180), JAZ10 (AT5G13220), GH3.15 (AT5G13370), TUA3 (AT5G19770), PtbHLH121 (Potri.004G168100), GsbHLH121 (XP_028243911), CcbHLH121 (ESR36836), and CsbHLH121 (KGN61896).

## 3 Results

### 3.1 Loss of function *bhlh121* mutations lead to salt hypersensitivity in *Arabidopsis*

To investigate whether *bHLH121* is involved in plant salt stress tolerance, we first analyzed its expression in response to NaCl treatment. These analyses identified bHLH121 as a candidate salt-responsive gene, as its expression was significantly induced by salt stress in a dose- and time-dependent manner (Supplementary Fig. S1). We previously generated two CRISPR-Cas9-mediated knockout mutants in the WT background, designated *bhlh121-7* and *bhlh121-8*; these mutants exhibit severe growth defects and impaired Fe deficiency-responsive gene expression under Fe depletion (Zhou et al., 2026). To dissect the functional role of bHLH121 in salt tolerance, we germinated these two mutants and examined their growth phenotypes at 75, 100, and 150 mM NaCl. Both mutants displayed salt hypersensitivity, as evidenced by significantly reduced root elongation (Supplementary Fig. S2). Because both mutants exhibited similar salt-sensitive phenotypes, we selected *bhlh121-7* as a representative mutant for subsequent analyses.

A native promoter-driven bHLH121-GFP fusion protein successfully restored root growth and biomass in the *bhlh121-7* mutant under salt stress (Supplementary Fig. S3), validating the functionality of bHLH121-GFP and confirming that *bHLH121* loss is causal for the salt-sensitive phenotype. Intriguingly, although the loss-of-function *bhlh121-7* mutant displayed pronounced salt sensitivity, 35S promoter-driven overexpression of *bHLH121* in WT plants failed to enhance salt tolerance, as no significant differences in root growth or biomass were observed compared to WT (Fig. 1A–C). These results support the notion that *bHLH121* function is indispensable for salt stress tolerance in *Arabidopsis*.

**Figure 1.**
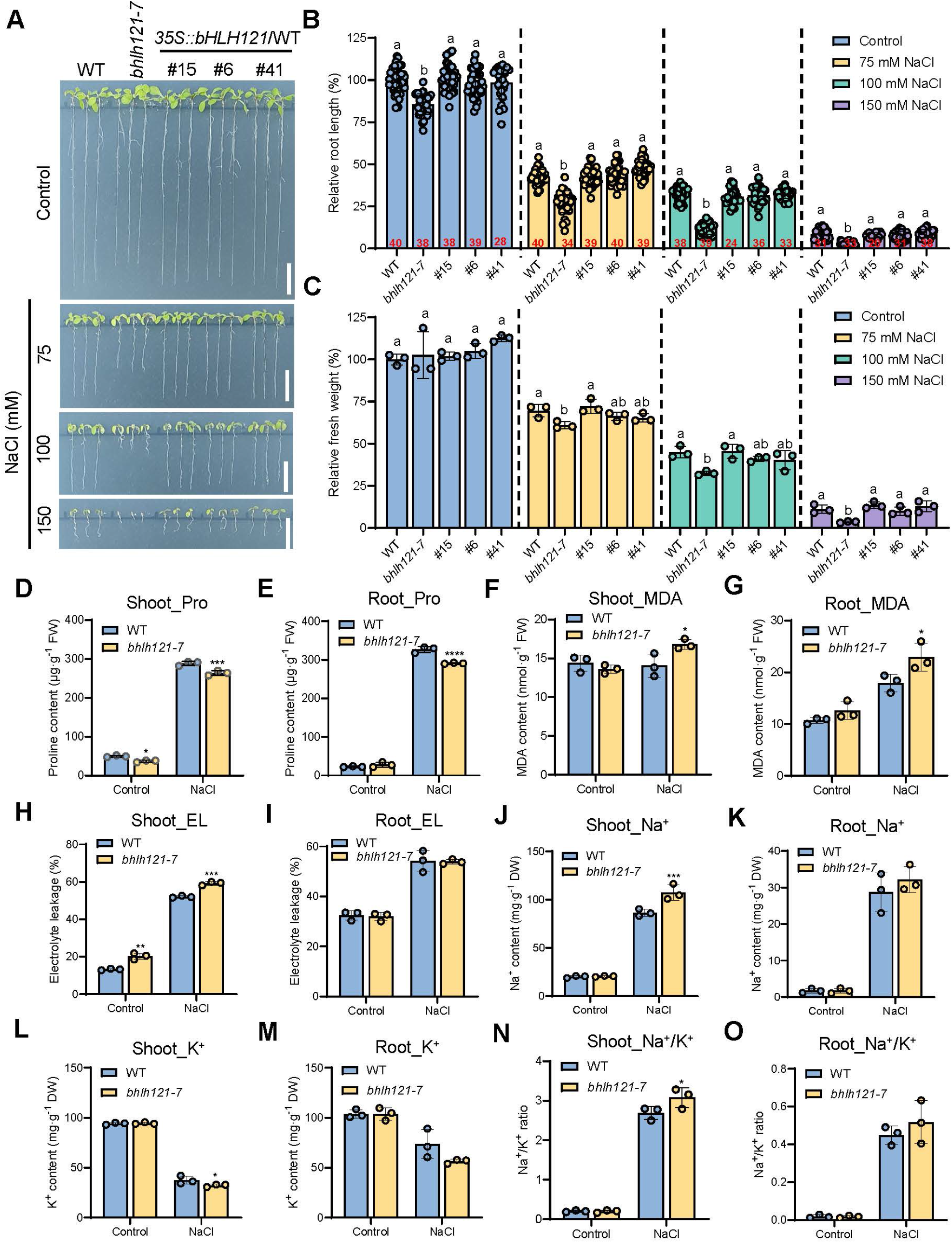
The *bhlh121* mutant is hypersensitive to salt stress. (A) Phenotypes of the *bhlh121-7* mutant and three *bHLH121* overexpression lines grown on ES medium supplemented with 0, 75, 100, or 150 mM NaCl for 10 days. Scale bars, 1 cm. (B) Relative root length. (C) Relative fresh weight. (D, E) Proline contents. (F, G) Malondialdehyde (MDA) contents. (H, I) Electrolyte leakage. (J–M) Na⁺ and K⁺ contents. (N, O) Na⁺/K⁺ ratios. Data are presented as means ± SD; sample sizes are indicated by circles. Different letters indicate significant differences according to one-way ANOVA followed by Tukey’s multiple comparison test (*P* < 0.05); asterisks indicate significant differences from WT according to Student’s t-test (\**P* < 0.05, \*\**P* < 0.01, \*\*\**P* < 0.001, \*\*\*\**P* < 0.0001).

To explore the physiological mechanisms underlying *bhlh121*-associated salt sensitivity, we first quantified proline accumulation, a stress tolerance indicator (Dai et al., 2018), in salt-stressed seedlings. Although proline content increased in both WT and *bhlh121-7* mutant, shoot and root proline levels in *bhlh121-7* were significantly lower than in WT under salt stress (Fig. 1D, E), pointing to a potential mechanism responsible for the mutant’s sensitive phenotype. To assess whether *bHLH121* contributes to antioxidant defense against salts, we measured the malondialdehyde (MDA) content and electrolyte leakage (EL), indicators of oxidative damage (Liu et al., 2021b). MDA induction by NaCl was more pronounced in roots than shoots in both WT and *bhlh121-7*; nevertheless, MDA levels were significantly higher in *bhlh121-7* than WT in both tissues (Fig. 1 F, G). Consistently, EL was triggered by salt stress, with shoot EL higher in *bhlh121-7* than WT under both control and NaCl treatments (Fig. 1H, I).

As ion homeostasis is a major determinant of salt tolerance (Motte et al., 2019; van Zelm et al., 2020) and bHLH121 is crucial for regulating metal ion transporters (Gao et al., 2020; Kim et al., 2019; Lei et al., 2020), we analyzed Na^+^ and K^+^ concentrations and Na^+^/K^+^ ratios in salt-stressed seedlings. In control groups, no obvious differences in basal Na^+^ and K^+^ levels were observed under standard growth conditions in either shoots or roots. By contrast, salt treatment dramatically elevated Na^+^ levels and reduced K^+^ levels in all samples (Fig. 1J–M). Compared with WT, Na^+^ accumulation and K^+^ reduction were more pronounced in shoots, whereas no significant changes were observed in roots. Consequently, Na^+^/K^+^ ratios in shoots, but not in roots, were slightly increased in *bhlh121-7* (Fig. 1N, O).

Collectively, these findings demonstrate that the salt stress sensitivity of *bhlh121-7* roots is due to impaired proline induction and increased MDA contents, whereas significantly increased EL and disrupted Na⁺/K⁺ homeostasis were detected in shoots. Therefore, the *bhlh121-7* mutation appears to impair the plant antioxidant response, especially in roots.

### 3.2 bHLH121 and SOS2 function in parallel pathways in salt tolerance

Although bHLH121 is a key transcription factor regulating Fe homeostasis, its role in plant salt tolerance has not been previously reported in *Arabidopsis*. To investigate the underlying mechanism, we examined the genetic relationship between the bHLH121-mediated salt tolerance pathway and the classical SOS pathway by generating a *bhlh121-7 sos2-T1* double mutant through genetic crossing. Under normal growth conditions, the *bhlh121-7* and *sos2-T1* single mutants and the *bhlh121-7 sos2-T1* double mutant showed no obvious growth defects (Fig. 2A). By contrast, when grown on ES medium supplemented with NaCl at a concentration as low as 25 mM, the *sos2-T1* single mutant displayed a pronounced salt-sensitive phenotype, characterized by significant inhibition of primary root elongation (Fig. 2A), consistent with previous reports (Ma et al., 2025). The *bhlh121-7* single mutant was less salt-sensitive than *sos2-T1* (Fig. 2A). Importantly, the primary root length of the *bhlh121-7 sos2-T1* double mutant was significantly shorter than that of WT and was also substantially shorter than that of either single mutant (Fig. 2A, B). Consistently, the fresh weight and chlorophyll content of the double mutant were lower than those of the two single mutants (Fig. 2C, D). This additive effect suggests that bHLH121 and SOS2 may act in parallel genetic pathways to modulate salt stress responses. To further test this hypothesis, we examined *SOS1* expression, a key downstream gene in the SOS pathway, under salt stress. NaCl treatment induced significantly higher *SOS1* expression in the *bhlh121-7 sos2-T1* double mutant than in WT plants. In contrast, *SOS1* induction in the *bhlh121-7* and *sos2-T1* single mutants was comparable to that in WT (Fig. 2E). These results suggest that the enhanced salt sensitivity of the double mutant is accompanied by altered *SOS1* regulation, consistent with bHLH121 and SOS2 acting in parallel pathways.

**Figure 2.**
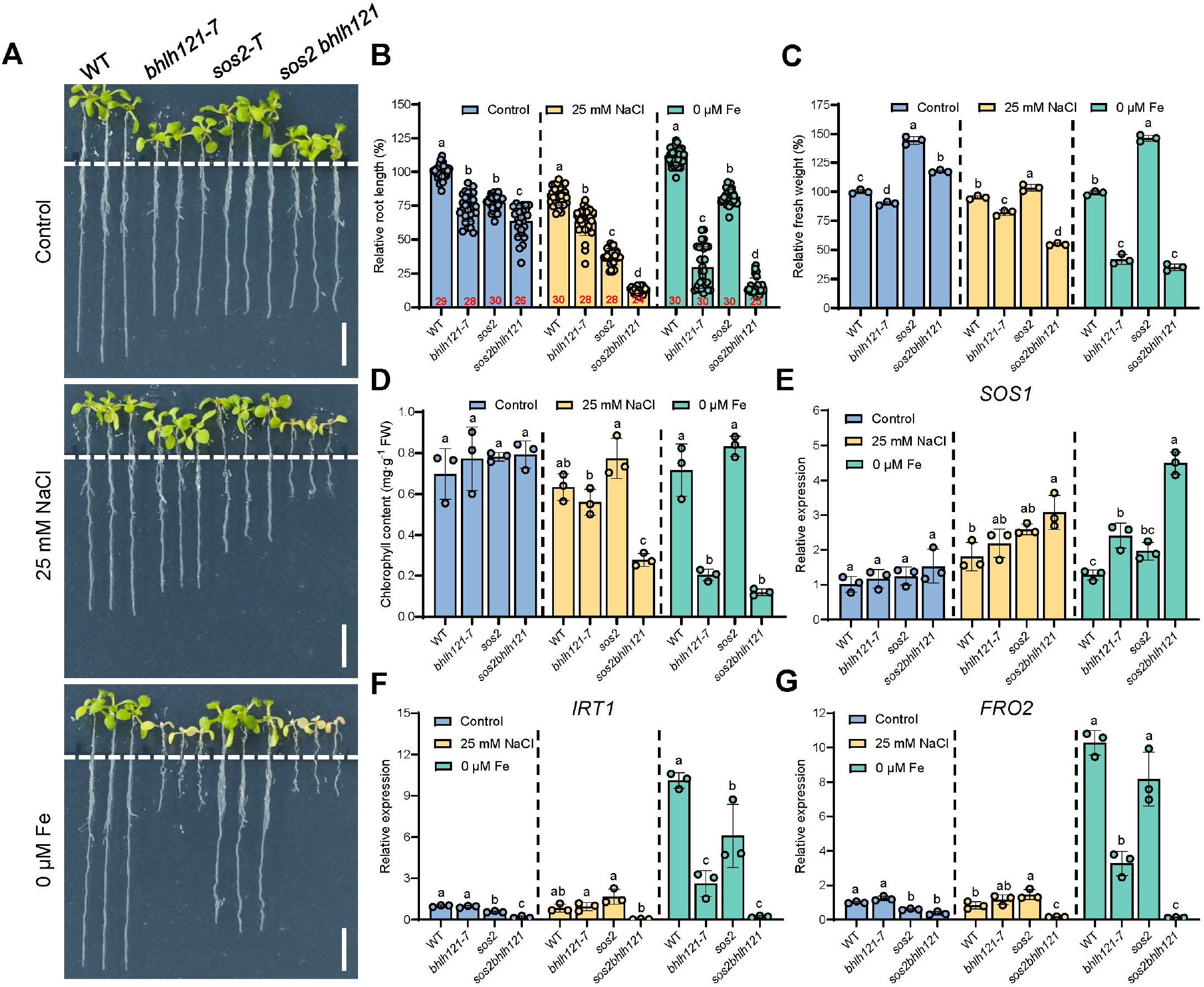
*bHLH121* and *SOS2* act additively in response to salt stress. (A) Phenotypes of WT, the *bhlh121-7* mutant, the *sos2-T1* mutant, and the *bhlh121-7 sos2-T1* double mutant grown on ES medium for 5 d and then transferred to ES medium supplemented with 25 mM NaCl or 0 μM Fe for 7 d. Scale bars, 1 cm. (B) Relative root length. (C) Relative fresh weight. (D) Chlorophyll content. (E–G) Relative expression levels of *SOS1* (E), *IRT1* (F), and *FRO2* (G). Seedlings grown on ES medium for 5 d were transferred to medium supplemented with 25 mM NaCl or 0 μM Fe for 7 d, and root samples were collected for qRT-PCR analysis. Data are presented as means ± SD; sample sizes are indicated by circles. Statistical differences were analyzed by one-way ANOVA followed by Tukey’s multiple comparison test. Different letters indicate significant differences (*P* < 0.05).

We also examined the phenotypes of the *bhlh121-7 sos2-T1* double mutant grown on low-Fe (0 μM Fe) medium. Introduction of the *sos2-T1* mutation into the *bhlh121-7* mutant did not cause obvious growth defects (Fig. 2A). Measurements of root length, fresh weight, and chlorophyll content consistently showed that the *bhlh121-7 sos2-T1* double mutant grew as well as the *bhlh121-7* single mutant (Fig. 2B–D). Notably, *IRT1* and *FRO2*, two marker genes for the Fe deficiency response, showed slightly reduced expression in the double mutant compared with the *bhlh121-7* single mutant (Fig. 2F, G). These data suggest that bHLH121 and SOS2 do not have substantially overlapping roles in the Fe deficiency response.

Taken together, these results suggest that loss of *bHLH121* does not obviously impair SOS pathway-mediated signaling output, consistent with the possibility that bHLH121 functions independently of the SOS pathway to confer salt tolerance.

### 3.3 bHLH121 is crucial for ROS homeostasis and root growth under salt stress

To determine whether ROS accumulation increased abnormally due to bHLH121 dysfunction, we first examined the effects of reduced glutathione (GSH) and potassium iodide (KI), two H_2_O_2_ scavengers, on salt stress-induced root growth inhibition. Although substantial growth inhibition of *bhlh121-7* seedlings was observed with both 500 μM KI and 200 μM GSH in the absence of NaCl (Fig. 3A, B), under salt stress both KI and GSH restored root growth more effectively in *bhlh121-7* than in WT, resulting in root growth recovery with KI (WT, −6.1% vs. *bhlh121-7*, +9.5%) and with GSH (WT, +4.4% vs. *bhlh121-7*, +10.0%) (Fig. 3A, C).

**Figure 3.**
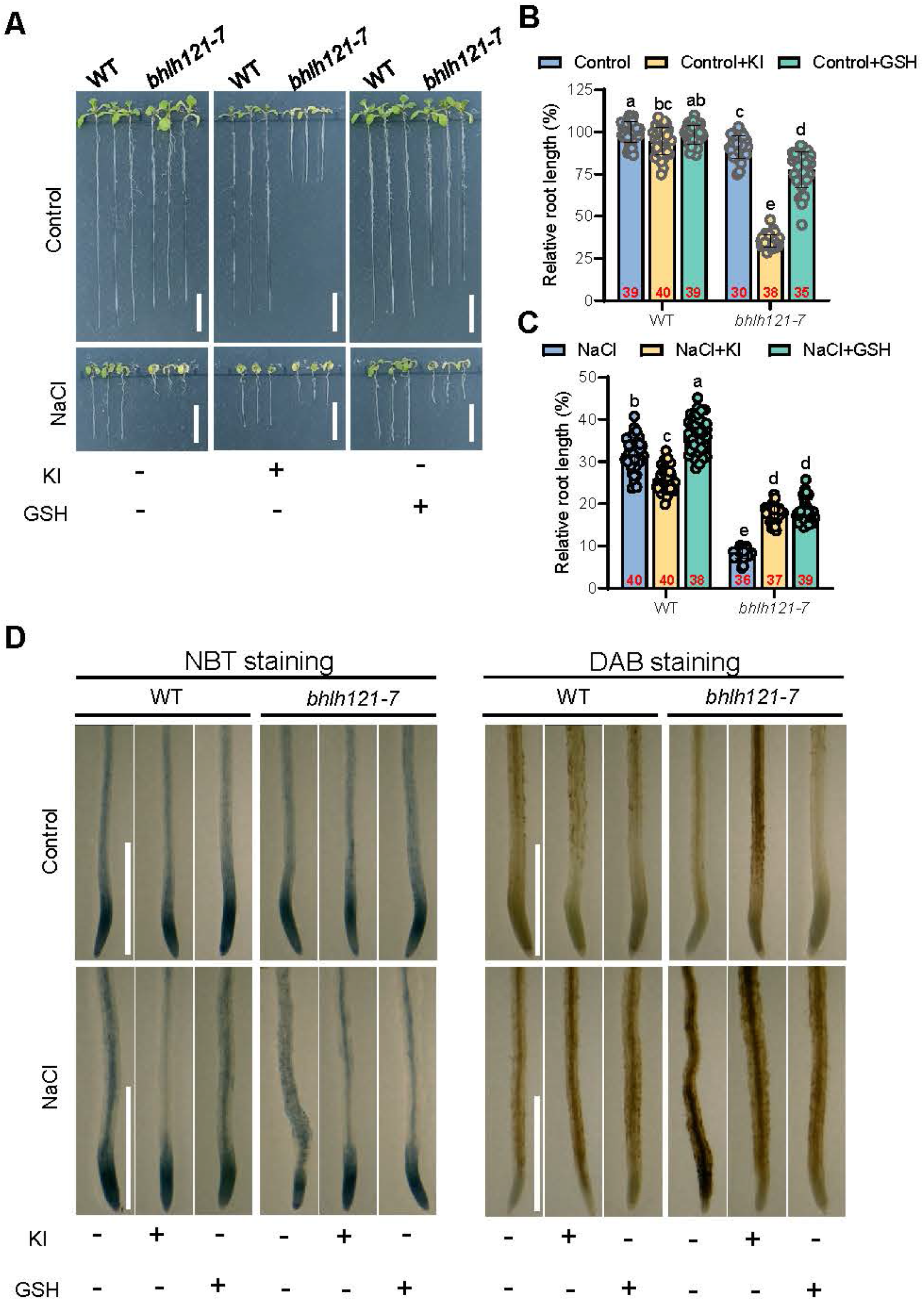
Application of ROS scavengers KI and GSH restores the root growth of the *bhlh121-7* mutant under salt stress. (A) Phenotype of the *bhlh121-7* mutant grown on ES medium supplemented with reactive oxygen species (ROS) scavengers KI (500 μM) or GSH (200 μM) under salt stress for 10 d. Scale bars, 1 cm. (B, C) Relative root length. (D) ROS detection by NBT (O_2_^•−^) and DAB staining (H_2_O_2_). Data are presented as means ± SD; sample sizes are indicated by circles. Statistical differences were analyzed using one-way ANOVA followed by Tukey’s multiple comparisons test. Different letters indicate significant differences (*P* < 0.05).

Since superoxide (O_2_^•**−**^) and H_2_O_2_ are two major ROS involved in root tip growth under salt stress (Dang et al., 2025), their accumulation was detected using nitroblue tetrazolium (NBT) staining for O_2_^•**−**^ (Tsukagoshi et al., 2010) and 3,3′-diaminobenzidine (DAB) staining for H_2_O_2_ (Daudi et al., 2012). In line with this, ROS staining showed reduced H_2_O_2_ accumulation in the presence of KI or GSH under salt stress (Fig. 3D). Notably, O_2_^•**−**^ levels indicated by NBT staining were comparable in *bhlh121-7* and WT under control conditions, whereas slight decreases were observed in the mutant under NaCl treatment (Fig. 3D). Importantly, KI and GSH mitigated salt-induced perturbations in O_2_^•**−**^ levels. Moreover, ROS homeostasis was also restored in bHLH121 complementation lines (Supplementary Fig. S4). These results strongly suggest that bHLH121 is crucial for maintaining ROS homeostasis in roots under high salinity.

ROS spatiotemporal dynamics are vital for plant growth and stress responses. We sought to determine whether ROS imbalance occurs shortly after exposure to high salinity. To this end, we monitored ROS levels at 0, 3, 6, 12, and 24 h after NaCl treatment in the roots of seedlings. Significant differences in O_2_^•**−**^ and H_2_O_2_ signal intensities in root tips were observed after 6 h of salt treatment (Fig. 4). For O_2_^•−^, the two peaks and one trough of NBT signals indicate maximum and minimum O_2_^•−^ levels across the root apical meristem. Salt treatment disrupted the O_2_^•**−**^ distribution pattern in both WT and the mutant (Fig. 4A–C). For DAB staining reflecting H_2_O_2_ distribution, peak signals were predominantly shifted toward the late meristem and elongation zone in control plants (Fig. 4, D–F). Comparable H_2_O_2_ signal intensities were captured in WT, *bhlh121-7*, and bHLH121-complemented lines without NaCl (Fig. 4). In contrast, after NaCl treatment, H_2_O_2_ levels in the *bhlh121-7* mutant were dramatically increased and significantly higher than those in WT and salt-insensitive mutants (Fig. 4D–F). Salt stress also triggered H_2_O_2_ accumulation beyond the meristem and elongation zones (Fig. 4D–F). These results indicate that ROS balance is acutely disrupted in the *bhlh121-7* mutant, with the O_2_^•−^ / H_2_O_2_ ratio significantly decreased compared with WT under salt stress.

**Figure 4.**
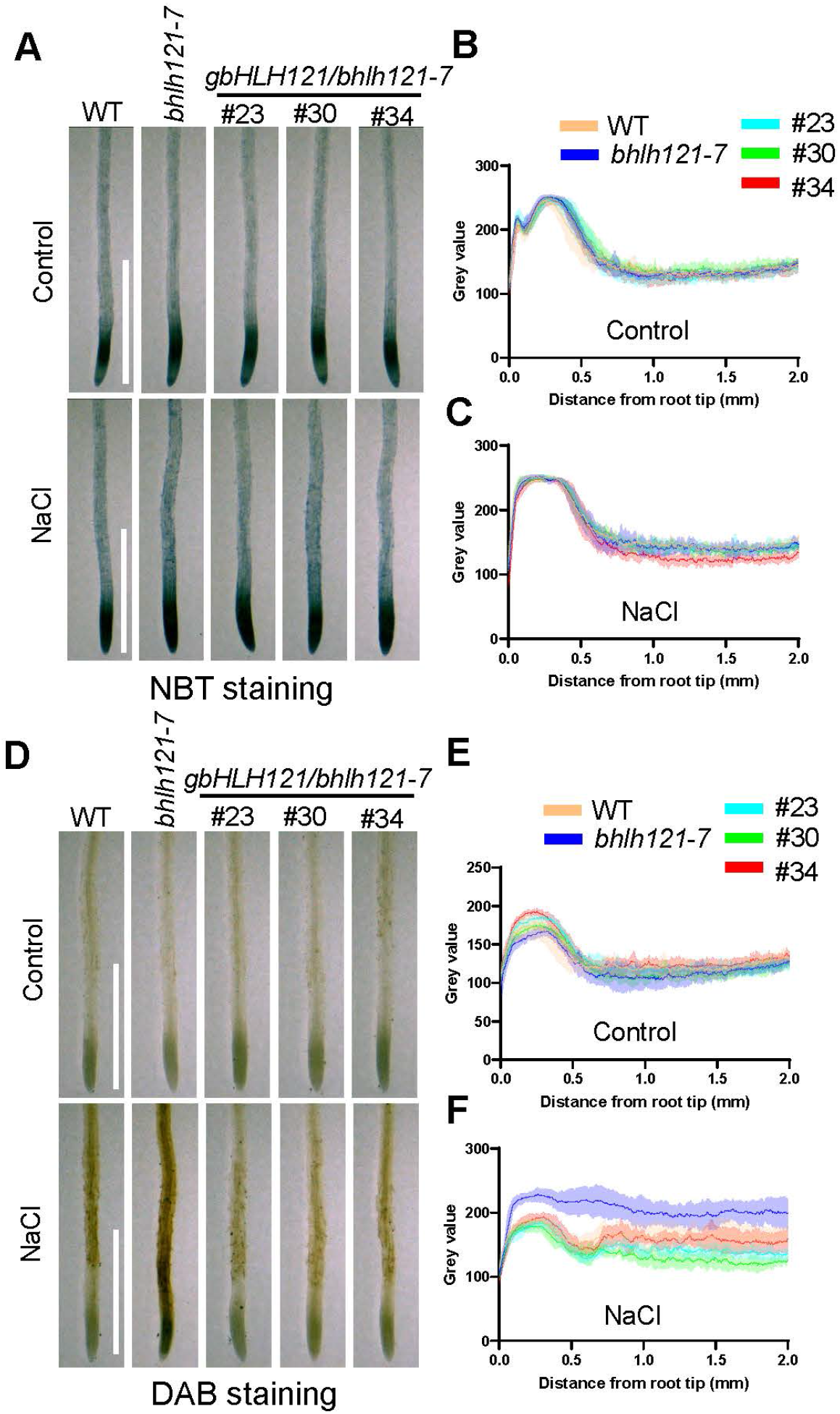
Sharp ROS accumulation in the root tips of the *bhlh121-7* mutant under short-term salt stress. (A–C) Accumulation and quantification of NBT staining (O_2_^•–^) of WT, *bhlh121-7*, and three independent *bHLH121*-complemented transgenic lines. (D–F) Accumulation and quantification of DAB staining (H_2_O_2_) of WT, *bhlh121-7*, and three *bHLH121*-complemented transgenic lines. Seedlings grown on ES medium for 7 d were transferred to liquid ES medium supplemented with or without 100 mM NaCl and treated for 6 h before staining and observation. Staining was quantified using gray values of linear ROIs extending 2 mm along the longitudinal axis from the root tip. Scale bars, 1 mm.

### 3.4 bHLH121 is required for root apical meristem maintenance in salt-stressed roots

Since root apical meristem (RAM) size is a major determinant of root growth rate, we reasoned that the markedly shorter roots of *bhlh121-7* than of WT plants might result from an impaired RAM. To test this possibility, we measured RAM size in salt-treated *bhlh121-7* plants. As expected, RAM size in salt-treated roots decreased by ∼63.9% in *bhlh121-7* versus ∼30.2% in WT (Fig. 5A–D), whereas the number of RAM cells decreased by ∼63.1% in *bhlh121-7* vs. ∼40.2% in WT (Fig. 5A, B, E, F). These results suggest that bHLH121 plays an important role in RAM maintenance under salt stress.

**Figure 5.**
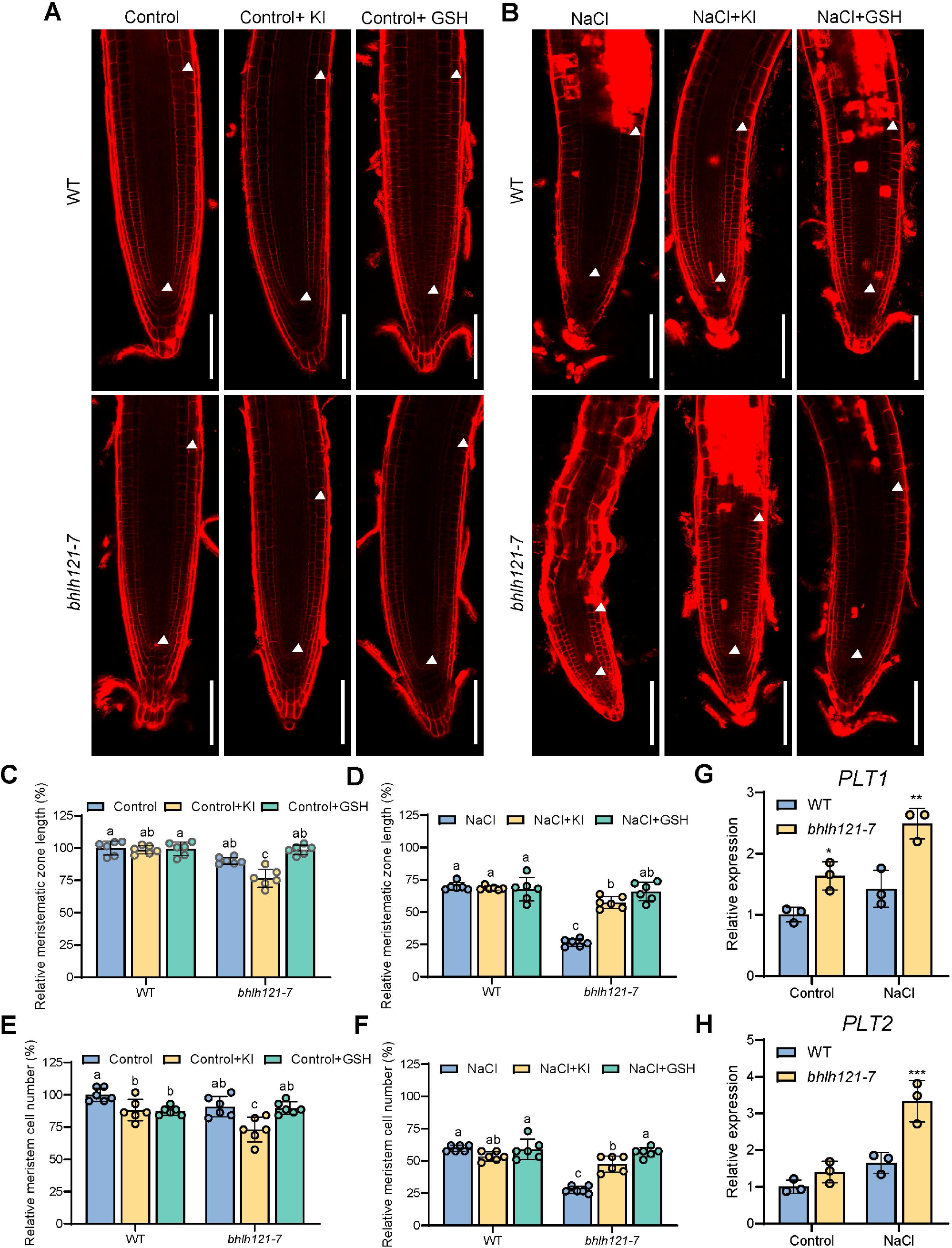
bHLH121 is critical for ROS-dependent regulation of root apical meristem (RAM) size under salt stress. (A, B) Confocal microscopy images of the RAM in WT and the *bhlh121-7* mutant under control conditions (A) and salt stress (B). Roots of 10-day-old seedlings grown on medium with or without the ROS scavengers KI and GSH were stained with propidium iodide (PI) and observed. Scale bars, 100 μm. White triangles indicate the boundaries of the meristematic zone. (C, D) Quantification of RAM length in control (C) and NaCl-treated (D) seedlings. (E, F) Quantification of RAM cell number in control (E) and NaCl-treated (F) seedlings. (G, H) Relative expression levels of *PLT1* (G) and *PLT2* (H). Data are presented as means ± SD; sample sizes are indicated by circles. Different letters indicate significant differences according to one-way ANOVA followed by Tukey’s multiple comparison test (*P* < 0.05); asterisks indicate significant differences from WT according to Student’s t-test (\**P* < 0.05, \*\**P* < 0.01, \*\*\**P* < 0.001).

Given that root apical meristem (RAM) size is determined by the O_2_^•−^ to H_2_O_2_ ratio gradients in the root tip (Dang et al., 2025; Tsukagoshi, 2016), we suspected that the rapid disruption of ROS balance negatively affected RAM size. To test whether modifying oxidative conditions could restore WT-like salt tolerance and RAM size, we treated the *bhlh121-7* mutant with GSH and KI. Consistent with previous results (Fig. 3), both KI and GSH substantially alleviated salt stress-induced root growth inhibition in *bhlh121-7*. Notably, in WT plants, while KI and GSH slightly reduced RAM cell number under control conditions, they had minimal impact on RAM size under salt stress (Fig. 5A, C, E). Nevertheless, both chemicals effectively restored RAM length and cell number, which were severely reduced by high salinity (Fig. 5B, D, F). We also examined the expression levels of *PLETHORA1* (*PLT1*) and *PLT2*, two key genes required for RAM maintenance (Aida et al., 2004). In agreement with our mRNA-seq results, both genes were notably upregulated upon salt stress and displayed significantly higher expression levels in *bhlh121-7* than in WT (Fig. 5G, H), suggesting that bHLH121 dysfunction alters the regulation of RAM maintenance genes.

Altogether, these findings suggest that loss of bHLH121 function results in ROS imbalance in root tips, presumably exacerbating salt-induced root growth inhibition by interfering with meristem maintenance.

### 3.5 Transcriptome analyses support a role for bHLH121 in ROS homeostasis under salt stress

Based on the above results, we speculated that bHLH121 is required for maintaining RAM size to sustain root growth under high salinity. To obtain molecular evidence, we performed mRNA-seq on roots of 10-day-old seedlings grown in ES medium and treated with or without 100 mM NaCl. Under control conditions, 207 differentially expressed genes (DEGs) were identified in *bhlh121-7* roots compared with WT, comprising 157 upregulated and 50 downregulated genes (Supplementary Fig. S5; Supplementary Table S2). Consistent with previously described mRNA-seq results for *uri-1* (Kim et al., 2019), Fe deficiency-responsive genes such as *FRO2* and *IRT1* were remarkably downregulated in *bhlh121-7* (Supplementary Fig. S5C). In contrast, *IMA1/2* and *bHLH100/101* were substantially upregulated (Supplementary Fig. S5C), likely reflecting a compensatory feedback response to the loss of bHLH121, which phenocopies Fe deficiency (Wang et al., 2024).

For salt-responsive genes, 886 (377 upregulated and 509 downregulated) and 1374 (517 upregulated and 857 downregulated) DEGs were identified in WT and *bhlh121-7* roots, respectively (Fig. 6A, B; Supplementary Tables S3 and S4). Gene Ontology (GO) enrichment analyses revealed both overlapping (Supplementary Fig. S6) and specific biological processes in WT and *bhlh121-7* samples (Fig. 6C–F). For example, many DEGs in both WT and *bhlh121-7* roots are involved in stress, water deprivation, hormone responses, hypoxia, cell wall organization or biogenesis, and defense responses (Supplementary Fig. S6). In contrast, cell wall loosening, L-glutamate transmembrane transport, and amino acid transport were the top enriched GO terms specific to WT (Fig. 6C, D). In the *bhlh121-7* mutant, jasmonic acid (JA) response, suberin biosynthetic process, and xylem development were the most significantly enriched biological processes (Fig. 6E, F).

**Figure 6.**
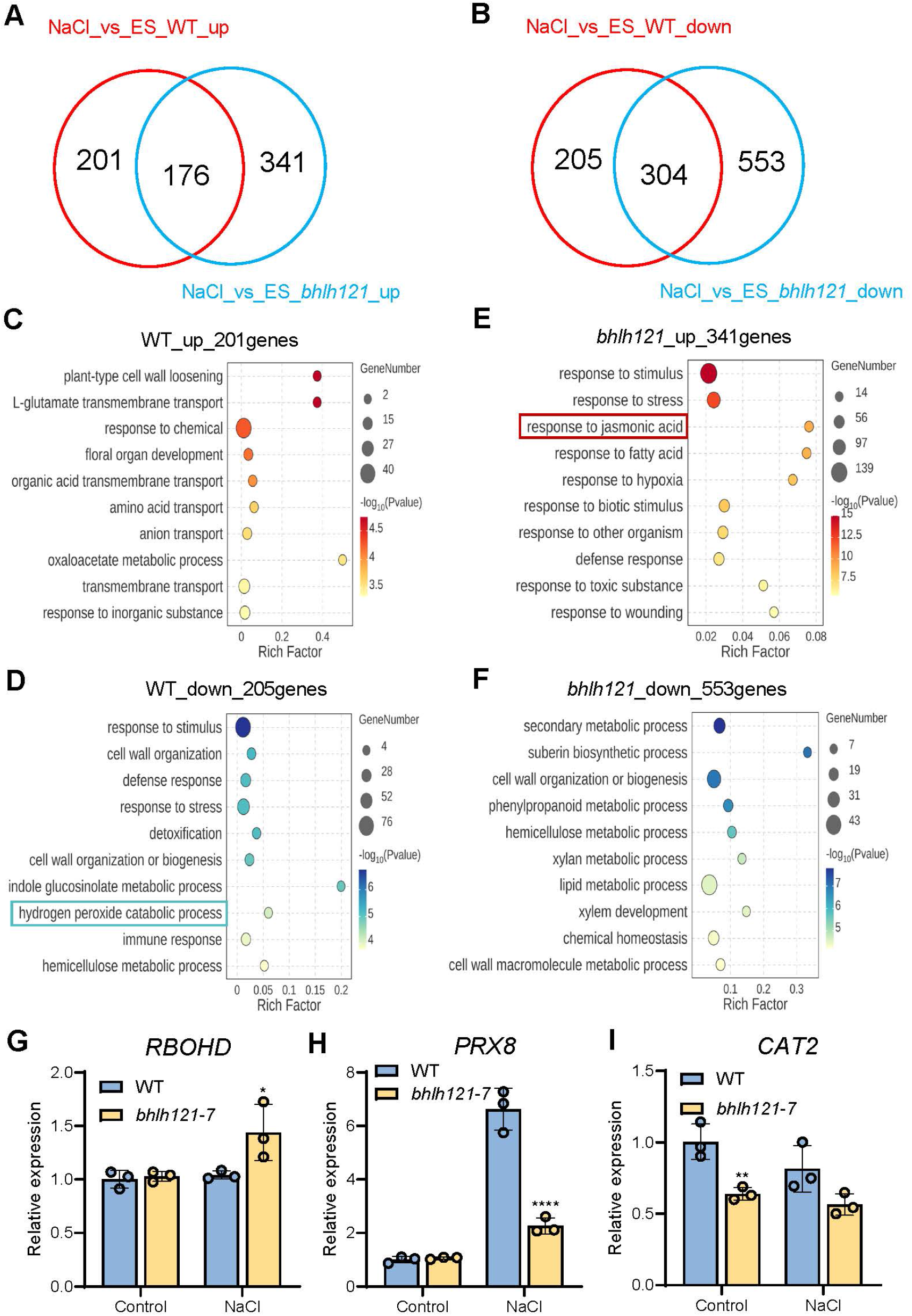
Transcriptome analysis indicates enhanced ROS biosynthesis and impaired H_2_O_2_ scavenging capacity of salt-stressed *bhlh121* plants. (A, B) Venn diagram analysis of salt-upregulated (A) and down-regulated (B) genes between WT and *bhlh121-7*. (C–F) Representative selection of enriched gene ontology (GO) terms among differentially expressed genes (DEGs) that are specifically up- or downregulated upon salt stress in WT (C, D) and *bhlh121-7* mutant (E, F). (G–I) Relative expression of *RBOHD* (G), *PRX8* (H), and *CAT2* (I). Seedlings were germinated and grown for 10 d in control and NaCl (100 mM) agar plates. Root samples were collected and subjected to mRNA-Seq and qRT-PCR. Data are presented as means ± SD (n = 3). Asterisks indicate significant differences from WT according to Student’s t-test (\**P* < 0.05, \*\**P* < 0.01, \*\*\*\**P* < 0.0001).

Notably, DEGs associated with JA response were upregulated in *bhlh121-7*, whereas genes involved in the hydrogen peroxide (H_2_O_2_) catabolic process were downregulated in WT (Fig. 6C–F). Given that ROS homeostasis in *Arabidopsis* roots is tightly regulated during stress responses, these results align with our earlier physiological analyses. We hypothesized that bHLH121 is required to suppress salt stress-induced ROS bursts. This prompted us to investigate which genes in ROS metabolic pathways were significantly changed in *bhlh121-7* under salt stress. We first examined the expression profiles of *RBOH* genes, which encode ROS-producing enzymes that generate superoxide anions (O_2_^•−^) and H_2_O_2_ in response to stress (Mittler et al., 2022). Most *RBOH* genes detected in our mRNA-seq data showed no obvious expression changes, except for *RBOHD* and *RBOHF* (Supplementary Fig. S7A). Moreover, most genes encoding superoxide dismutases (SODs; except FSD1) and peroxiredoxins (PRXs) were upregulated in WT but showed impaired induction (e.g., PRX8) or even downregulation in *bhlh121-7* (Supplementary Fig. S7B).

To further validate their salt-induced responses, we performed qRT-PCR for the ROS metabolic genes *RBOHD*, *PRX8*, and *CAT2* (Fig. 6G–I). *CAT2* was selected because of its important role in H_2_O_2_ scavenging under salt stress (Hu et al., 2026; Liu et al., 2025; Zhuang et al., 2025). In the absence of NaCl, *RBOHD* and *PRX8* showed comparable expression in *bhlh121-7* and WT. After 10 days of salt treatment, *RBOHD* was slightly induced in *bhlh121-7* relative to WT, whereas *PRX8* induction was suppressed (Fig. 6G, H). In contrast, *CAT2* expression was lower in *bhlh121-7* than in WT under control conditions, and NaCl had minimal effect on *CAT2* expression in both genotypes (Fig. 6I). These results are in line with our transcriptome analyses, suggesting that loss of bHLH121 impairs the regulation of key genes involved in ROS metabolism.

Collectively, these results suggest that bHLH121 mediates transcriptional adaptation to salt stress. Specifically, antioxidant responses are upregulated by bHLH121 under high salinity, suggesting an important role for bHLH121 in maintaining ROS homeostasis during salt stress.

### 3.6 Overexpression of *CAT2* restore redox homeostasis and salt sensitivity of *bhlh121* mutant

Since *CAT2* was downregulated in the *bhlh121* mutant (Fig. 6I) and was previously identified as one of the binding targets of bHLH121 in planta by ChIP-seq (Kim et al., 2019), we asked whether bHLH121 modulates ROS homeostasis by regulating *CAT2* expression. To test this hypothesis, we first examined the direct binding of bHLH121 to the *CAT2* promoter using an electrophoretic mobility shift assay (EMSA). As shown in Fig. 7A, a biotin-labeled *CAT2* promoter probe (Biotin-*CAT2*) formed a specific protein–DNA complex with recombinant bHLH121, whereas no such shifted band was detected when the His-TF tag protein was used as a negative control. The specificity of this interaction was further confirmed by competition assays: the addition of excess unlabeled probe (Cold-*CAT2*) effectively diminished the shifted band, whereas the mutated probe with disrupted E-box motifs (Cold-*CAT2-m*) failed to compete for binding, indicating that bHLH121 directly associates with the *CAT2* promoter through the E-box element.

**Figure 7.**
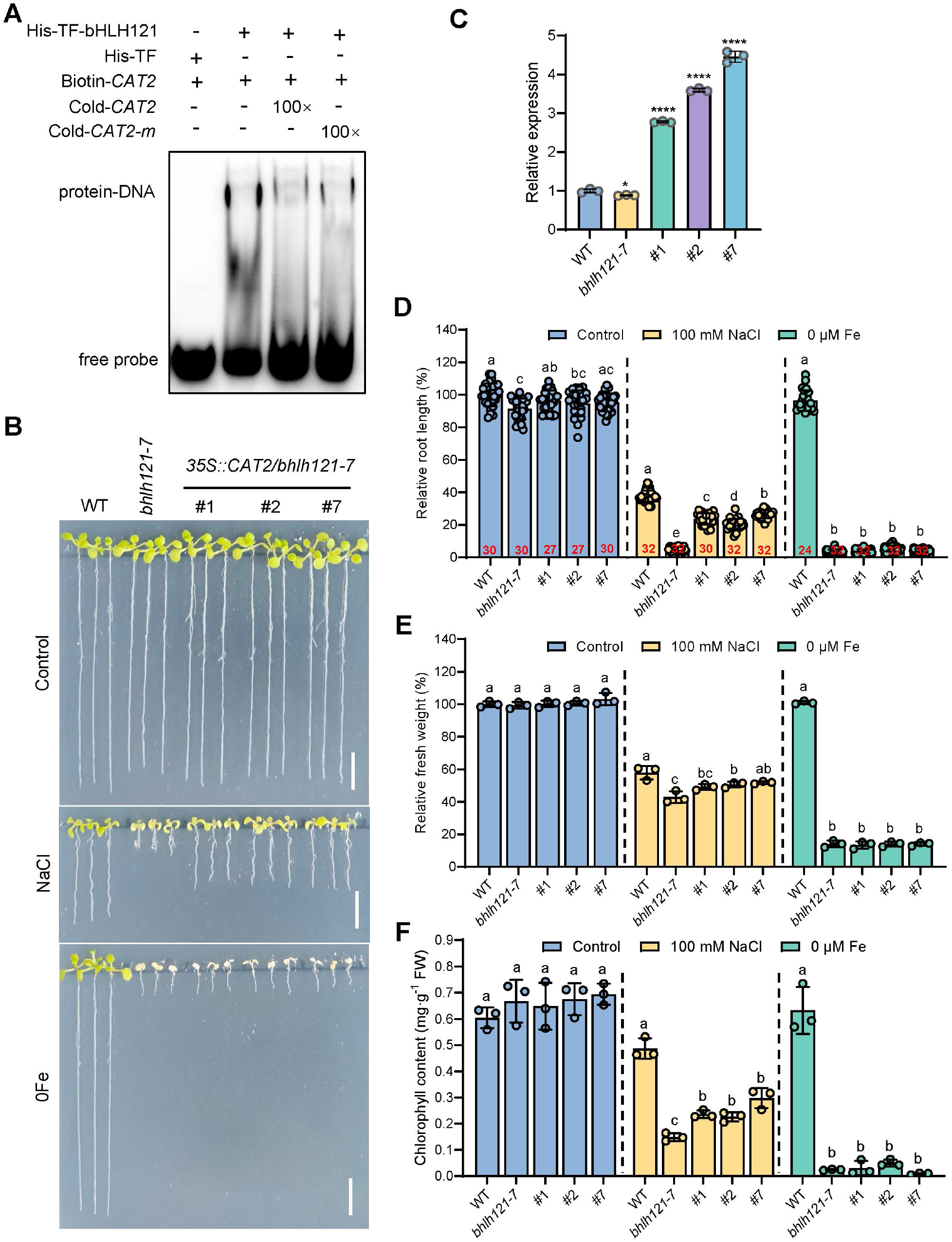
Overexpression of *CAT2* rescues the salt sensitivity of the *bhlh121* mutant. (A) EMSA showing that bHLH121 binds to the promoter of *CAT2*. Biotin-*CAT2*, biotin-labeled probe; Cold-*CAT2*, unlabeled probe; Cold-*CAT2-m*, unlabeled probe with two mutated E-boxes. Biotin-*CAT2* incubated with His-TF served as the negative control. (B) Phenotypes of 35S promoter-driven *CAT2* overexpression transgenic lines in the *bhlh121-7* background grown on ES medium supplemented with 100 mM NaCl or on low-iron medium (0 μM Fe) for 10 d. Scale bars, 1 cm. (C) Relative expression of *CAT2*. Root of 10-d-old seedlings cultured on ES medium were harvested for qRT-PCR. (D) Relative root length. (E) Relative fresh weight. (F) Chlorophyll content. Data are presented as means ± SD; sample sizes are indicated by circles. Statistical differences were analyzed using one-way ANOVA followed by Tukey’s multiple comparisons test. Different letters indicate significant differences (*P* < 0.05). Asterisks indicate significant differences from the WT according to Student’s t-test (\**P* < 0.05, \*\*\*\**P* < 0.0001).

To determine whether *CAT2* acts genetically downstream of bHLH121 in mediating salt tolerance, we generated transgenic lines overexpressing *CAT2* under the control of the 35S promoter in the *bhlh121-7* mutant background (*35S::CAT2*/*bhlh121-7*). When grown on ES medium supplemented with 100 mM NaCl for 10 d, the *35S::CAT2/bhlh121-7* transgenic lines showed significantly improved shoot and root growth and greener leaves relative to the *bhlh121-7* mutant (Fig. 7B). Quantitative reverse transcription PCR (qRT-PCR) analysis confirmed that *CAT2* transcript levels were significantly elevated in the *35S::CAT2/bhlh121-7* lines compared with both WT and the *bhlh121-7* mutant (Fig. 7C), confirming successful overexpression of *CAT2* in the tested transgenic lines. We further quantified physiological parameters to evaluate the extent of phenotypic rescue. The results showed that the *35S::CAT2/bhlh121-7* lines largely restored root length and fresh weight to levels approaching those of WT plants (Fig. 7D, E). In addition, chlorophyll content, which was substantially decreased in the *bhlh121-7* mutant due to stress-induced chlorosis, was significantly increased in the *35S::CAT2/bhlh121-7* plants (Fig. 7F). Notably, we also examined the phenotypes of *35S::CAT2/bhlh121-7* plants grown on ES medium under low-iron conditions (0 μM Fe) for 10 d. Interestingly, unlike the rescue observed under salt stress, the *35S::CAT2/bhlh121-7* plants did not rescue the growth inhibition or leaf chlorosis of the *bhlh121-7* mutant under Fe-deficient conditions (Fig. 7B–F), suggesting that the bHLH121–CAT2 module acts independently of Fe homeostasis.

To further corroborate that *CAT2* overexpression alleviates the salt hypersensitivity of the *bhlh121* mutant by restoring ROS homeostasis, we first measured the H_2_O_2_ content in roots. As expected, salt stress caused an increase in H_2_O_2_ content, but no significant difference was observed among WT, *bhlh121* mutant, and *35S::CAT2/bhlh121* seedling roots (Fig. 8A). We then determined CAT activity in roots. CAT activity increased under salt stress, but differences were observed among lines: it was significantly reduced in the *bhlh121* mutant, whereas overexpression of *CAT2* restored the activity to WT levels in *35S::CAT2/bhlh121* seedling roots (Fig. 8B). By contrast, POD activity also increased under salt stress, but the differences among lines were not significant, suggesting that POD may not be the main contributor to the salt sensitivity of the *bhlh121* mutant (Fig. 8C). Notably, SOD activity decreased under salt stress, possibly due to the prolonged stress duration, and no difference was detected among the tested lines (Fig. 8D). Together, these results support that CAT2 alleviates the salt-sensitive root phenotype of the *bhlh121* mutant by mitigating abnormal H_2_O_2_ accumulation.

**Figure 8.**
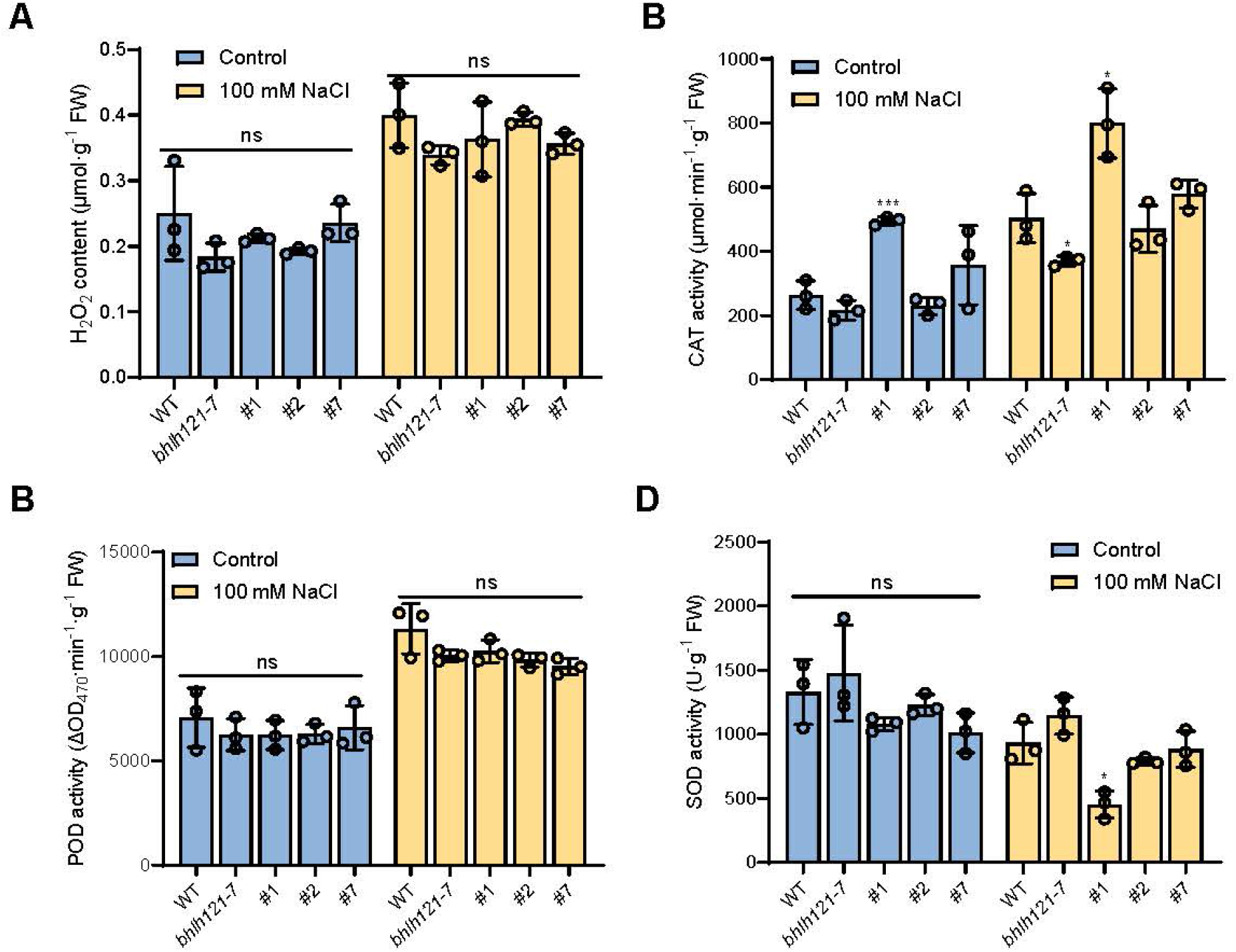
Overexpression of *CAT2* increases the CAT activity of the *bhlh121* mutant. (A) H_2_O_2_ content. (B) CAT activity. (C) POD activity. (D) SOD activity. Seedlings grown on ES medium for 7 d were transferred to salt stress (100 mM NaCl) for 7 d, after which root samples were collected for measurement. Data are presented as means ± SD; sample sizes are indicated by circles. Asterisks indicate significant differences from the WT according to Student’s t-test (\**P* < 0.05, \*\*\**P* < 0.001).

To monitor the distributions of superoxide anion (O_2_^•–^) and hydrogen peroxide (H_2_O_2_) in root tips, NBT and DAB staining were performed, respectively. Consistent with previous results (Figs. 3 and 4), no obvious differences in O_2_^•–^ levels were detected among the tested genotypes under either control or salt conditions (Fig. 9A–C). In contrast, all three tested *35S::CAT2/bhlh121-7* lines displayed substantially reduced staining signals for salt-induced H_2_O_2_, with levels comparable to those of WT plants (Fig. 9D). Quantitative analysis of the gray values along linear regions of interest (ROIs) extending 2 mm from the root tip further confirmed that *CAT2* overexpression effectively attenuated the salt-induced ROS burst in the *bhlh121-7* mutant (Fig. 9E, F). These results provide histochemical evidence that *CAT2* overexpression restores ROS homeostasis in the root tips of the *bhlh121* mutant under salt stress.

**Figure 9.**
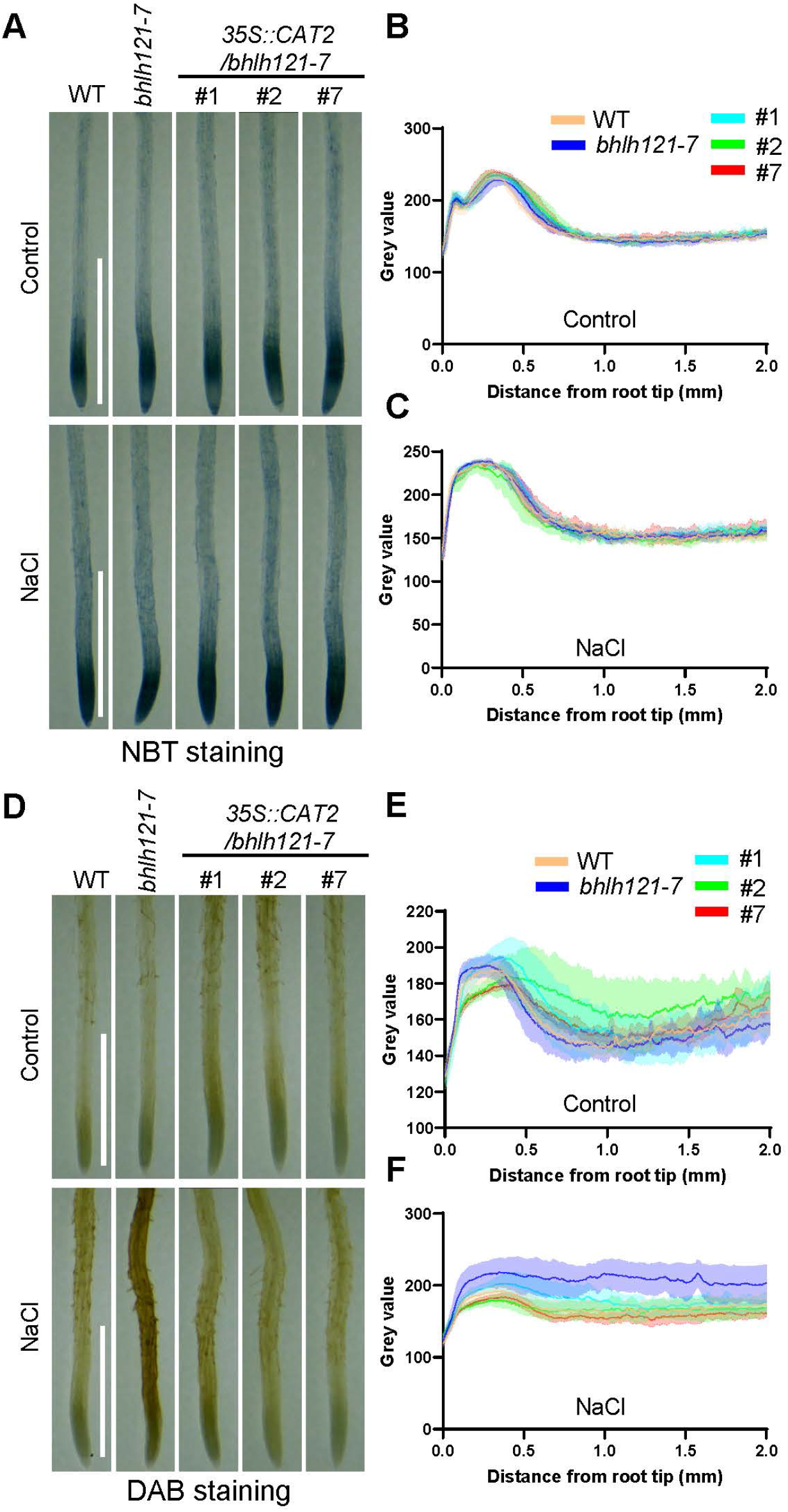
Overexpression of *CAT2* rescues the ROS balance in the root tips of the *bhlh121* mutant under salt stress. (A−C) Accumulation and quantification of NBT staining (O_2_^•−^) in WT, *bhlh121-7*, and three *35S::CAT2*/*bhlh121* transgenic lines. (D−F) Accumulation and quantification of DAB staining (H_2_O_2_) of WT, *bhlh121-7*, and three *35S::CAT2*/*bhlh121* transgenic lines. Seedlings grown on ES medium for 7 d were transferred to liquid ES medium supplemented with or without 100 mM NaCl and treated for 6 h before staining and observation. Staining was quantified using gray values of linear ROIs extending 2 mm along the longitudinal axis from the root tip. Scale bars, 1 mm.

Collectively, these results demonstrate that *CAT2* overexpression potently rescues the salt-sensitive phenotype of the *bhlh121* mutant, supporting the notion that bHLH121 functions upstream of *CAT2* to regulate salt tolerance, at least in part, by modulating ROS homeostasis through CAT2-mediated H_2_O_2_ scavenging.

### 3.7 Auxin and JA pathways are affected in the *bhlh121-7* mutant

ROS homeostasis is tightly associated with hormone signaling in multiple aspects of plant stress responses. Apart from ROS homeostasis genes, comparative transcriptome analysis under salt stress identified a set of upregulated genes specific to *bhlh121-7*. Among these, JA biosynthesis and signaling pathways were significantly enriched (Fig. 6E). Consistently, a number of genes involved in JA biosynthesis, metabolism, and signaling were significantly upregulated in the *bhlh121-7* mutant compared with WT (Supplementary Fig. S8A). Furthermore, we observed a distinct expression pattern of auxin-related genes, most of which were downregulated in the *bhlh121-7* mutant compared with WT (Supplementary Fig. S8B).

To determine whether JA and auxin signaling contribute to the reduced salt tolerance of the *bhlh121-7* mutant, we performed pharmacological tests using the JA biosynthesis inhibitor DIECA (Farmer et al., 1994) and the auxin IAA. For the JA pathway, all tested concentrations of exogenous DIECA (25, 50, and 100 μM) effectively enhanced root growth of the *bhlh121-7* mutant under salt stress, whereas in WT plants it had no effect or was inhibitory at high concentrations (Fig. 10A–C). Similarly, for the auxin pathway, all tested concentrations of exogenous IAA (10, 50, and 100 nM) suppressed root growth in both the *bhlh121-7* mutant and WT under control conditions, whereas they significantly restored root growth in the *bhlh121-7* mutant but not in WT under salt stress (Fig.10D–F). We further examined the expression levels of three key genes involved in JA signaling (*JAZ1* and *JAZ10*) (Liu et al., 2021a) and auxin homeostasis (*GH3.15*) (Sherp et al., 2018) in response to salt in the *bhlh121-7* mutant. Consistent with the transcriptome analyses, both *JAZ1* and *JAZ10* were strongly induced in salt-stressed *bhlh121-7* but not in WT (Fig. 10G, H), suggesting that JA signaling is activated in the *bhlh121-7* mutant. Similarly, *GH3.15* displayed much higher induction in *bhlh121-7* than in WT (Fig. 10I), supporting the notion that bHLH121 functions in maintaining auxin homeostasis.

**Figure 10.**
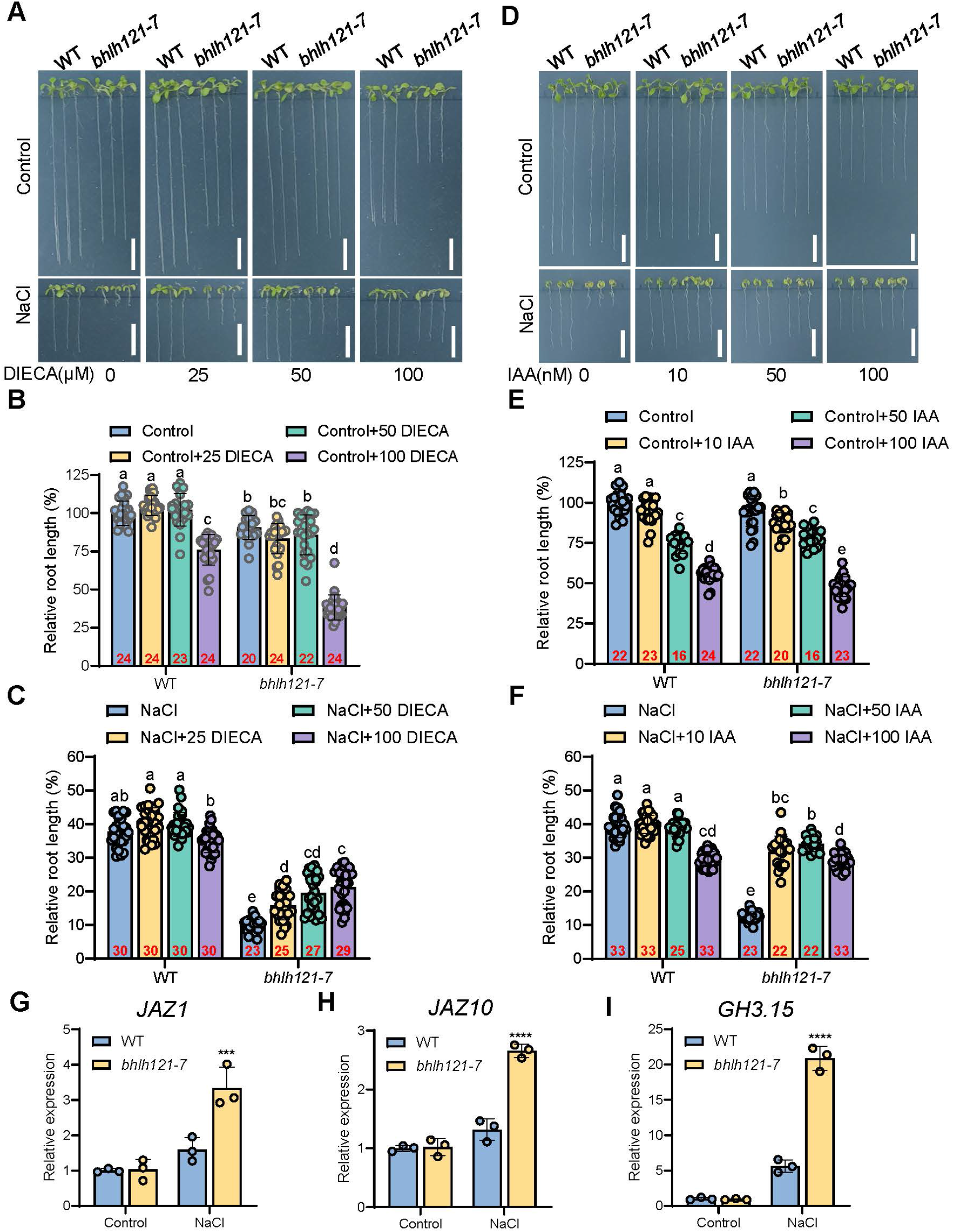
Exogenous application of DIECA and IAA rescues the inhibition of root elongation in the *bhlh121-7* mutant under salt stress. (A−C) Seedlings were grown on medium supplemented with jasmonic acid inhibitor DIECA for 10 d (A). Scale bars, 1 cm. Primary root lengths of seedlings from control (B) and NaCl (C) groups were measured. (D−F) Seedlings were grown on medium supplemented with the auxin IAA for 10 d (D). Scale bars, 1 cm. Primary root lengths of seedlings from the control (E) and NaCl (F) groups were measured. (G-I) Relative expression of *JAZ1*, *JAZ10* and *GH3.15*. Data are presented as means ± SD; sample sizes are indicated by circles. Different letters indicate significant differences according to one-way ANOVA followed by Tukey’s multiple comparisons test (*P* < 0.05); asterisks indicate significant differences from the WT according to Student’s t-test (\*\*\**P* < 0.001, \*\*\*\**P* < 0.0001).

Overall, these findings suggest that bHLH121 is important for modulating both JA signaling and auxin homeostasis under salt stress, thereby contributing to salt tolerance in WT plants.

### 3.8 Conserved function of bHLH121 in salt tolerance across several dicot species

To assess the potential of bHLH121 for crop improvement, we investigated whether its function in salt tolerance is conserved across several plant species. Putative bHLH121 homologs from *Populus trichocarpa*, *Glycine soja*, *Cucumis sativus*, and *Citrus clementina* were previously expressed in the *bhlh121-7* mutant and showed conserved function in the Fe deficiency response (Zhou et al., 2026). Using the same transgenic plants, we tested whether overexpression of these bHLH121 homologous genes rescued the salt hypersensitivity of the *bhlh121-7* mutant. Indeed, all four genes substantially rescued the salt-sensitive phenotypes in both roots (Fig. 11) and shoots (Supplementary Fig. S9). Consistent with the effect of *bHLH121* overexpression in WT (Fig. 1A), *bHLH121* overexpression in the *bhlh121-7* mutant restored root length and seedling fresh weight to WT-like levels but did not enhance salt tolerance beyond WT levels (Fig. 11). Altogether, these results suggest that the role of bHLH121 in conferring salt tolerance is conserved in several dicot species, highlighting the potential of using bHLH121 homologs to improve salt tolerance across a broad range of dicot crops.

**Figure 11.**
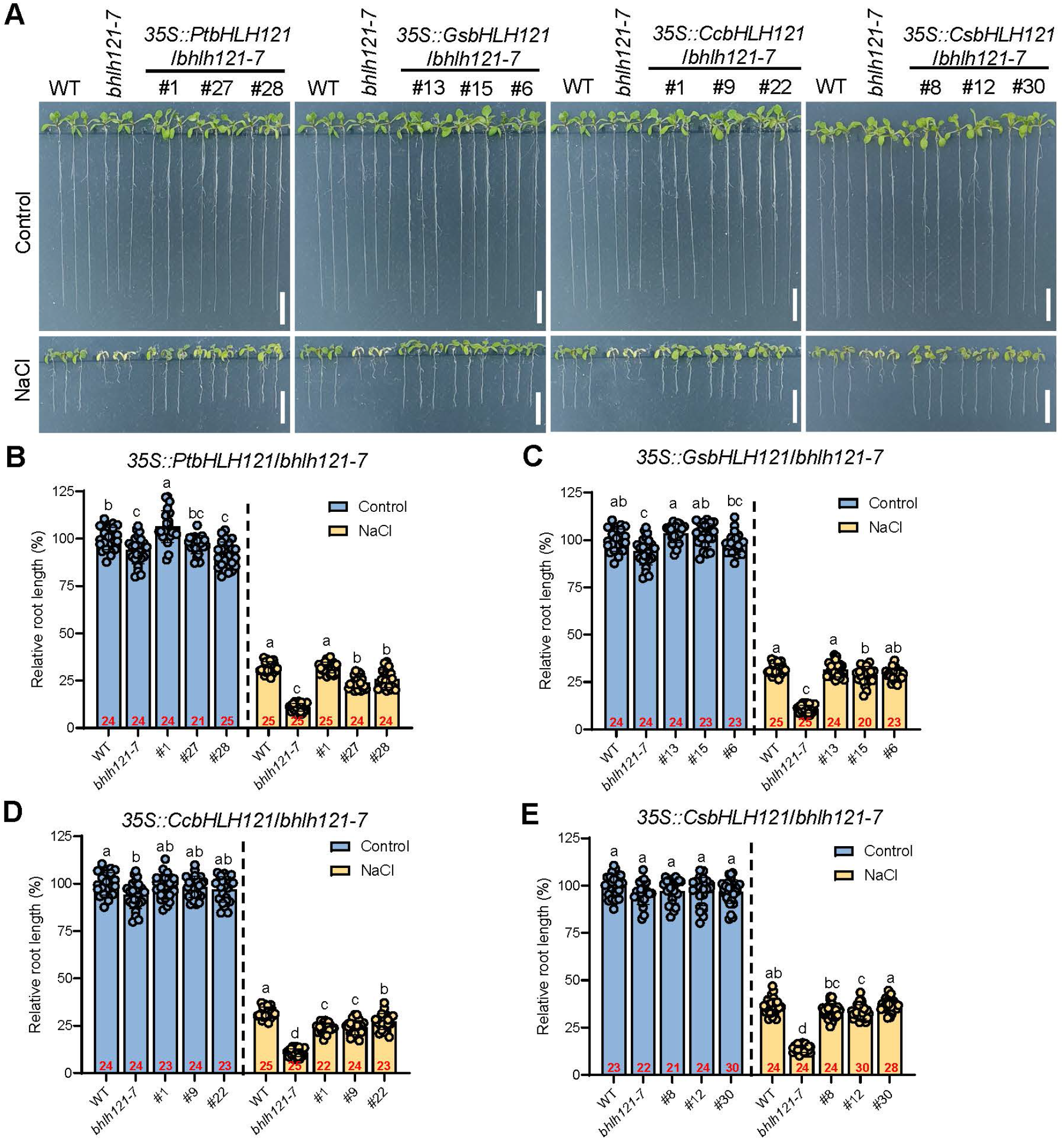
Overexpression of *bHLH121* homologous genes rescues the salt sensitivity of the *bhlh121* mutant. (A) Phenotypes of 35S promoter-driven *Populus PtbHLH121*, soybean *GsbHLH121*, Citrus *CcbHLH121*, and Cucumber *CsbHLH121* overexpression transgenic lines in the *bhlh121-7* background grown on ES medium supplemented with 100 mM NaCl for 10 d. Scale bars, 1 cm. (B–E) Relative root lengths. Data are presented as means ± SD; sample sizes are indicated by circles. Statistical differences were analyzed using one-way ANOVA followed by Tukey’s multiple comparisons test. Different letters indicate significant differences (*P* < 0.05).

## 4 Discussion

The escalating challenge of soil salinity, intensified by global climate change, poses a significant threat to agricultural productivity and food security worldwide. Salinity severely constrains plant growth and yield, and decades of research have sought to unravel the molecular underpinnings of salt stress responses. Seminal work, such as the identification of the Salt Overly Sensitive (SOS) signaling pathway, provided foundational insights into ionic homeostasis (Yang et al., 2018; Zhu, 2002). Despite these advances, the genetic arsenal available for engineering salt-tolerant crops remains limited, underscoring the need to identify novel regulatory components. Intriguingly, recent findings demonstrate that Fe homeostasis components can coordinately regulate both Fe nutrition and salt tolerance across diverse plant species (Li et al., 2017; Liang et al., 2022; Zhang et al., 2025), suggesting an intrinsic link between the Fe homeostasis regulatory network and salt stress responses. However, the mechanistic link coordinating Fe homeostasis and salt adaptation has remained elusive. Our study identifies the transcription factor bHLH121 as a previously unrecognized central player in salt tolerance. The *bhlh121* mutant exhibited profound hypersensitivity. This distinct phenotype positions bHLH121 not merely as a conduit for Fe-deficiency signals but as a potential integrator of Fe homeostasis and salt stress adaptation. Further transcriptomic and biochemical evidence indicates that the bHLH121–CAT2 module exerts its essential function by modulating the redox homeostasis within the root apex, thereby preserving meristem activity and root growth under saline conditions (Supplementary Fig. S10). These findings provide insight into how a core Fe signaling component is co-opted to mediate oxidative stress management, offering a fresh perspective on the interplay between nutrient sensing and abiotic stress resilience.

Recent studies suggest that peanut bHLH121 functions beyond Fe homeostasis by targeting ROS-related genes such as *SOD*, *POD*, and *CAT* (Zhao et al., 2024), which are well-documented regulators of plant growth and stress responses. The present work establishes a specific role for the bHLH121–CAT2 module in salt tolerance in *Arabidopsis*. Although Fe homeostasis is tightly associated with ROS dynamics, our results indicate that *CAT2* overexpression rescues salt hypersensitivity but not Fe deficiency-related growth defects (Fig. 7), strongly suggesting an Fe homeostasis–independent axis regulated by the bHLH121–CAT2 module. Direct binding of bHLH121 to the *CAT2* promoter was previously observed by ChIP-seq (Kim et al., 2019), and we confirmed this direct binding in vitro (Fig. 7A). Nevertheless, whether additional factors are required for *CAT2* expression remains to be investigated. Given that bHLH121 interacts and cooperates with IVc bHLH transcription factors to regulate downstream targets in the Fe regulatory network, likely because it lacks an activation or repression domain (Lei et al., 2020), we hypothesize that bHLH121 may similarly function as part of a transcriptional complex responsible for salt stress resilience.

The WT-like salt-tolerant phenotypes of complementation and overexpression transgenic lines, regardless of *bHLH121* expression levels, suggest that bHLH121 may undergo post-transcriptional modification for its function in salt tolerance. In fact, phosphorylation of bHLH121 is crucial for the Fe deficiency response in *Arabidopsis* (Kim et al., 2019), raising the intriguing question of whether phosphorylation also contributes to its role in regulating plant salt tolerance. Moreover, SOS2 from the SOS pathway is a well-documented salt stress-activated kinase that phosphorylates a subset of target proteins associated with ion nutrition (Ma et al., 2025). Although our results suggest that bHLH121 may act in parallel with SOS2 under salt stress, potential interactions between bHLH121 downstream targets, such as transporters, and SOS2 cannot be excluded.

Our results further indicate the involvement of JA and auxin signaling components in bHLH121-mediated salt tolerance. The salt hypersensitivity of *bhlh121* mutant roots may arise from a detrimental interplay between disrupted ROS homeostasis and hormone signaling. Overproduction of ROS, driven by *RBOHD* overexpression, coupled with impaired scavenging capacity reflected by low *CAT2* and *PRX8* expression (Fig. 6), creates severe oxidative stress. This directly damages root cells and is further amplified by hyperactivation of the JA pathway, as evidenced by feedback upregulation of *JAZ1* and *JAZ10* (Fig. 10). Concurrently, the *bhlh121-7* mutant initiates a maladaptive stress response by drastically suppressing auxin levels through induction of the auxin-inactivating enzyme *GH3.15* (Sherp et al., 2018). This represents an extreme growth–defense trade-off, in which resources are diverted from growth at an excessive cost. The resulting combination of rampant oxidative damage and near-complete shutdown of growth-promoting auxin signaling arrests root development. Phenotypic rescue achieved by inhibiting JA synthesis with DIECA or by supplying exogenous auxin confirms this mechanistic model: DIECA dampens the JA–ROS feedback loop, alleviating oxidative stress, while external IAA bypasses the internal auxin blockade to reactivate growth. Thus, the *bhlh121* mutant phenotype emerges from a pathologically reinforced network in which ROS, JA, and auxin signals synergize to lock the plant in an over-defensive state, ultimately compromising fitness under salt stress (Supplementary Fig. S10). The precise molecular mechanism(s) by which bHLH121 coordinates JA and auxin signaling warrant further investigation.

Catalases play a pivotal role in plant detoxification by mitigating ROS accumulation under abiotic stress (Li et al., 2015; Mhamdi et al., 2012; Zhang et al., 2021). These enzymes catalyze the dismutation of two H_2_O_2_ molecules into H_2_O and molecular oxygen (Frugoli et al., 1996). The Arabidopsis genome encodes three catalase isoforms, CAT1, CAT2, and CAT3, each consisting of 492 amino acids with high sequence similarity (Frugoli et al., 1996; Mhamdi et al., 2010). Among these, CAT2 is the predominant isoform, accounting for approximately 80% of total catalase activity, whereas loss of CAT3 causes only minor reductions and CAT1 deletion has negligible effects (Du et al., 2008; Mhamdi et al., 2010; Zou et al., 2015). Consistent with this hierarchy, the *cat1 cat2 cat3* triple mutant displays severe redox homeostasis disruption and hypersensitivity to salt stress (Su et al., 2018). Further supporting the conserved role of catalases in stress adaptation, silencing of *SmCAT4* in eggplant (*Solanum melongena*) compromises salt tolerance, concomitant with elevated H_2_O_2_ accumulation and diminished catalase activity (Shen et al., 2023). Moreover, jasmonic acid modulates salt tolerance through MYC2-mediated transcriptional repression of *CAT2* (Song et al., 2021), while the CRCK3–CAT2 module contributes to salt stress resilience by maintaining cellular redox homeostasis (Zhuang et al., 2025). Recent findings reveal an additional layer of CAT2 regulation: a redox-regulated RCC1-like protein, CAIR1, controls catalase activity by facilitating its peroxisomal import and preventing its aggregation (Hu et al., 2026). These findings underscore the essential function of catalases in plant adaptive responses to environmental stresses and provide a mechanistic context for the bHLH121–CAT2 axis described here.

ROS dynamics in plants constitute a crucial signaling hub that governs growth, development, and stress adaptation (Castro et al., 2021; Mittler et al., 2022; Tsukagoshi, 2016). The production of ROS via the Fenton reaction (Halliwell et al., 1992) highlights an intrinsic link between ROS dynamics and Fe homeostasis; however, the precise interplay between Fe nutrition and the ROS burst triggered by salt stress remains poorly understood. Though the present study suggests that bHLH121–CAT2 module acts independently of Fe homeostasis regulatory genes (Fig. 7), we cannot exclude the possibility that bHLH121 regulates salt tolerance in an Fe-dependent manner. It is important to emphasize that Fe homeostasis encompasses multiple dynamic processes, including uptake, transport, distribution, storage, and allocation, and is systemically regulated (Liang, 2022; Wang et al., 2024). Based on the present data, we propose that bHLH121 specifically mediates the integration of systemic Fe signaling with stress responses, including ROS and hormone signaling. A recent study highlighted the role of heat-induced nutritional stress under prolonged high-temperature conditions and suggested that Fe homeostasis represents a promising target for enhancing crop resilience to climate extremes (Minami et al., 2025). Whether bHLH121 also functions in plant responses to environmental stresses beyond salinity constitutes an important direction for future research.

In response to stress, plant roots, as the primary interface sensing the external environment, rapidly curtail growth to reallocate resources—a strategic adaptation that hinges on precise regulation of meristematic activity (Liu et al., 2015; Motte et al., 2019). Salt-induced ROS bursts in the root tip are harmful and cause growth arrest and cellular dysfunction (Dang et al., 2025; Xie et al., 2011). Critically, the size of the root apical meristem is determined by the spatial gradient of the O_2_^•**−**^ to H_2_O_2_ ratio (Tsukagoshi, 2016; Tsukagoshi et al., 2010). Therefore, ROS homeostasis in the root must be tightly controlled during plant stress responses. The spatiotemporal dynamics of ROS are intricately governed by numerous nonenzymatic antioxidants and detoxifying enzymes regulated by hormone signaling, Ca^2+^ fluxes, and MAPK cascades (Castro et al., 2021). However, the identity of upstream regulators that fine-tune this localized redox landscape has remained elusive. Our identification of bHLH121 as a regulator of *CAT2* expression and ROS homeostasis in the root apex provides a candidate for such an upstream regulator and offers a framework for understanding how Fe signaling components are integrated into salt stress responses.

In agricultural practice, managing micronutrients such as Fe to enhance plant tolerance and secure crop yield in saline soils remains challenging (Kong et al., 2022; Yan et al., 2023). H_2_O_2_, for instance, has been proposed as a potential crop growth regulator (Iqbal et al., 2025). Our study provides mechanistic insights that may underlie the additive effects of combining such strategies to combat high salinity. Further investigation is needed to determine whether stress-induced ROS patterns or the artificial application of ROS reciprocally influence Fe homeostasis. In conclusion, this study not only advances our understanding of the physiological and molecular mechanisms underlying bHLH121-mediated salt tolerance but also opens avenues for bHLH121-based molecular strategies to enhance crop resilience against salinity stress, a challenge likely to be intensified by global climate change.

## Author contributions

P.L., C.L., and R.W. conceived and designed the experiments; P.Z., Y.P., R.W., Y.F., B.L., and X.D. performed the experiments; P.L. and C.L. performed the mRNA-seq analysis; C.L. and R.W. helped with the data analysis; C.L., P.Z., R.S., Y.C, M.I.H. and P.L. wrote and revised the article; P.L., C.L., L.L., R.S., and Y.C. provided funding. All authors contributed to the review and editing of the manuscript.

## Acknowledgements

We thank Prof. Yan Guo (China Agricultural University) for kindly sharing *sos2-T1* seeds. This work was supported by the National Natural Science Foundation of China (No. 32300274, No. 32070279), National Key R&D Program of China (No. 2023YFD1702205), Project of Priority and Key Areas, Institute of Soil Science, Chinese Academy of Sciences (No. ISSASIP2222, No. ISSASIP2206), and Natural Science Foundation of Jiangsu Province (No. BK20221560).

## Supplementary data

**Supplementary Fig. 1.**
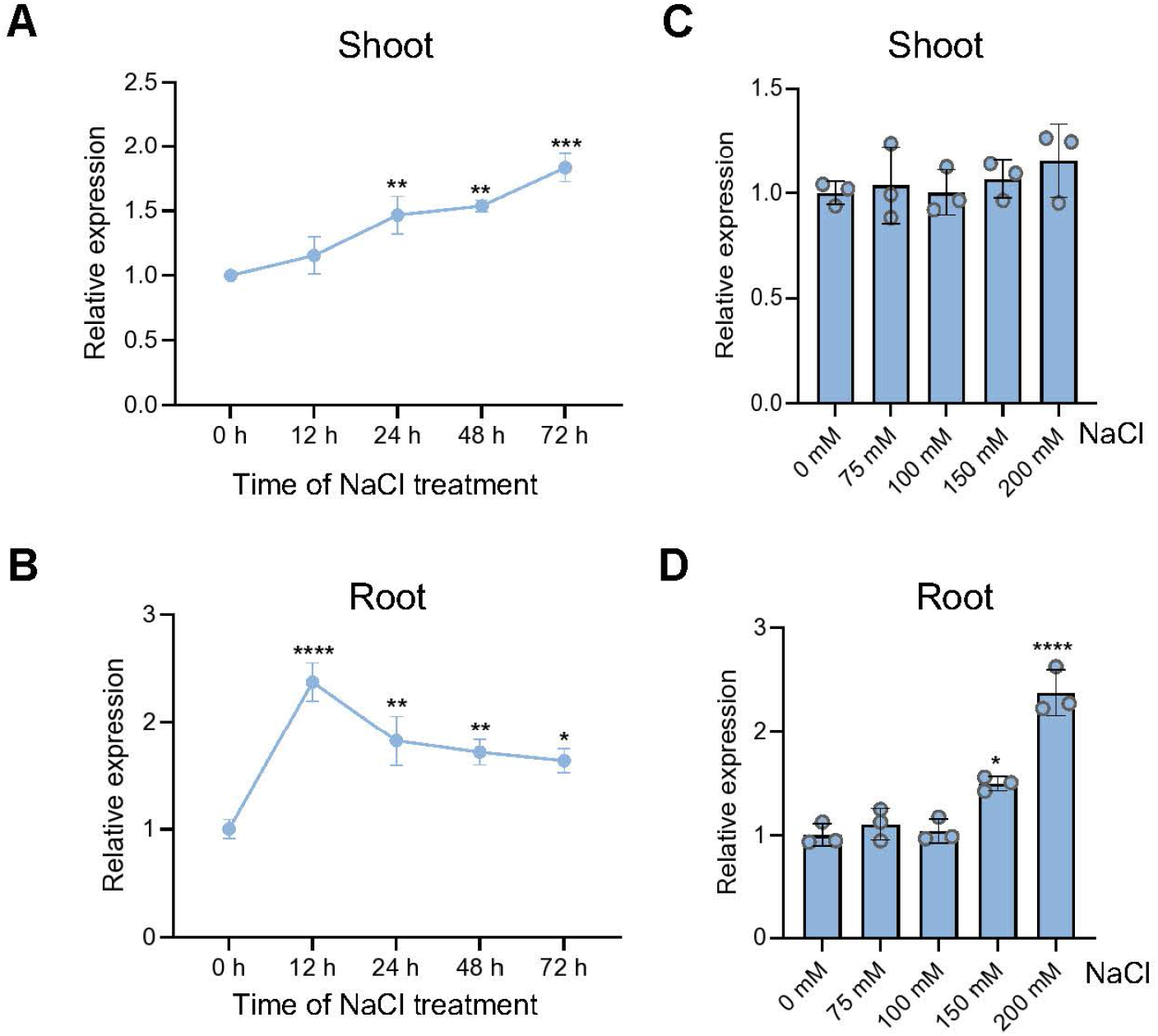
Gene expression analysis of *bHLH121* in Arabidopsis seedlings in response to salt stress. Relative gene expression of *bHLH121* was determined by RT-qPCR. (A, B) Arabidopsis Col-0 seedlings grown on ES medium for 10 d were transferred to ES medium with 200 mM NaCl for treatment. Shoots (A) and roots (B) samples were collected separately at indicated time points. (C, D) 10-day-old Col-0 seedlings were transferred to ES medium supplemented with varying concentrations of NaCl (75 mM, 100 mM, 150 mM, 200 mM) for treatment. After 12 h, shoots (C) and roots (D) samples were collected separately. *Tubulin α-3* was used as an internal reference control, and transcript levels were calculated using the 2^-ΔΔCT^ method. Data are means ± sd (n = 3 technical repeats). Statistical significance was determined according to Student’s t-test (\**P* < 0.05, \*\**P* < 0.01, \*\*\**P* < 0.001, \*\*\*\**P* < 0.0001).

**Supplementary Fig. 2.**
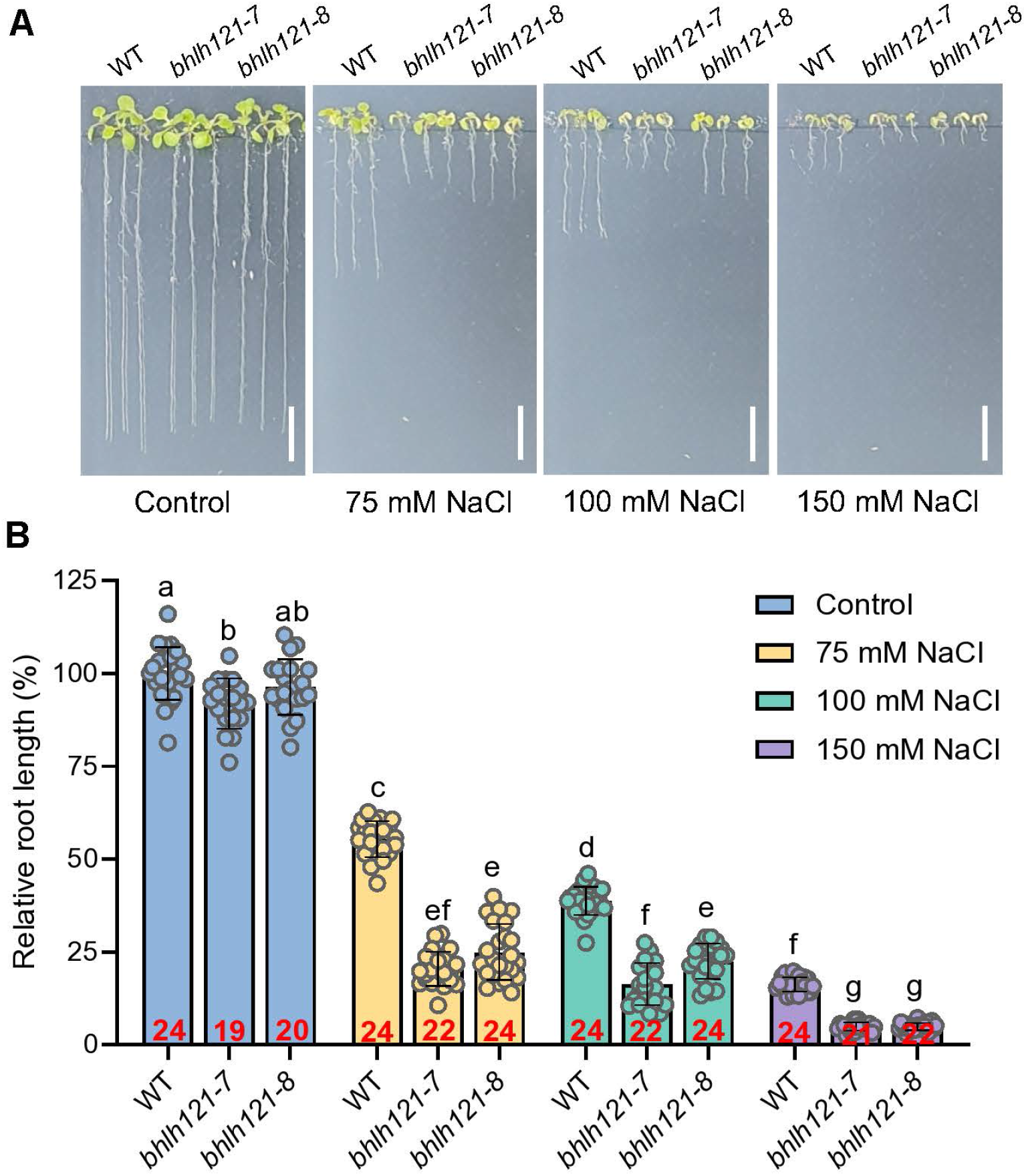
Both *bhlh121-7* and *bhlh121-8* mutants show hypersensitivity to salt stress. (A) Phenotypes of WT, *bhlh121-7*, and *bhlh121-8* plants grown on ES mediums supplemented with 0, 75, 100, 150 mM NaCl for 10 d. Scale = 1 cm. (B) Relative root length of WT, *bhlh121-7*, and *bhlh121-8* mutants under control or salt stress conditions. Data are presented as means ± sd (sample size is indicated by circles). The statistical differences were analyzed using a one-way ANOVA followed by Tukey’s multiple comparisons test. Different letters indicate significant differences (*P* < 0.05).

**Supplementary Fig. 3.**
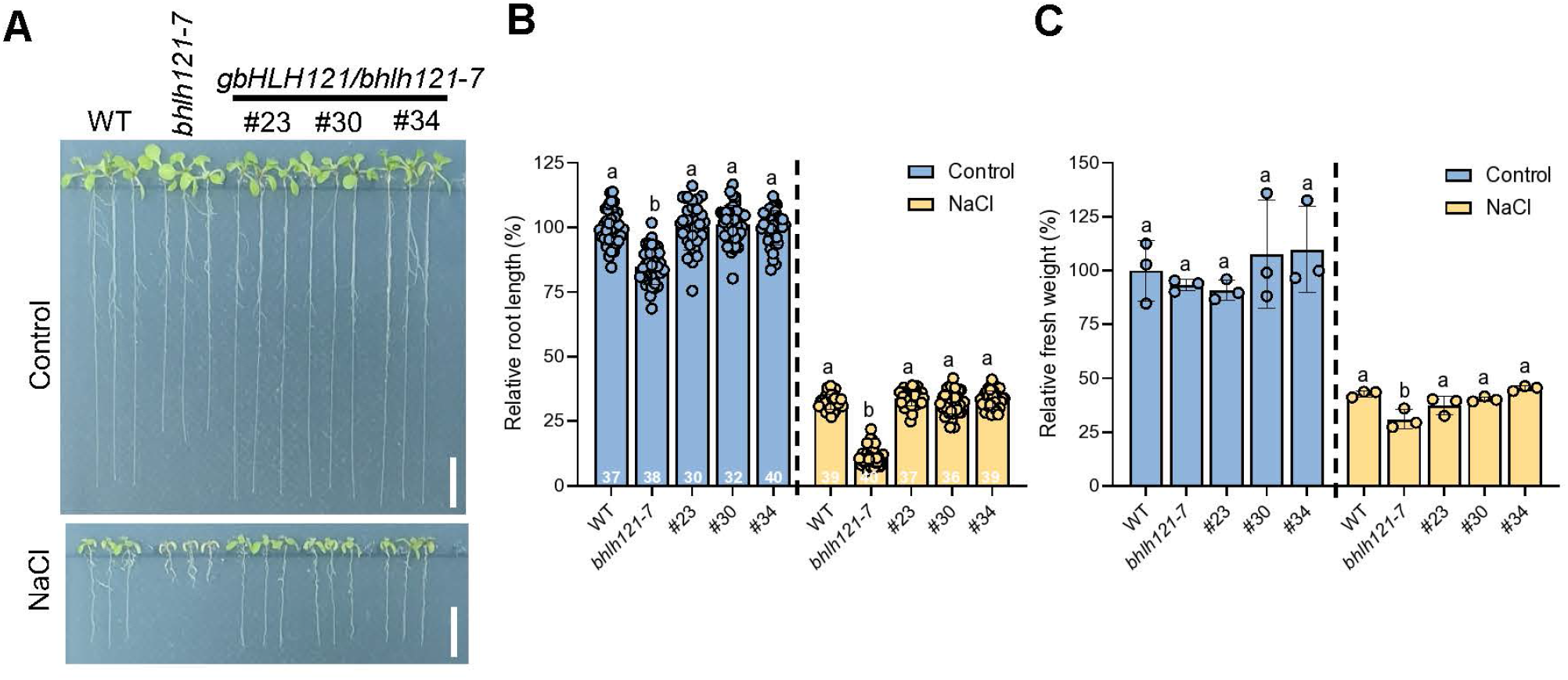
The full-length *bHLH121* gene rescues the salt hypersensitivity of the *bhlh121* mutant. (A) Phenotypes of the *bHLH121*-complementary transgenic lines grown on ES medium supplemented with 100 mM NaCl for 10 days. Scale = 1 cm. (B) Relative root length. (C) Relative fresh weight. Data are presented as means ± sd (sample size is indicated by circles). Different letters indicate significant differences for a one-way ANOVA followed by Tukey’s multiple comparisons test (*P* < 0.05).

**Supplementary Fig. 4.**
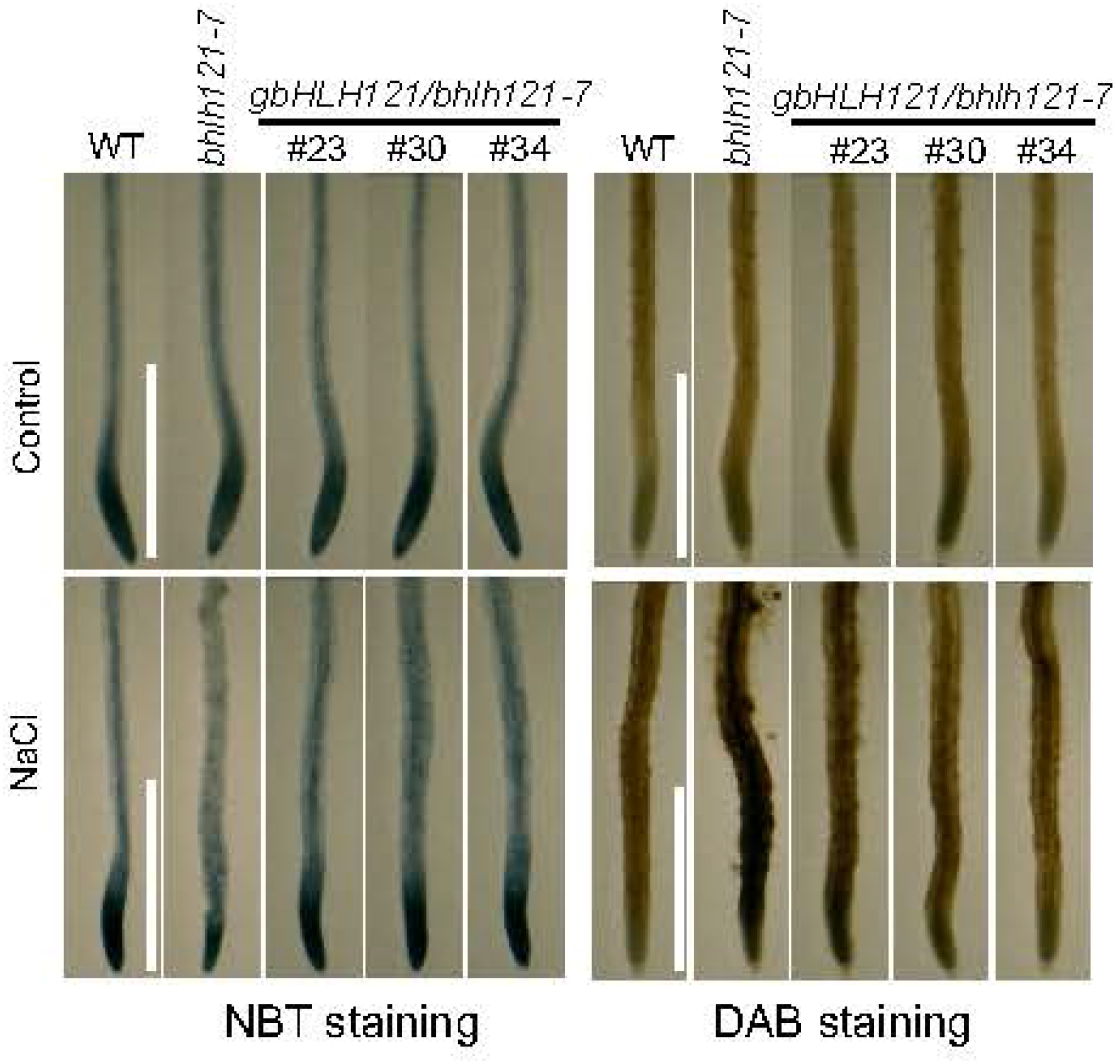
The full-length *bHLH121* gene rescues ROS imbalance in the *bhlh121-7* mutant under salt stress. Seedlings grown on ES medium supplemented with 100 mM NaCl for 10 days were sampled for ROS detection by NBT (O_2_^•–^) and DAB staining (H_2_O_2_).

**Supplementary Fig. 5.**
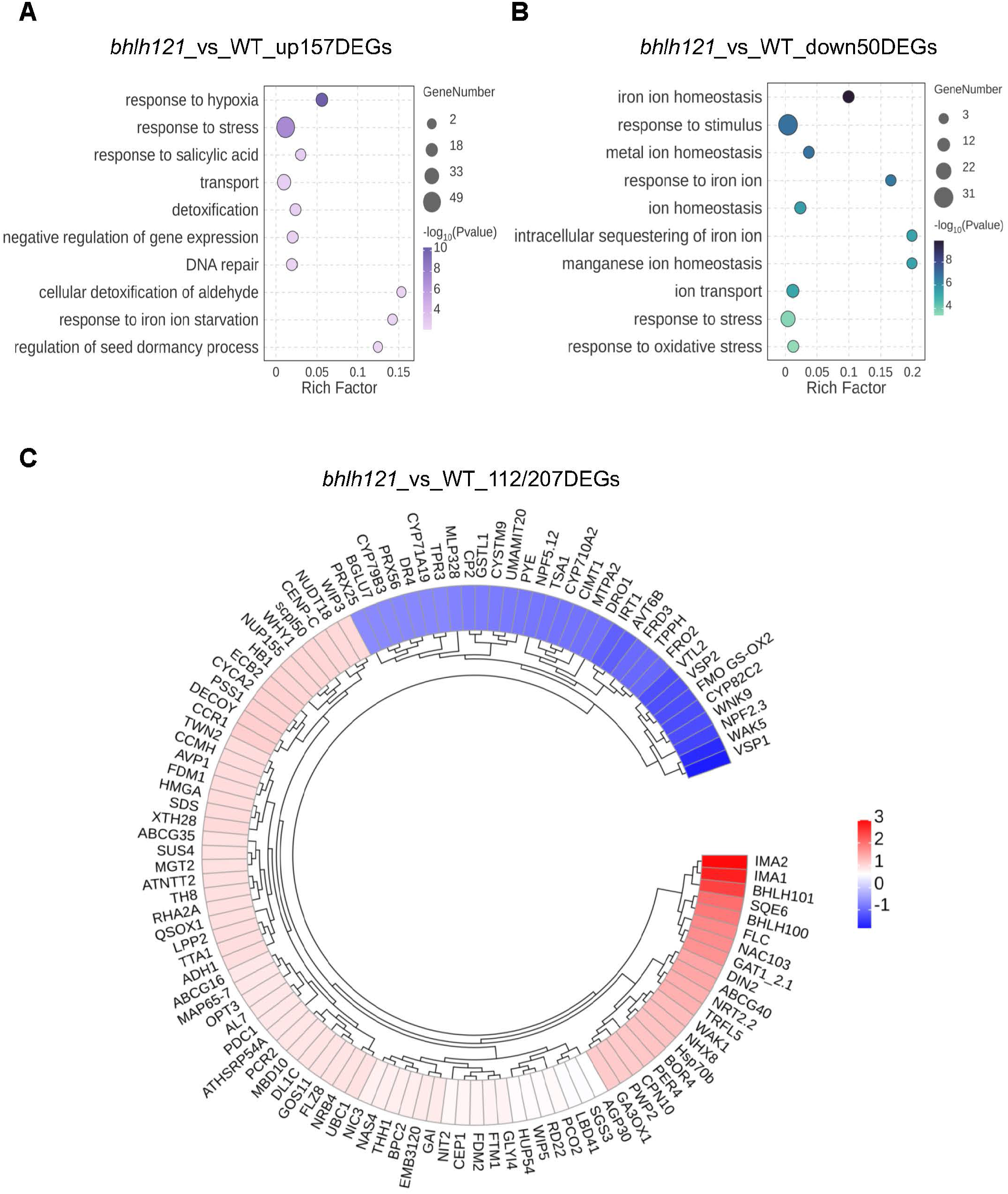
Gene Ontology (GO) and heatmap analysis of differentially expressed genes (DEGs) between WT and *bhlh121-7* mutant. (A) Top 10 GO terms of upregulated DEGs in *bhlh121* mutant compared to WT. (B) Top 10 GO terms of downregulated DEGs in *bhlh121* mutant compared to WT.(C) Heatmap results showing 112 representative genes out of 207 total DEGs between *bhlh121* and WT roots.

**Supplementary Fig. 6.**
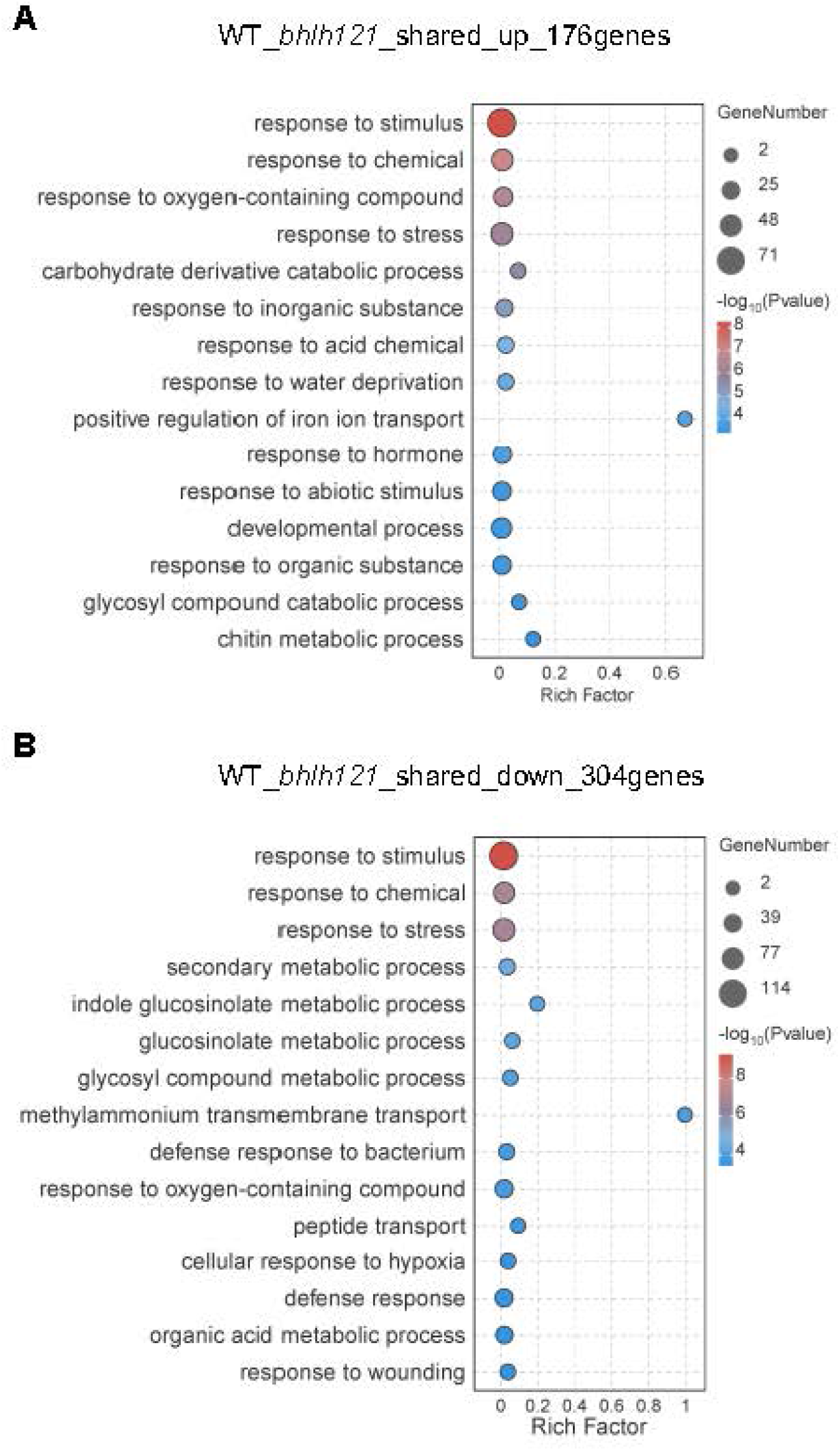
Gene Ontology (GO) enrichment analysis of shared differentially expressed genes (DEGs) in salt-stressed WT and *bhlh121-7* roots. (A) Top 15 GO terms of shared upregulated 176 DEGs in *bhlh121* mutant and WT. (B) Top 15 GO terms of shared downregulated 304 DEGs in *bhlh121* mutant and WT.

**Supplementary Fig. 7.**
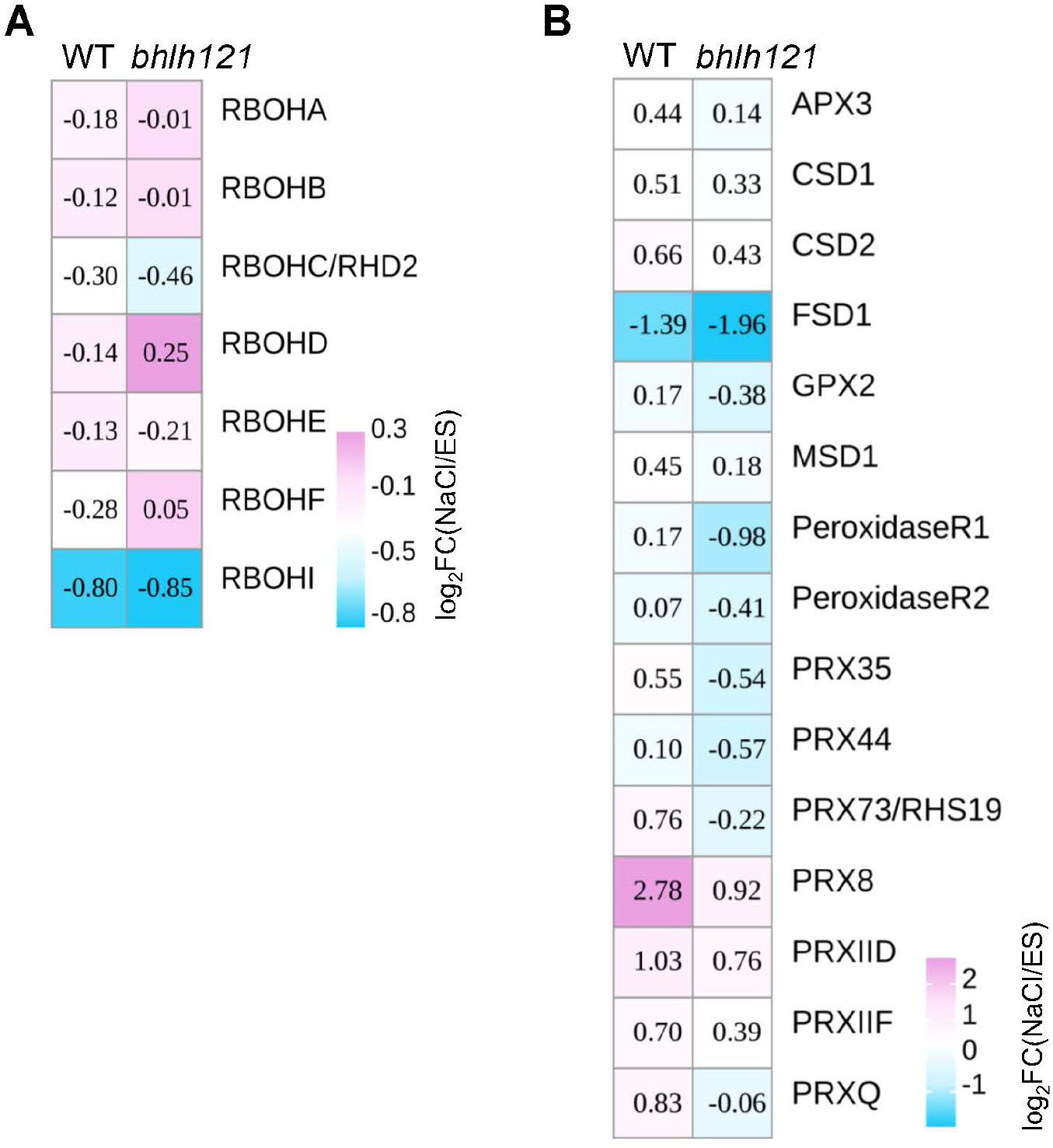
Heatmap analysis of ROS related genes in WT and *bhlh121* mutant under salt stress. Two manually curated lists of *RBOH* genes (A) and other genes in ROS homeostasis (B) were compared in WT and *bhlh121* mutant.

**Supplementary Fig. 8.**
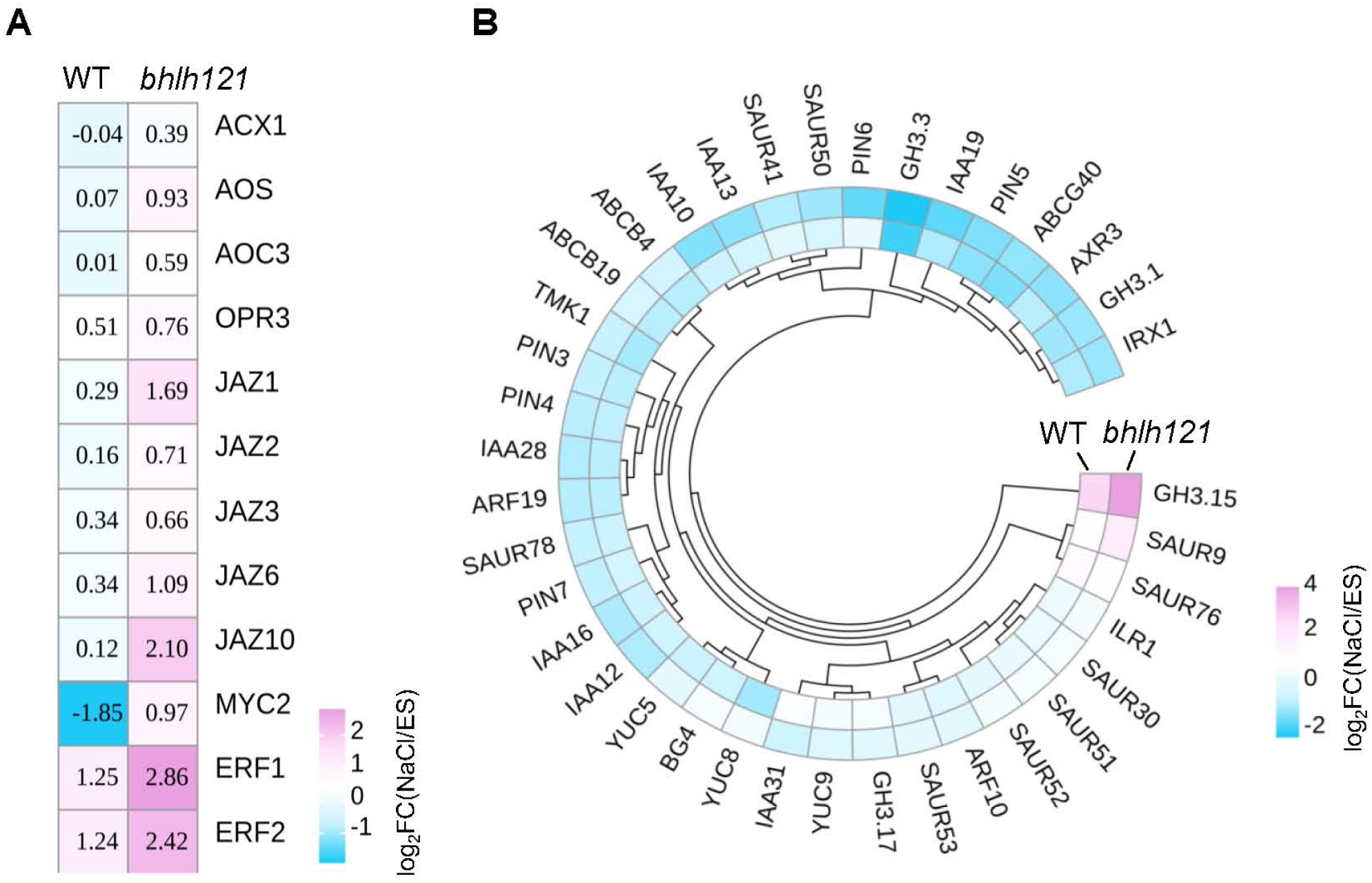
Transcriptome analyses indicate the involvement of jasmonic acid (JA) signaling and auxin homeostasis in the salt-sensitive phenotype of the *bhlh121-7* mutant. (A) Heatmap analysis of the expression changes of JA pathway genes in WT and *bhlh121* mutant under salt stress. (B) Heatmap analysis of the expression changes of auxin pathway genes in WT and *bhlh121* mutant under salt stress.

**Supplementary Fig. 9.**
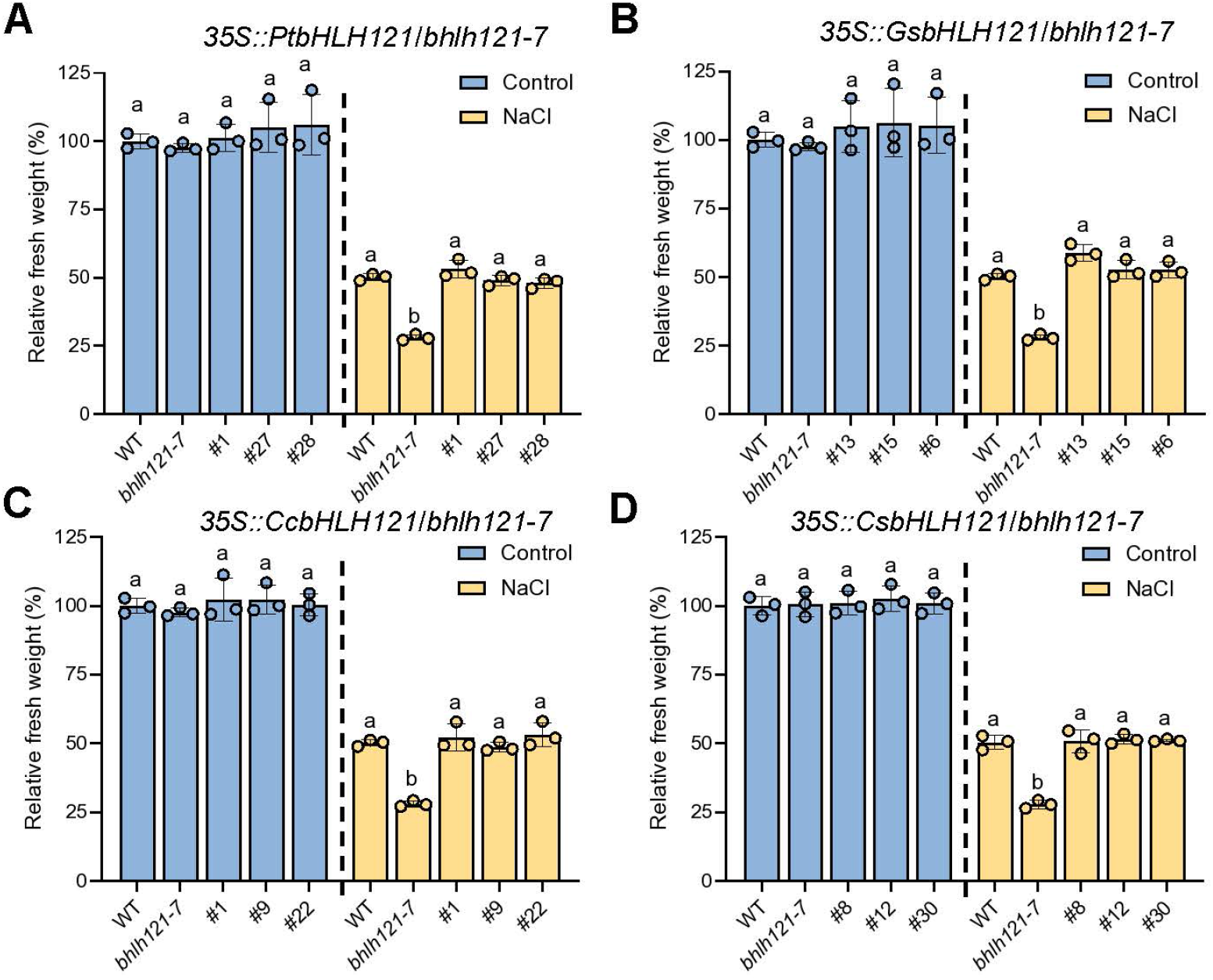
Effects of ectopic expression of *bHLH121* homologous genes in the *bhlh121-7* mutant under salt stress. (A-D) Relative fresh weight of 35S promoter driven *Populus PtbHLH121* (A), soybean *GsbHLH121* (B), Citrus *CcbHLH121* (C), and Cucumber *CsbHLH121* (D) overexpression transgenic lines in *bhlh121-7* background grown on ES medium supplemented with 100 mM NaCl for 10 days. The statistical differences were analyzed using a one-way ANOVA followed by Tukey’s multiple comparisons test. Different letters indicate significant differences (*P* < 0.05).

**Supplementary Fig. 10.**
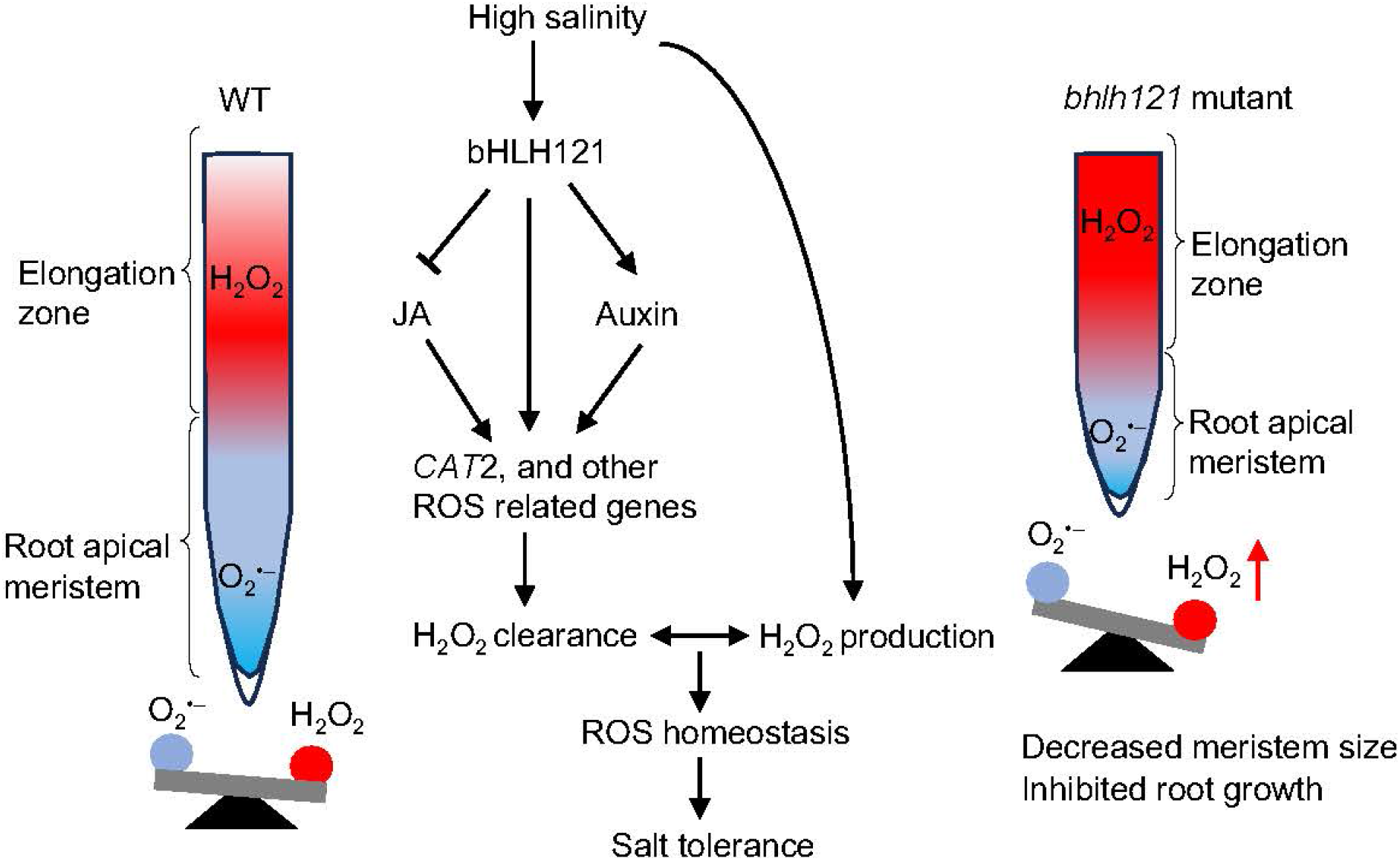
A proposed working model illustrating the role of bHLH121 in maintaining RAM size and root tip growth under salt stress in *Arabidopsis*. Under high salinity, bHLH121 positively regulates salt tolerance by orchestrating a complex regulatory network. bHLH121 promotes the expression of *CAT2* and other ROS-related genes directly, as well as indirectly through the regulation of jasmonate (JA) and auxin signaling pathways. This regulation leads to an increase in H_2_O_2_ clearance, effectively maintains ROS homeostasis and ultimately confers salt tolerance (middle panel). In wild-type (WT) roots, this bHLH121-dependent mechanism maintains a proper balance between O_2_^•-^ and H_2_O_2_, ensuring normal root apical meristem size and root elongation under salt stress (left panel). In contrast, loss of function of bHLH121 (*bhlh121* mutant) disrupts this balance, resulting in excessive H_2_O_2_ accumulation, reduced meristem size, and inhibited root growth, thereby decreasing salt tolerance (right panel). The balance scale at the bottom represents the relative ROS equilibrium. bHLH121 confers salt stress tolerance via direct regulation of CAT2, revealing a bHLH121–CAT2 module that operates independently of the bHLH121-mediated iron homeostasis pathway.

**Table S1.**
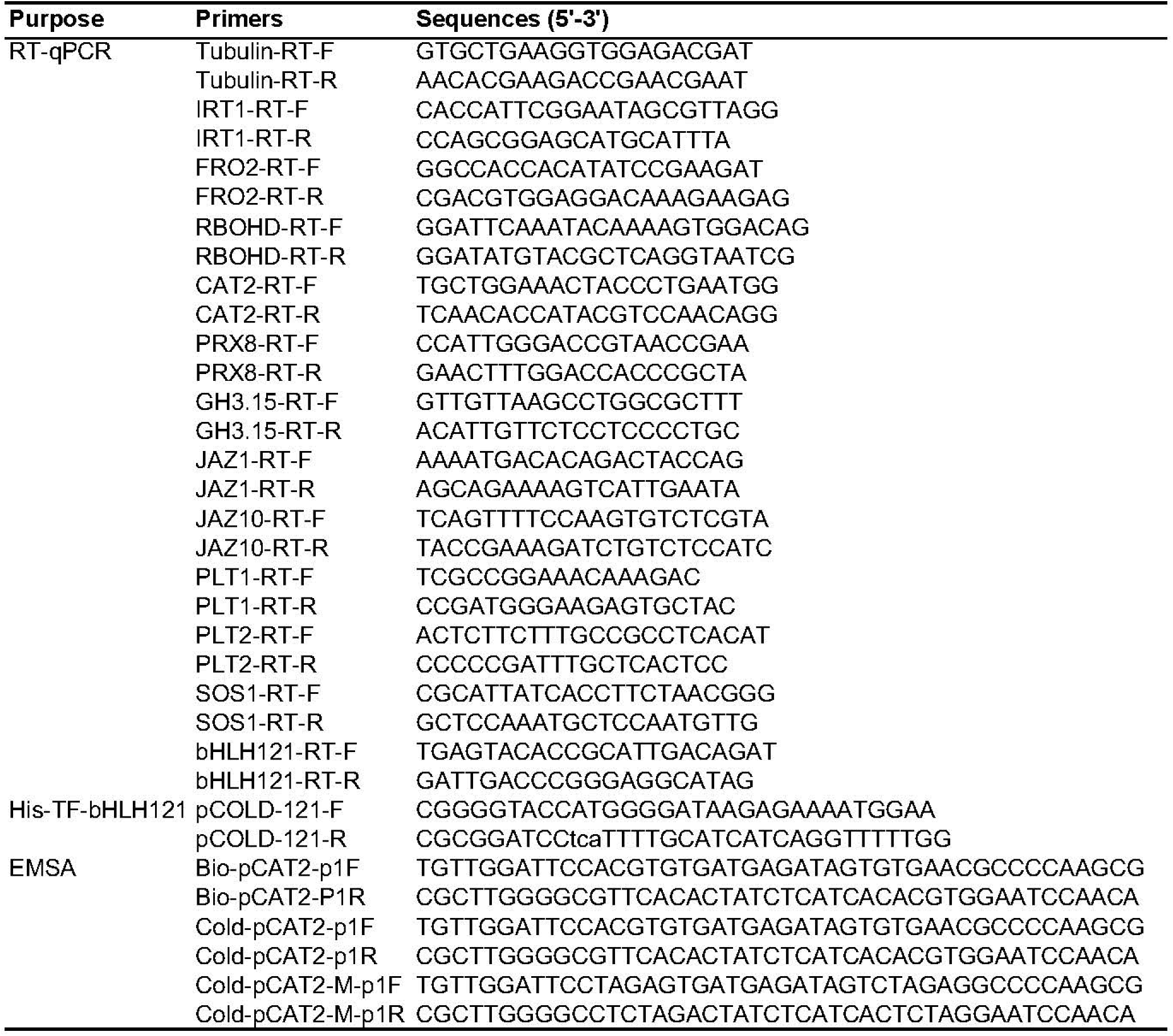
Oligos used in this study.

**Table S2.** List of differentially expressed genes in the roots between WT and *uri-2* mutant.

| gene_id | Col-0 | uri | log2(uri/Col-0) | q_value |
|---|---|---|---|---|
| AT1G47395 | 1.03123 | 8.60799 | 3.0613 | 0.00613065 |
| AT1G47400 | 0.929148 | 6.73302 | 2.85727 | 0.00613065 |
| AT5G04150 | 2.28779 | 13.0537 | 2.51242 | 0.0297101 |
| AT5G24160 | 0.250609 | 1.05812 | 2.07799 | 0.00613065 |
| AT2G41240 | 1.11389 | 4.36624 | 1.97078 | 0.00613065 |
| AT5G60610 | 0.470101 | 1.7541 | 1.89969 | 0.00613065 |
| AT2G17850 | 2.20114 | 8.14742 | 1.88809 | 0.0113181 |
| AT5G10140 | 1.35214 | 4.81935 | 1.83359 | 0.00613065 |
| AT5G64060 | 0.55852 | 1.91409 | 1.77698 | 0.0113181 |
| AT1G15040 | 3.9698 | 13.1282 | 1.72553 | 0.0427168 |
| AT1G65370 | 5.287 | 15.922 | 1.5905 | 0.00613065 |
| AT3G60140 | 0.473419 | 1.36727 | 1.53011 | 0.00613065 |
| AT1G15520 | 0.375568 | 1.07944 | 1.52313 | 0.00613065 |
| AT1G08100 | 2.02864 | 5.50907 | 1.4413 | 0.00613065 |
| AT1G15720 | 2.40289 | 6.45866 | 1.42646 | 0.00613065 |
| AT1G15625 | 9.31559 | 23.0801 | 1.30893 | 0.00613065 |
| AT1G21250 | 0.385376 | 0.949515 | 1.30092 | 0.0427168 |
| AT1G14660 | 1.63252 | 4.01412 | 1.29799 | 0.00613065 |
| AT5G05060 | 6.78054 | 16.3003 | 1.26543 | 0.00613065 |
| AT2G43620 | 6.08408 | 14.5548 | 1.25838 | 0.00613065 |
| AT1G15480 | 10.6288 | 25.283 | 1.2502 | 0.00613065 |
| AT1G16030 | 3.30785 | 7.81769 | 1.24085 | 0.00613065 |
| AT1G15385 | 3.30037 | 7.74318 | 1.2303 | 0.0254938 |
| AT1G15460 | 7.45662 | 17.4934 | 1.23022 | 0.00613065 |
| AT1G15710 | 9.97279 | 23.0323 | 1.20759 | 0.00613065 |
| AT1G14630 | 10.1644 | 23.3963 | 1.20275 | 0.00613065 |
| AT1G15420 | 28.3315 | 65.089 | 1.20001 | 0.00613065 |
| AT1G14770 | 4.41913 | 10.1143 | 1.19457 | 0.00613065 |
| AT1G14540 | 23.1702 | 52.9211 | 1.19157 | 0.00613065 |
| AT1G15630 | 2.05743 | 4.66559 | 1.18121 | 0.00613065 |
| AT1G14550 | 8.65982 | 19.5708 | 1.1763 | 0.00613065 |
| AT1G14980 | 178.528 | 401.168 | 1.16806 | 0.00613065 |
| AT1G16000 | 135.376 | 303.935 | 1.16679 | 0.00613065 |
| AT1G15140 | 14.5142 | 32.4778 | 1.16199 | 0.00613065 |
| AT1G15440 | 20.4853 | 45.7037 | 1.15772 | 0.00613065 |
| AT1G15550 | 39.4531 | 87.9834 | 1.15709 | 0.00613065 |
| AT1G14910 | 19.6677 | 43.6327 | 1.14958 | 0.00613065 |
| AT1G15640 | 3.79224 | 8.35229 | 1.13912 | 0.00613065 |
| AT1G14810 | 82.3863 | 181.397 | 1.13868 | 0.00613065 |
| AT1G14710 | 34.5778 | 75.8859 | 1.13399 | 0.00613065 |
| AT1G15530 | 5.92009 | 12.9749 | 1.13203 | 0.00613065 |
| AT1G15810 | 32.6303 | 71.0722 | 1.12307 | 0.00613065 |
| AT1G15950 | 197.968 | 429.842 | 1.11854 | 0.00613065 |
| AT1G14620 | 46.452 | 100.847 | 1.11835 | 0.00613065 |
| AT1G15930 | 499.251 | 1083.3 | 1.1176 | 0.00613065 |
| AT1G14650 | 28.0519 | 60.8351 | 1.11681 | 0.00613065 |
| AT1G14670 | 149.818 | 324.885 | 1.11672 | 0.00613065 |
| AT1G15110 | 23.6822 | 50.8335 | 1.10198 | 0.00613065 |
| AT1G15570 | 10.7701 | 23.0858 | 1.09998 | 0.00613065 |
| AT1G15510 | 1.79929 | 3.83966 | 1.09355 | 0.00613065 |
| AT1G14560 | 12.9557 | 27.2757 | 1.07403 | 0.00613065 |
| AT1G16040 | 12.1662 | 25.4893 | 1.06702 | 0.00613065 |
| AT1G15940 | 10.1976 | 21.1912 | 1.05524 | 0.00613065 |
| AT2G16060 | 124.882 | 259.193 | 1.05346 | 0.00613065 |
| AT1G14850 | 14.2354 | 29.431 | 1.04785 | 0.00613065 |
| AT1G14410 | 8.13422 | 16.8103 | 1.04727 | 0.00613065 |
| AT1G15000 | 10.9505 | 22.6091 | 1.0459 | 0.00613065 |
| AT1G15470 | 23.4364 | 48.3829 | 1.04575 | 0.00613065 |
| AT1G15130 | 49.8049 | 102.151 | 1.03635 | 0.00613065 |
| AT1G15660 | 12.6364 | 25.8213 | 1.03098 | 0.00613065 |
| AT1G15790 | 6.64976 | 13.5295 | 1.02473 | 0.00613065 |
| AT1G15290 | 10.4625 | 21.2704 | 1.02362 | 0.00613065 |
| AT1G14860 | 19.8462 | 40.2901 | 1.02156 | 0.00613065 |
| AT1G14590 | 4.32273 | 8.76692 | 1.02013 | 0.00613065 |
| AT4G10265 | 5.27224 | 10.6518 | 1.01461 | 0.0297101 |
| AT1G15270 | 417.808 | 842.578 | 1.01197 | 0.00613065 |
| AT1G14570 | 48.0594 | 96.7625 | 1.00963 | 0.00613065 |
| AT1G14960 | 16.0041 | 32.1975 | 1.0085 | 0.00613065 |
| AT1G14720 | 41.2375 | 82.8062 | 1.00578 | 0.00613065 |
| AT1G14750 | 1.96236 | 3.92672 | 1.00073 | 0.00613065 |
| AT1G15120 | 133.297 | 266.518 | 0.999592 | 0.00613065 |
| AT1G14900 | 58.0641 | 115.797 | 0.995882 | 0.00613065 |
| AT1G15910 | 13.7589 | 27.374 | 0.992445 | 0.00613065 |
| AT1G15690 | 438.102 | 869.776 | 0.989378 | 0.00613065 |
| AT1G15370 | 58.6392 | 116.354 | 0.988579 | 0.00613065 |
| AT1G15220 | 5.89613 | 11.6848 | 0.986792 | 0.0165233 |
| AT1G14610 | 63.2641 | 125.31 | 0.986038 | 0.00613065 |
| AT1G15730 | 4.00126 | 7.88289 | 0.978272 | 0.00613065 |
| AT2G24600 | 2.01422 | 3.95548 | 0.973635 | 0.00613065 |
| AT1G77120 | 141.829 | 278.13 | 0.971608 | 0.00613065 |
| AT1G15030 | 18.1656 | 35.6073 | 0.97096 | 0.00613065 |
| AT1G14740 | 45.3128 | 88.8156 | 0.970892 | 0.00613065 |
| AT1G15080 | 6.81557 | 13.3587 | 0.97087 | 0.00613065 |
| AT1G15860 | 42.4135 | 82.9638 | 0.967958 | 0.00613065 |
| AT1G15670 | 70.3279 | 137.544 | 0.967728 | 0.00613065 |
| AT1G15020 | 19.1358 | 37.2415 | 0.96064 | 0.00613065 |
| AT1G14450 | 294.61 | 571.784 | 0.956666 | 0.00613065 |
| AT1G16020 | 12.8386 | 24.8326 | 0.951742 | 0.00613065 |
| AT1G15100 | 122.102 | 235.858 | 0.949839 | 0.00613065 |
| AT1G69880 | 14.9813 | 28.8572 | 0.94577 | 0.00613065 |
| AT1G15500 | 142.773 | 274.932 | 0.945353 | 0.00613065 |
| AT1G16010 | 31.5376 | 60.6062 | 0.942391 | 0.00613065 |
| AT3G43190 | 34.0309 | 65.3793 | 0.941991 | 0.00613065 |
| AT1G15210 | 88.6221 | 170.029 | 0.940041 | 0.00613065 |
| AT1G14460 | 7.8694 | 15.0753 | 0.937862 | 0.00613065 |
| AT1G15410 | 8.44945 | 16.1755 | 0.936882 | 0.00613065 |
| AT1G34510 | 3.40208 | 6.47833 | 0.929207 | 0.0211272 |
| AT1G15400 | 99.1263 | 188.488 | 0.927134 | 0.00613065 |
| AT1G14400 | 199.291 | 378.023 | 0.923599 | 0.00613065 |
| AT2G18480 | 1.29448 | 2.45241 | 0.921828 | 0.0254938 |
| AT1G15780 | 19.1635 | 36.3032 | 0.921736 | 0.00613065 |
| AT1G14600 | 2.55699 | 4.83211 | 0.918208 | 0.0211272 |
| AT1G14970 | 6.18875 | 11.6794 | 0.916242 | 0.00613065 |
| AT3G58270 | 4.13465 | 7.79606 | 0.91498 | 0.00613065 |
| AT3G22550 | 6.13976 | 11.5381 | 0.910156 | 0.00613065 |
| AT1G15880 | 39.1962 | 73.6257 | 0.909494 | 0.00613065 |
| AT1G15280 | 25.4161 | 47.7118 | 0.908603 | 0.00613065 |
| AT1G14830 | 86.0425 | 161.234 | 0.906034 | 0.00613065 |
| AT1G14820 | 12.075 | 22.5293 | 0.899784 | 0.00613065 |
| AT1G15340 | 51.4744 | 95.9516 | 0.898453 | 0.00613065 |
| AT1G14870 | 209.253 | 389.073 | 0.89479 | 0.00613065 |
| AT1G15060 | 12.2303 | 22.6762 | 0.890716 | 0.00613065 |
| AT1G15240 | 14.4496 | 26.735 | 0.887696 | 0.00613065 |
| AT1G15310 | 4.89524 | 9.03657 | 0.884396 | 0.00613065 |
| AT4G33070 | 46.3816 | 85.3667 | 0.880122 | 0.00613065 |
| AT1G14510 | 39.0631 | 71.8778 | 0.879742 | 0.00613065 |
| AT4G16370 | 12.5362 | 23.0222 | 0.876932 | 0.00613065 |
| AT1G14690 | 21.8092 | 40.0374 | 0.876412 | 0.00613065 |
| AT3G55090 | 1.61618 | 2.96624 | 0.876051 | 0.0211272 |
| AT1G15170 | 12.8843 | 23.518 | 0.868149 | 0.00613065 |
| AT1G15350 | 61.5213 | 112.191 | 0.866802 | 0.00613065 |
| AT1G14480 | 3.13426 | 5.70441 | 0.863953 | 0.00613065 |
| AT1G15200 | 33.3589 | 60.5652 | 0.860419 | 0.00613065 |
| AT1G15920 | 18.1548 | 32.5909 | 0.84412 | 0.00613065 |
| AT1G14920 | 14.2957 | 25.5369 | 0.837001 | 0.00613065 |
| AT3G14900 | 2.23171 | 3.984 | 0.836069 | 0.0427168 |
| AT1G15490 | 13.9759 | 24.9049 | 0.833487 | 0.0211272 |
| AT1G14890 | 20.3437 | 36.1427 | 0.829123 | 0.00613065 |
| AT3G22060 | 5.68126 | 10.0099 | 0.81715 | 0.0113181 |
| AT1G14685 | 30.204 | 53.0641 | 0.812997 | 0.00613065 |
| AT1G15970 | 12.6395 | 22.2047 | 0.812928 | 0.00613065 |
| AT1G14530 | 11.4117 | 20.0349 | 0.812005 | 0.00613065 |
| AT1G15230 | 27.0258 | 47.1862 | 0.804025 | 0.00613065 |
| AT1G56430 | 8.52337 | 14.8483 | 0.800802 | 0.00613065 |
| AT5G23220 | 15.9536 | 27.616 | 0.791622 | 0.00613065 |
| AT1G15890 | 3.55166 | 6.11078 | 0.782863 | 0.00613065 |
| AT1G15380 | 194.416 | 329.818 | 0.762522 | 0.00613065 |
| AT1G43800 | 98.4506 | 166.772 | 0.760405 | 0.00613065 |
| AT1G15760 | 6.81349 | 11.4569 | 0.749746 | 0.0113181 |
| AT4G00380 | 2.72523 | 4.57662 | 0.747907 | 0.0344959 |
| AT5G50260 | 4.59171 | 7.70559 | 0.746875 | 0.0297101 |
| AT4G14020 | 19.7487 | 33.0979 | 0.744983 | 0.0113181 |
| AT3G44300 | 27.1384 | 44.8407 | 0.724472 | 0.00613065 |
| AT1G14730 | 48.5784 | 79.9284 | 0.718394 | 0.00613065 |
| AT3G10040 | 8.29197 | 13.638 | 0.717841 | 0.00613065 |
| AT1G19530 | 31.0902 | 50.982 | 0.713531 | 0.00613065 |
| AT3G28580 | 5.83045 | 9.56047 | 0.713474 | 0.0427168 |
| AT5G39890 | 24.1367 | 39.199 | 0.699586 | 0.00613065 |
| AT1G14580 | 5.16217 | 8.32888 | 0.690144 | 0.0211272 |
| AT5G25610 | 5.37131 | 8.62891 | 0.683905 | 0.0211272 |
| AT4G33560 | 47.5206 | 76.0476 | 0.678351 | 0.00613065 |
| AT1G30760 | 9.75843 | 15.5985 | 0.676686 | 0.00613065 |
| AT4G27450 | 62.1218 | 98.8377 | 0.669962 | 0.00613065 |
| AT2G33790 | 60.7682 | 94.6934 | 0.639948 | 0.00613065 |
| AT5G23570 | 5.80771 | 8.92666 | 0.620152 | 0.0254938 |
| AT3G02550 | 39.2398 | 60.1715 | 0.616763 | 0.00613065 |
| AT1G77145 | 12.8848 | 19.6746 | 0.61066 | 0.0211272 |
| AT2G41480 | 20.7285 | 13.5275 | -0.615717 | 0.0254938 |
| AT3G62740 | 17.2671 | 11.1998 | -0.624563 | 0.0113181 |
| AT2G22330 | 60.814 | 39.3834 | -0.626817 | 0.00613065 |
| AT3G26460 | 29.0554 | 18.786 | -0.629148 | 0.0254938 |
| AT5G15180 | 40.5213 | 26.1708 | -0.630723 | 0.0113181 |
| AT1G73330 | 624.771 | 402.891 | -0.632937 | 0.00613065 |
| AT4G13290 | 11.3421 | 7.25524 | -0.644597 | 0.0113181 |
| AT5G27030 | 13.7892 | 8.81999 | -0.644694 | 0.00613065 |
| AT3G12900 | 14.1395 | 9.0431 | -0.644846 | 0.0461861 |
| AT2G01520 | 635.054 | 395.675 | -0.682562 | 0.0165233 |
| AT5G54700 | 14.3818 | 8.65487 | -0.732661 | 0.00613065 |
| AT1G14120 | 20.2735 | 12.1588 | -0.737596 | 0.00613065 |
| AT4G11320 | 372.05 | 222.167 | -0.743854 | 0.00613065 |
| AT5G02780 | 27.9311 | 16.5936 | -0.751247 | 0.00613065 |
| AT1G73120 | 62.9357 | 37.2449 | -0.756836 | 0.00613065 |
| AT3G20340 | 24.8679 | 14.6638 | -0.762024 | 0.0113181 |
| AT3G22240 | 55.0221 | 32.2037 | -0.772783 | 0.0461861 |
| AT4G08290 | 12.2207 | 7.10545 | -0.782326 | 0.0254938 |
| AT3G49160 | 4.25018 | 2.46088 | -0.788351 | 0.0211272 |
| AT3G47640 | 10.1926 | 5.84962 | -0.801112 | 0.0254938 |
| AT1G72140 | 9.86448 | 5.55051 | -0.829622 | 0.00613065 |
| AT1G52410 | 3.35209 | 1.85636 | -0.852586 | 0.0113181 |
| AT2G34490 | 6.41321 | 3.54437 | -0.855518 | 0.00613065 |
| AT5G37990 | 40.6422 | 22.4488 | -0.856344 | 0.00613065 |
| AT3G58810 | 23.3966 | 12.8145 | -0.86852 | 0.00613065 |
| AT4G23680 | 26.359 | 14.4092 | -0.871309 | 0.00613065 |
| AT1G72490 | 3.99352 | 2.14559 | -0.896287 | 0.0113181 |
| AT5G38000 | 5.83722 | 3.03904 | -0.941665 | 0.0165233 |
| AT1G77520 | 11.6097 | 6.00682 | -0.950662 | 0.00613065 |
| AT3G08040 | 29.5889 | 15.2733 | -0.954041 | 0.00613065 |
| AT1G26240 | 2.70535 | 1.38874 | -0.96204 | 0.0461861 |
| AT4G39770 | 5.74917 | 2.93487 | -0.970059 | 0.0113181 |
| AT1G01580 | 103.509 | 52.4795 | -0.979924 | 0.00613065 |
| AT4G14060 | 24.8019 | 12.0472 | -1.04176 | 0.00613065 |
| AT4G19690 | 207.362 | 100.365 | -1.04689 | 0.00613065 |
| AT5G38820 | 4.80123 | 2.28234 | -1.07289 | 0.00613065 |
| AT2G27402 | 6.64087 | 3.08277 | -1.10715 | 0.00613065 |
| AT2G35736 | 33.8783 | 14.5214 | -1.22217 | 0.00613065 |
| AT1G76800 | 28.8382 | 12.2697 | -1.23289 | 0.00613065 |
| AT5G24770 | 24.3481 | 10.264 | -1.24621 | 0.00613065 |
| AT1G62540 | 2.36524 | 0.992223 | -1.25325 | 0.0113181 |
| AT5G28080 | 5.5502 | 2.30932 | -1.26507 | 0.00613065 |
| AT4G31970 | 4.13761 | 1.70468 | -1.2793 | 0.00613065 |
| AT1G21140 | 62.0205 | 25.0227 | -1.30951 | 0.00613065 |
| AT3G25190 | 180.017 | 72.3508 | -1.31505 | 0.00613065 |
| AT3G45680 | 4.5897 | 1.79058 | -1.35797 | 0.00613065 |
| AT1G21230 | 2.0213 | 0.6509 | -1.63477 | 0.00613065 |
| AT5G24780 | 10.4958 | 3.05583 | -1.78018 | 0.00613065 |
| AT3G33528 | 3.56218 | 0.996558 | -1.83774 | 0.00613065 |
| AT4G09200 | 2.43027 | 0.145525 | -4.06177 | 0.00613065 |

**Table S3.**
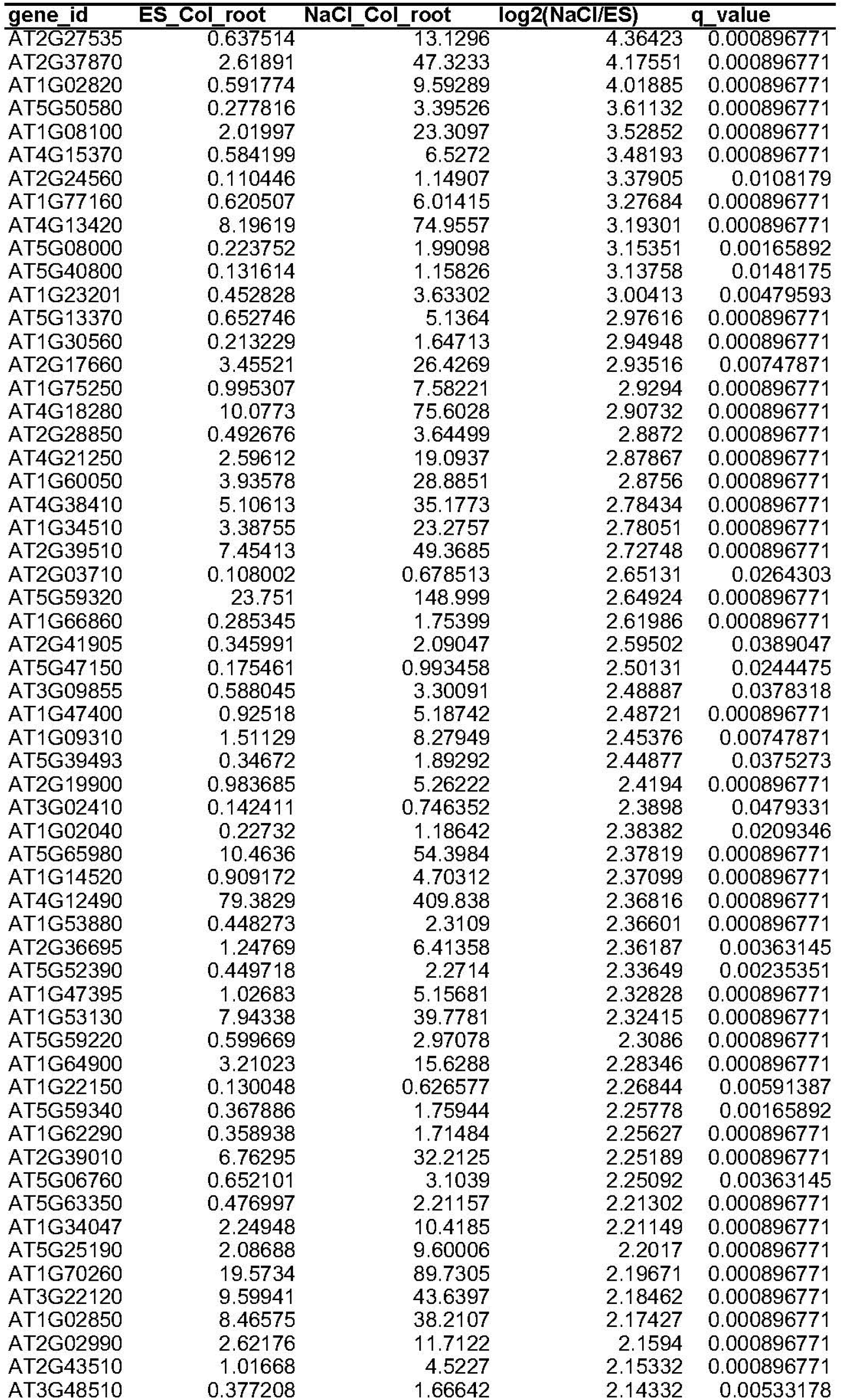
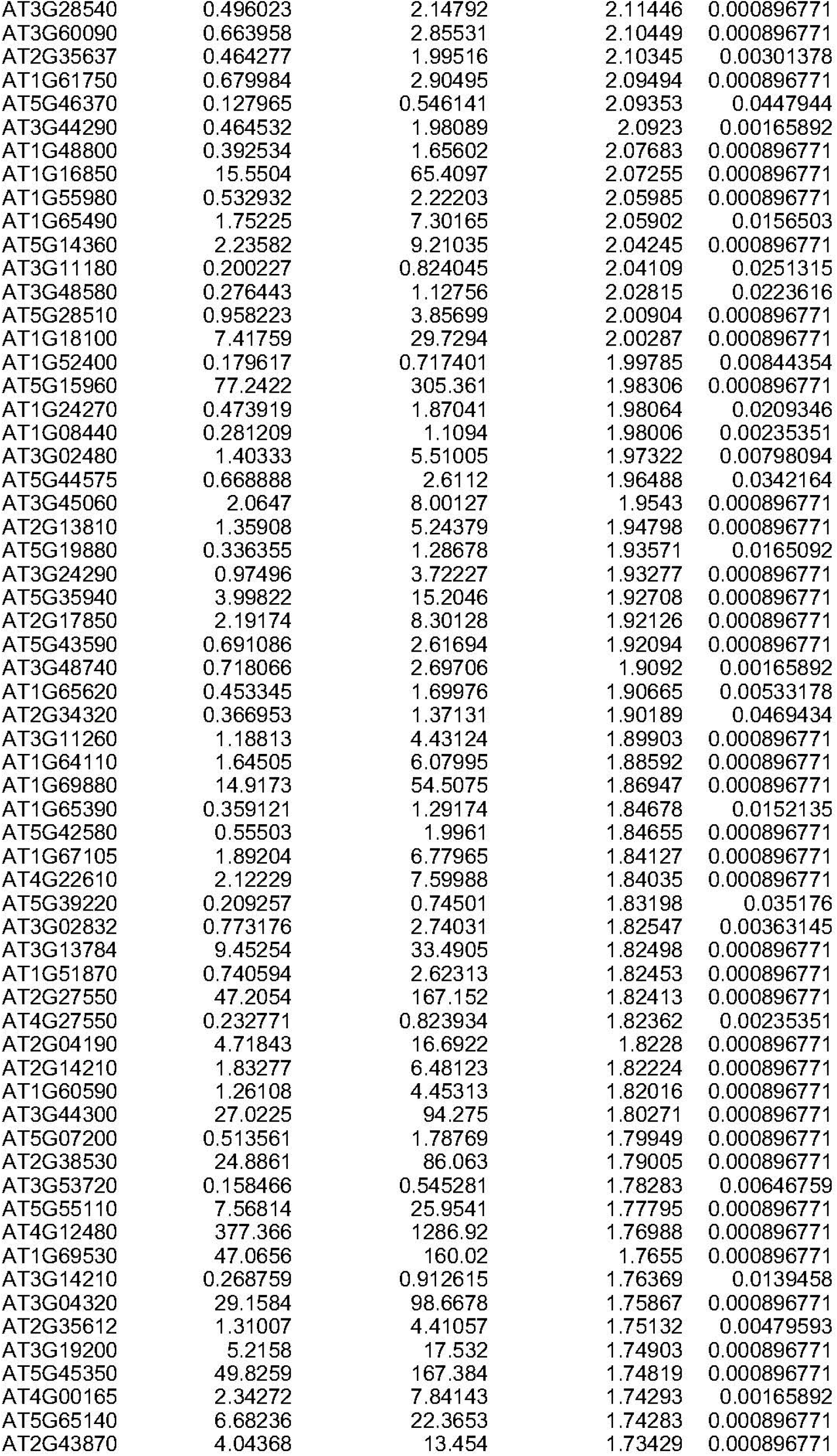
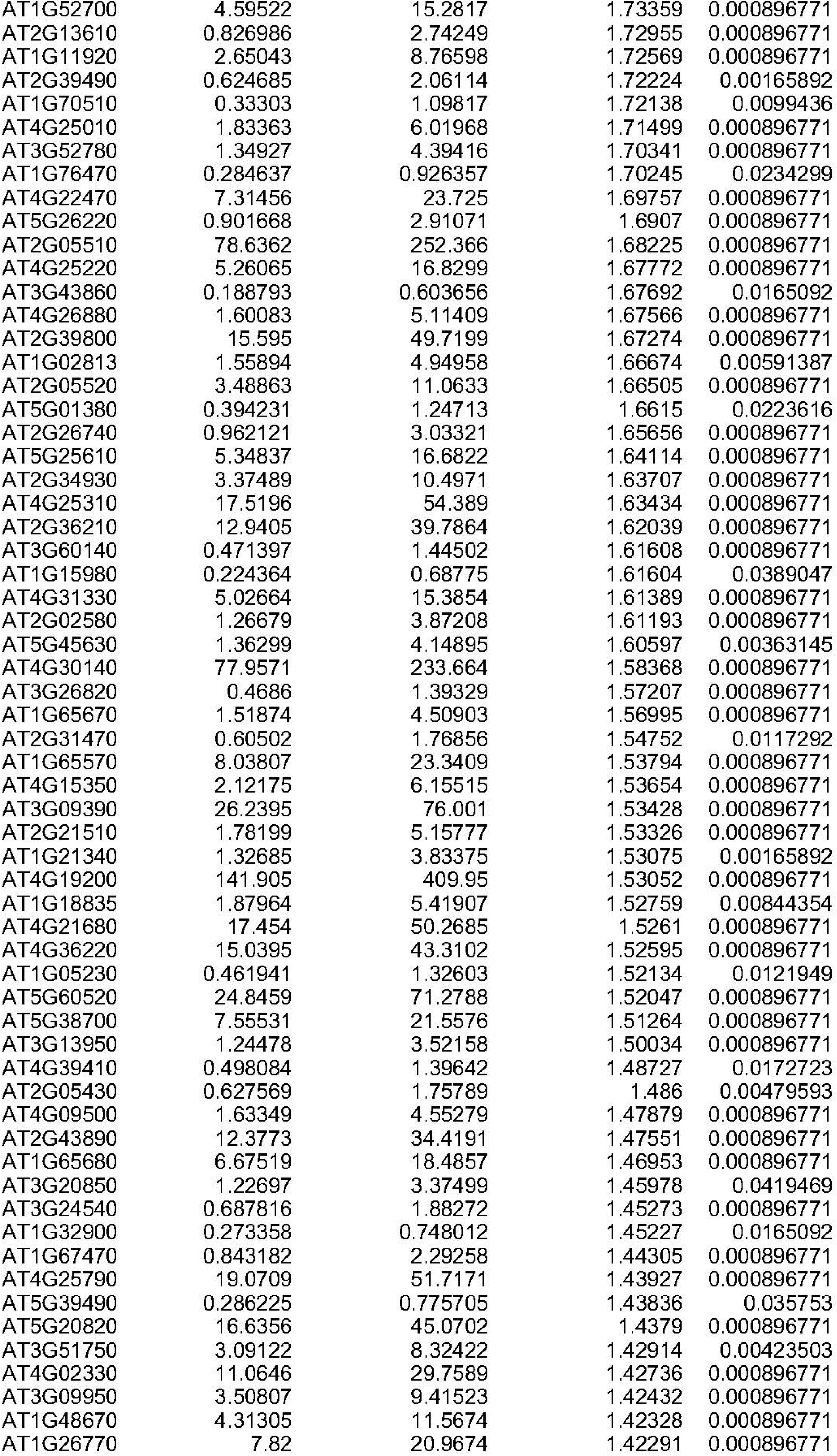
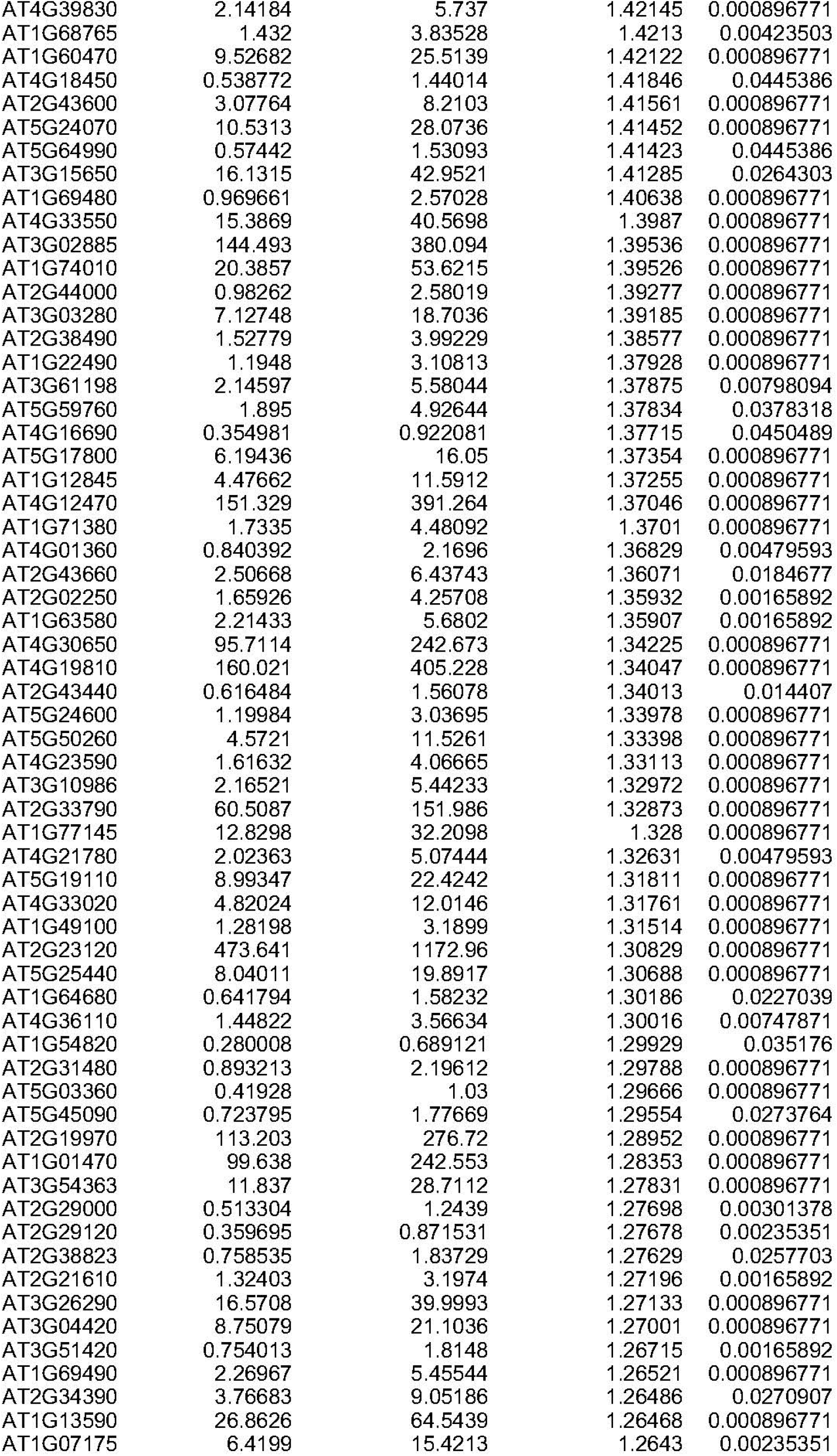
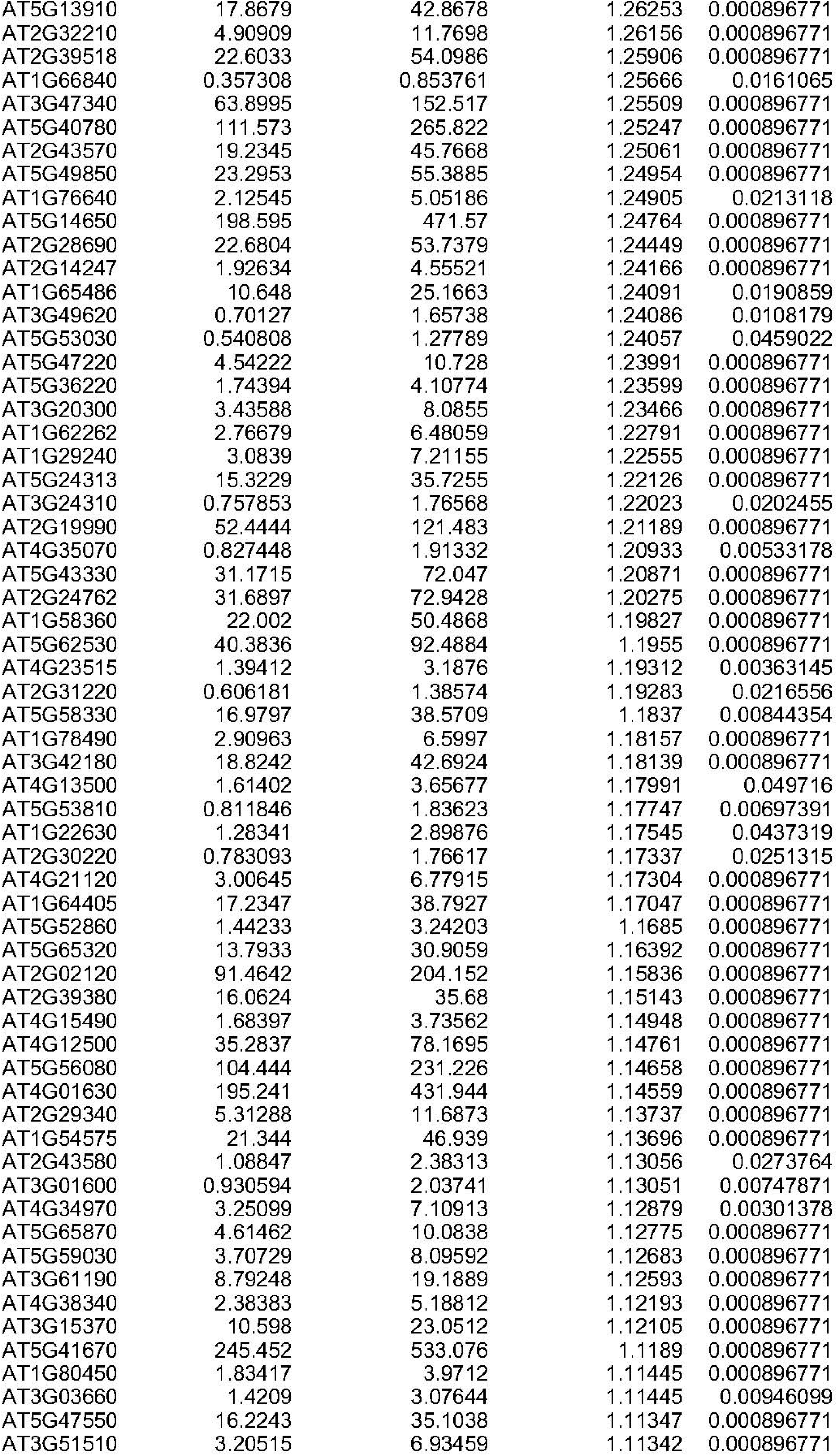
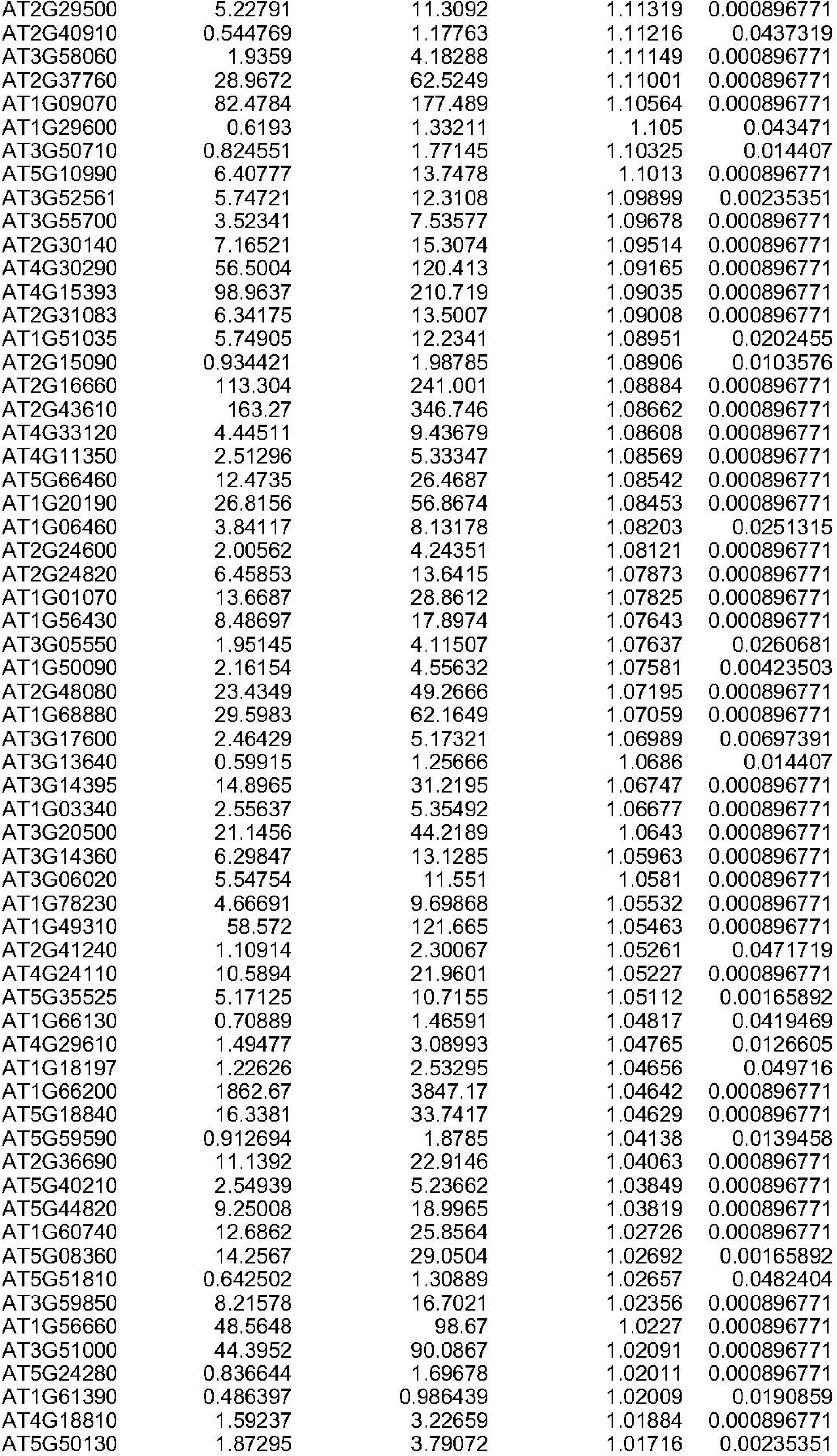
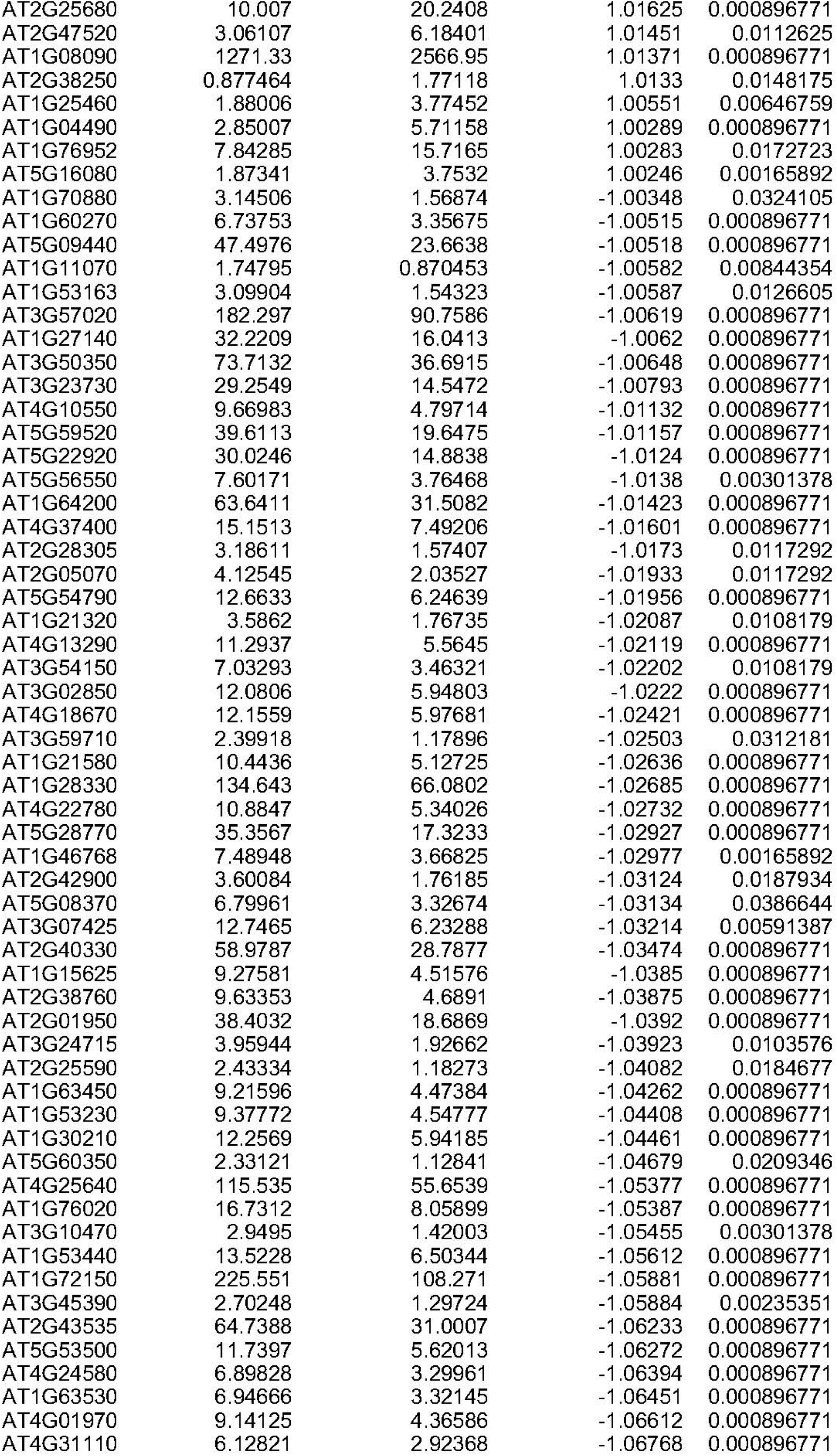
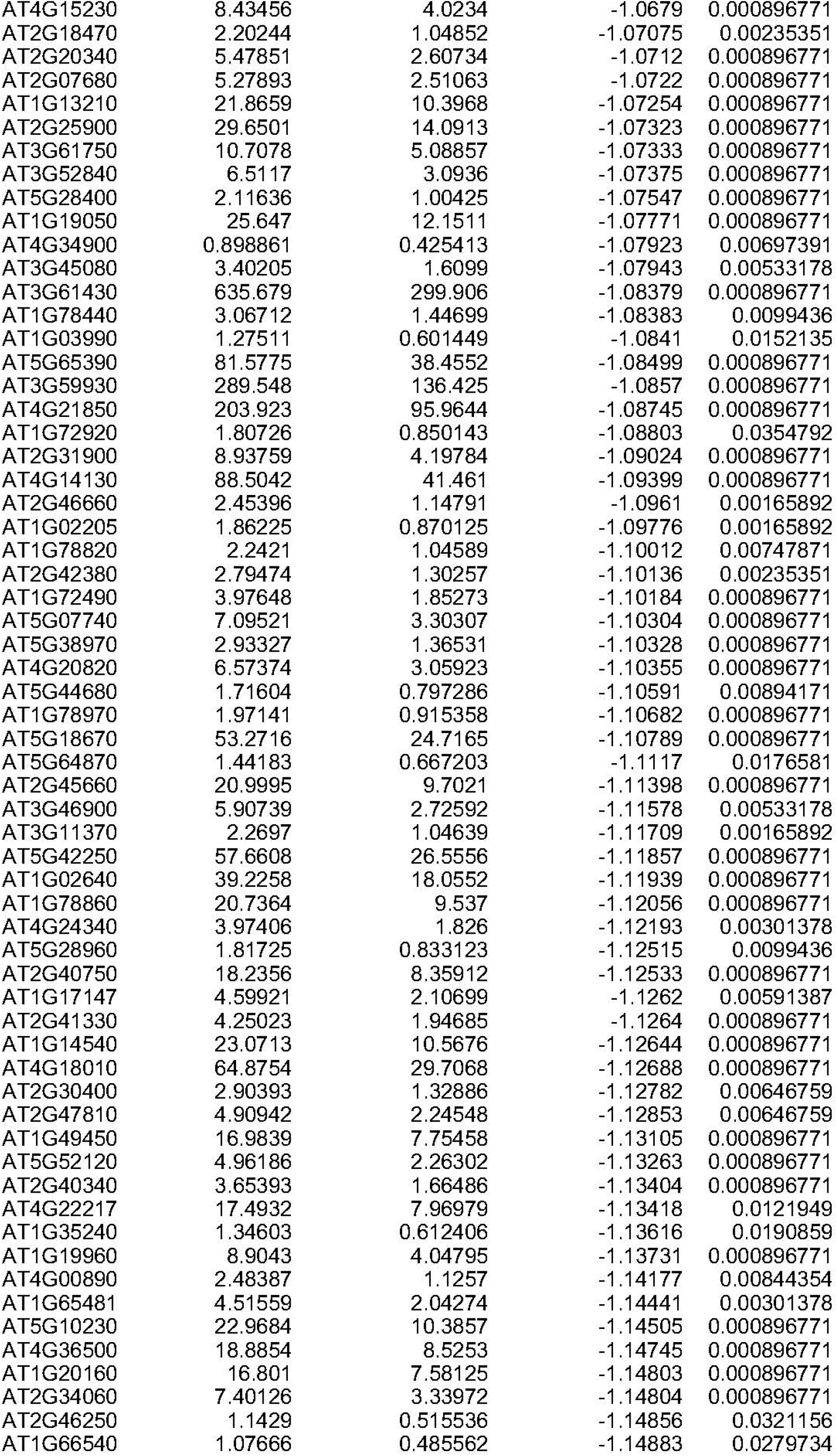
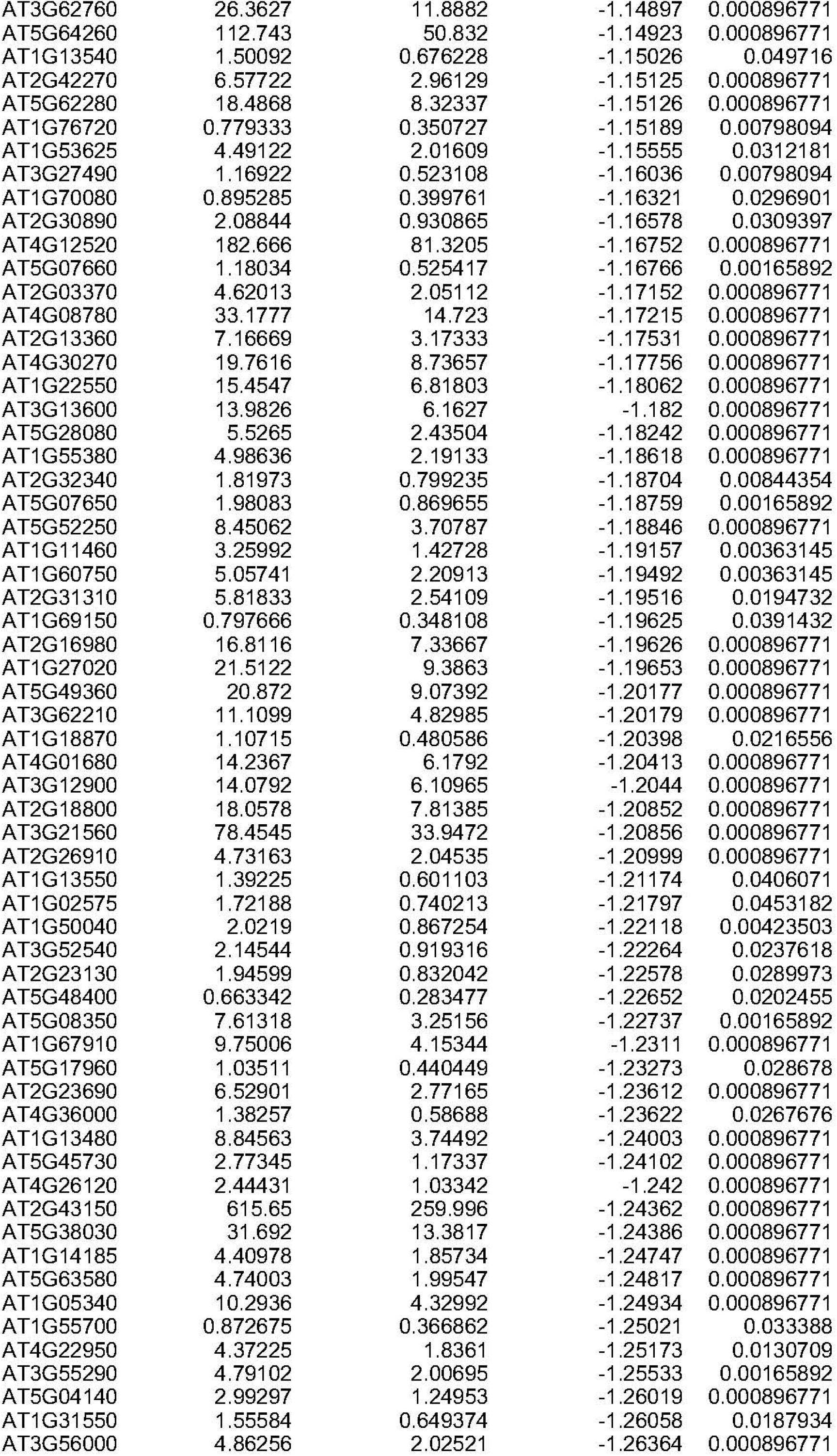
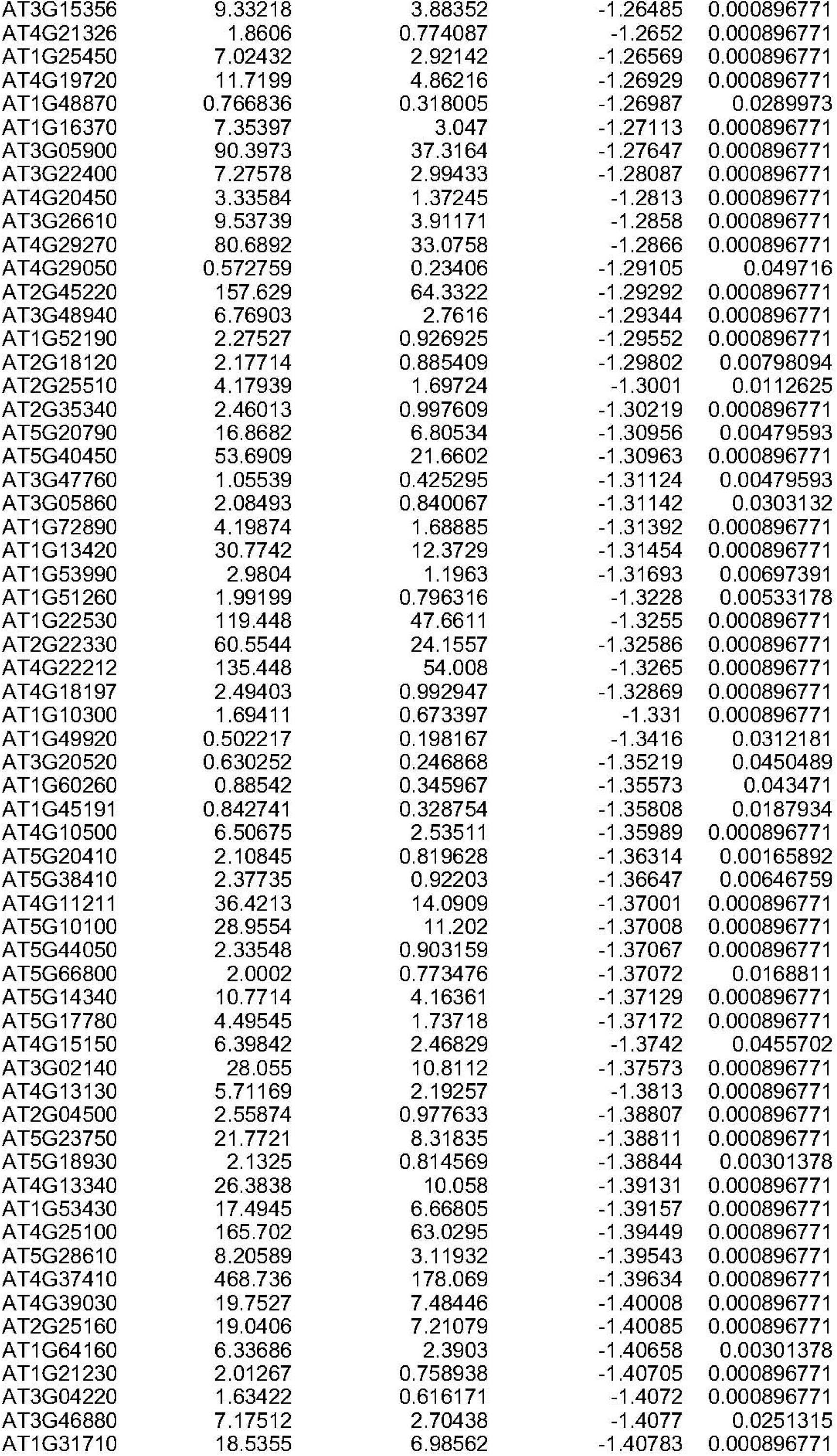
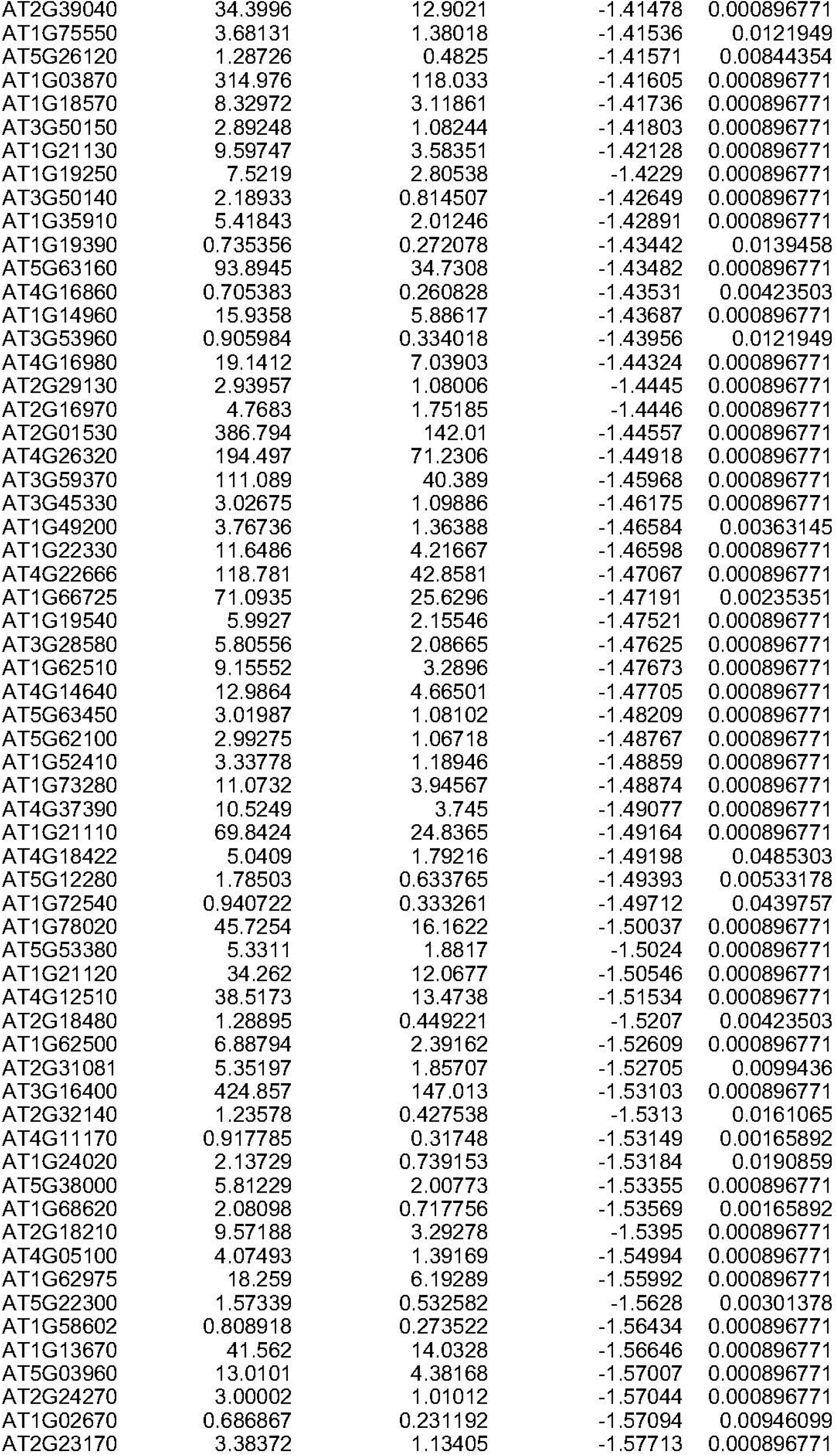
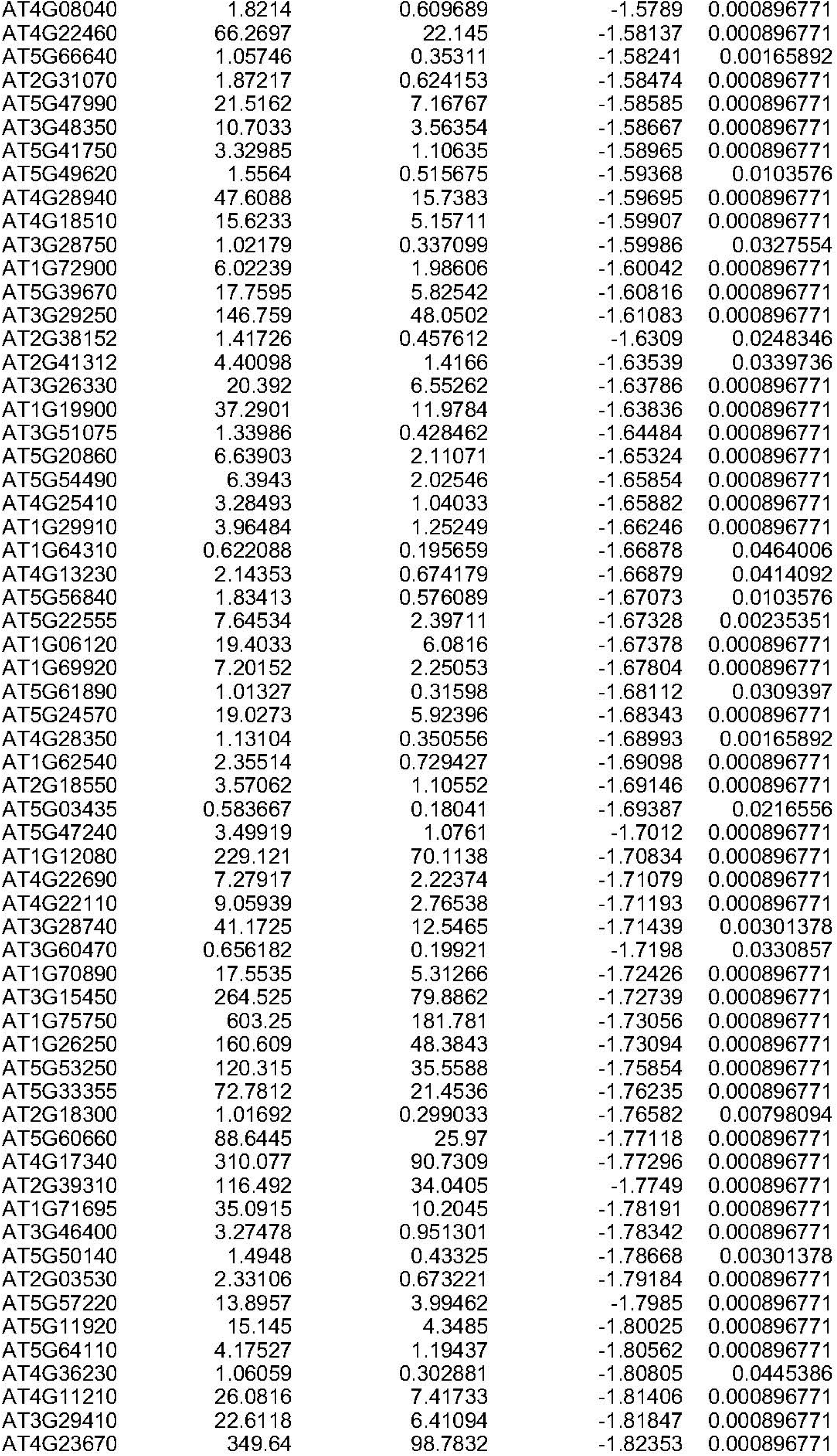
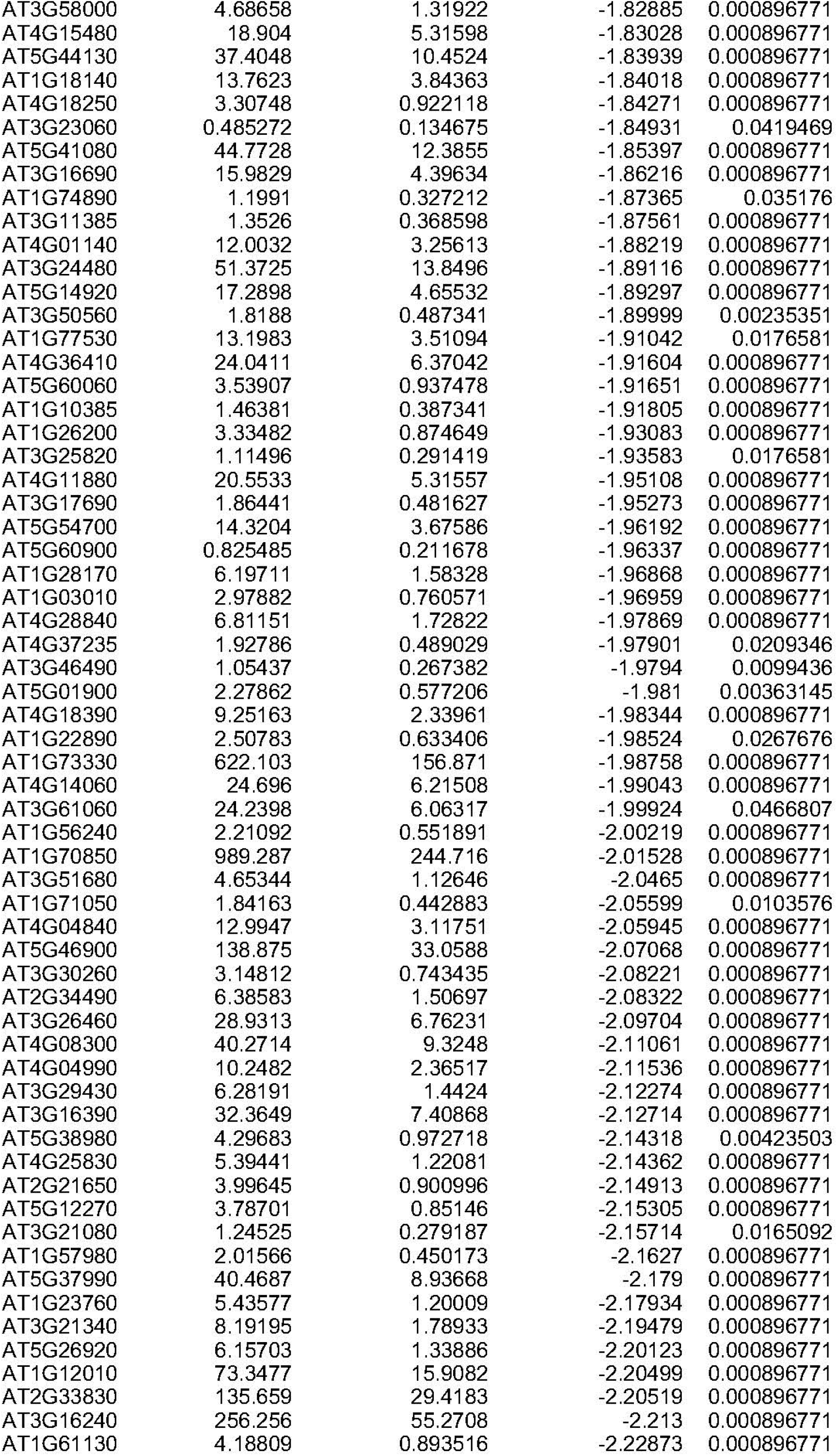
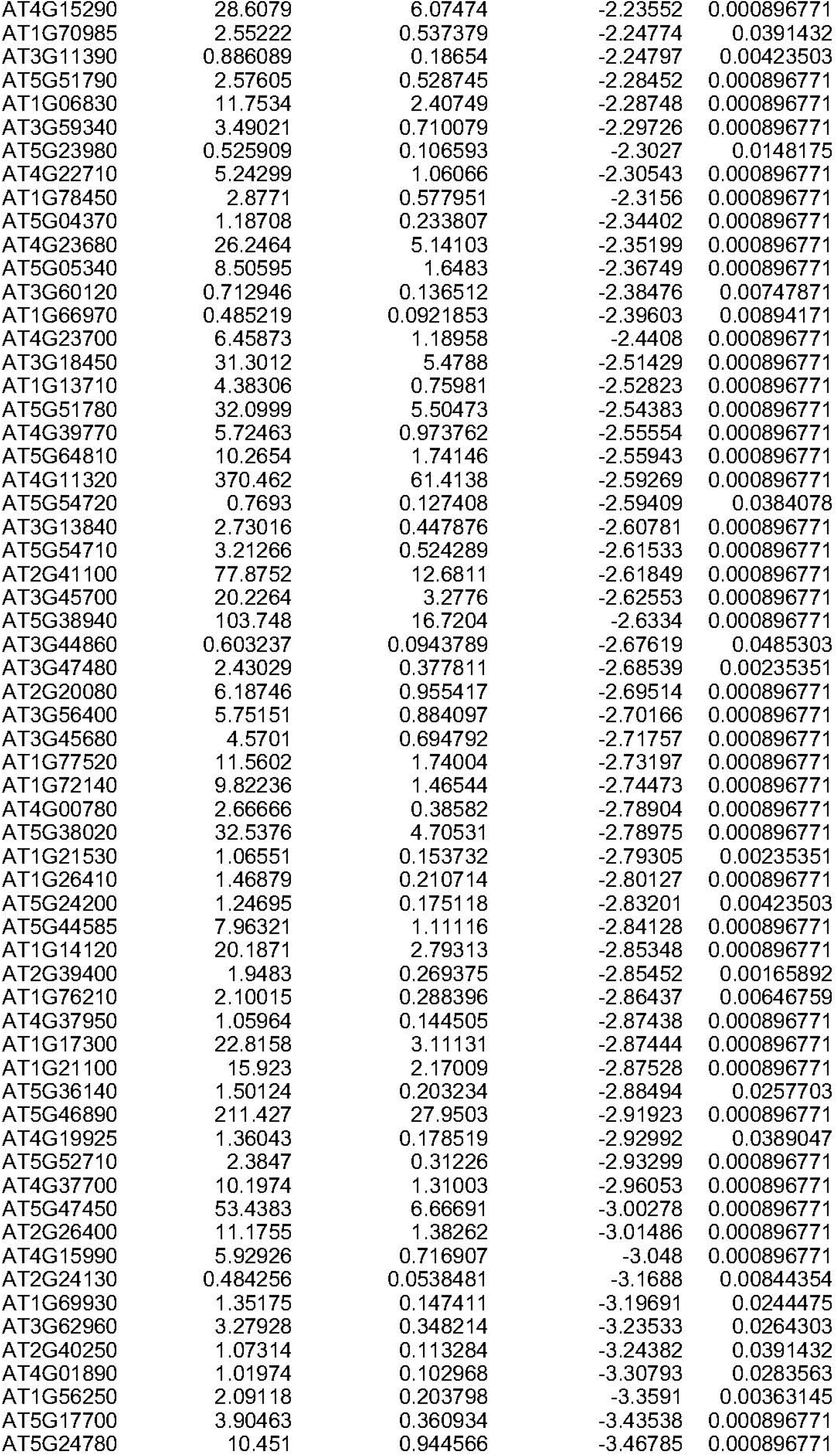
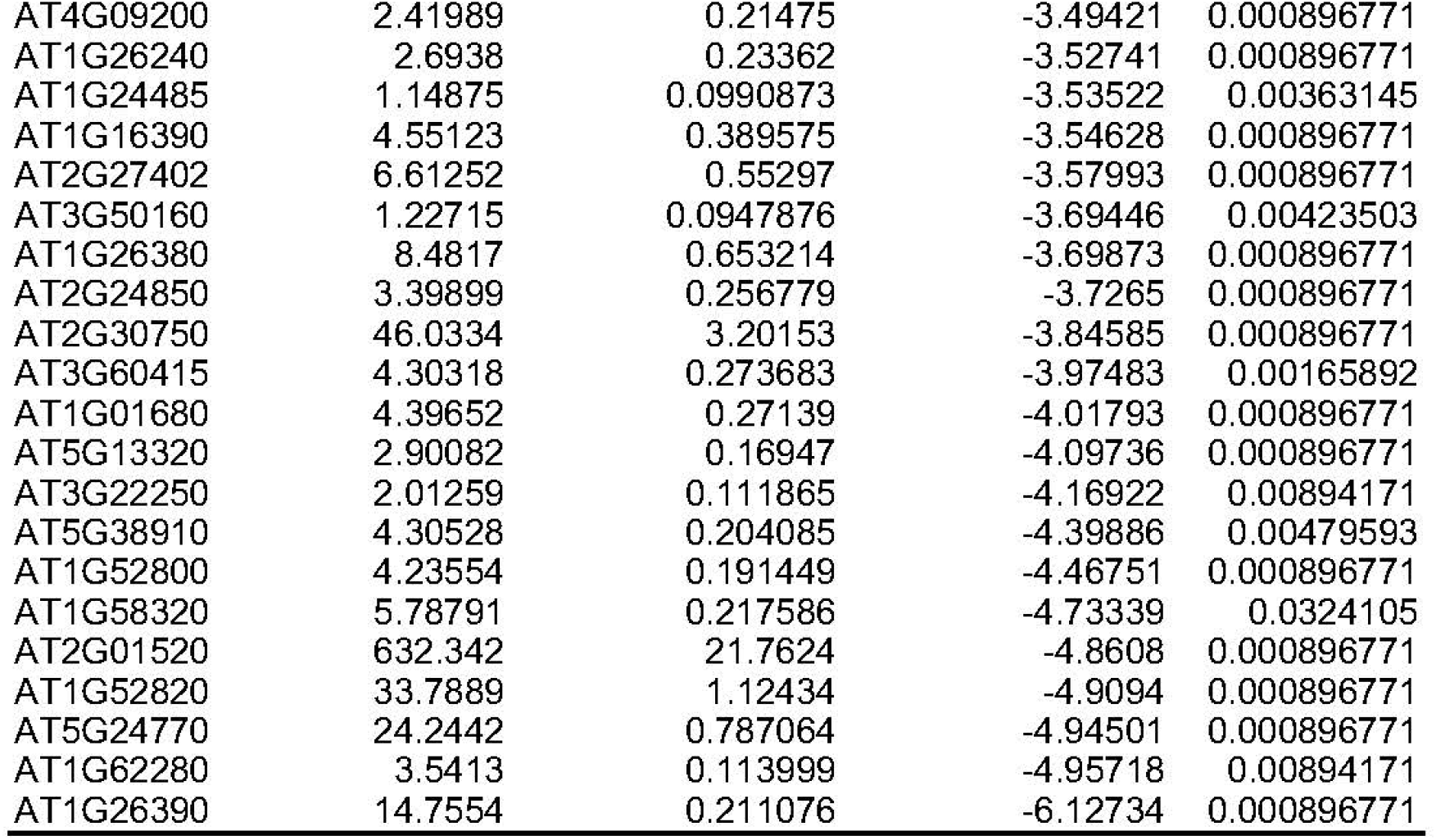
List of salt-responsive differentially expressed genes in the roots of WT.

**Table S4.**
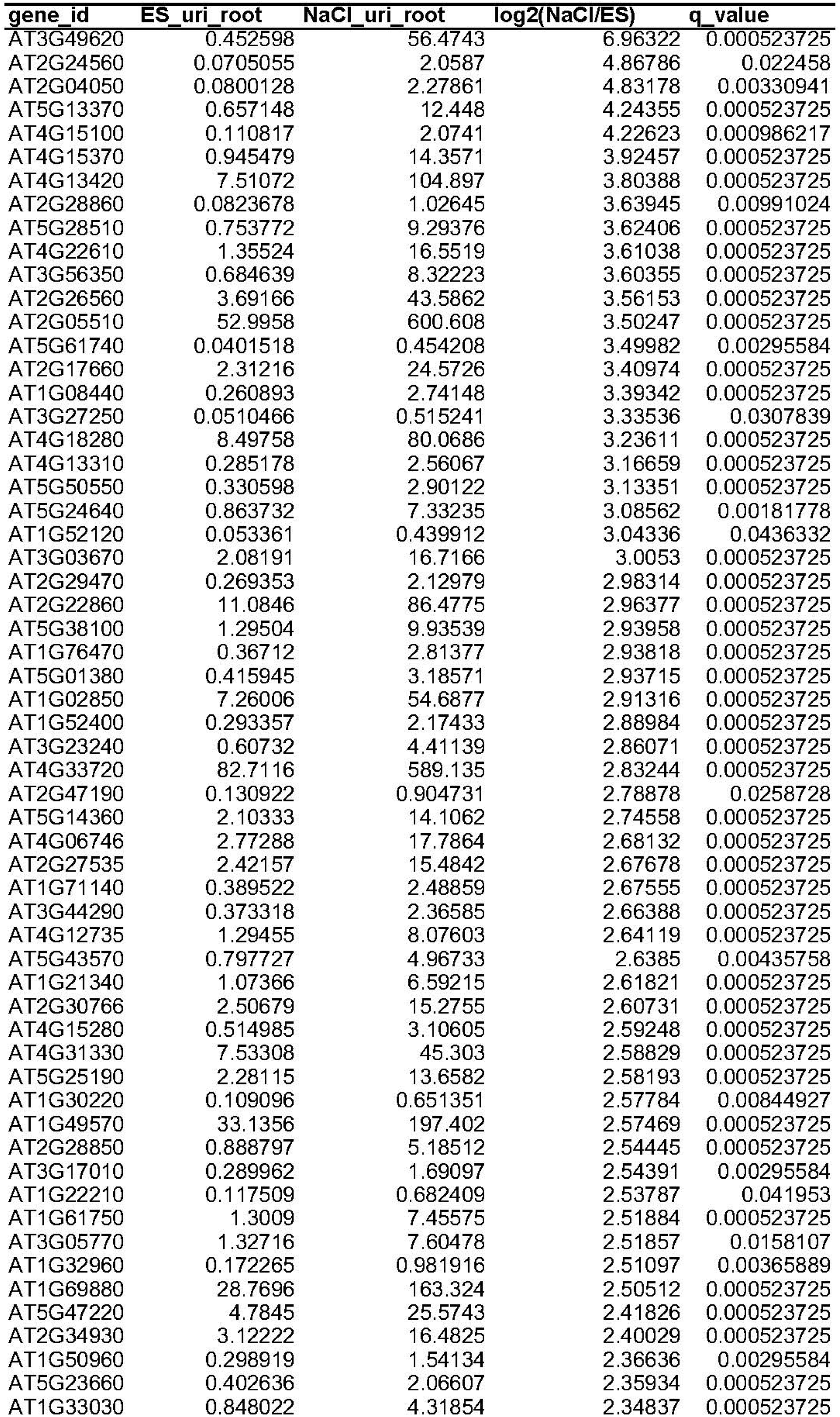
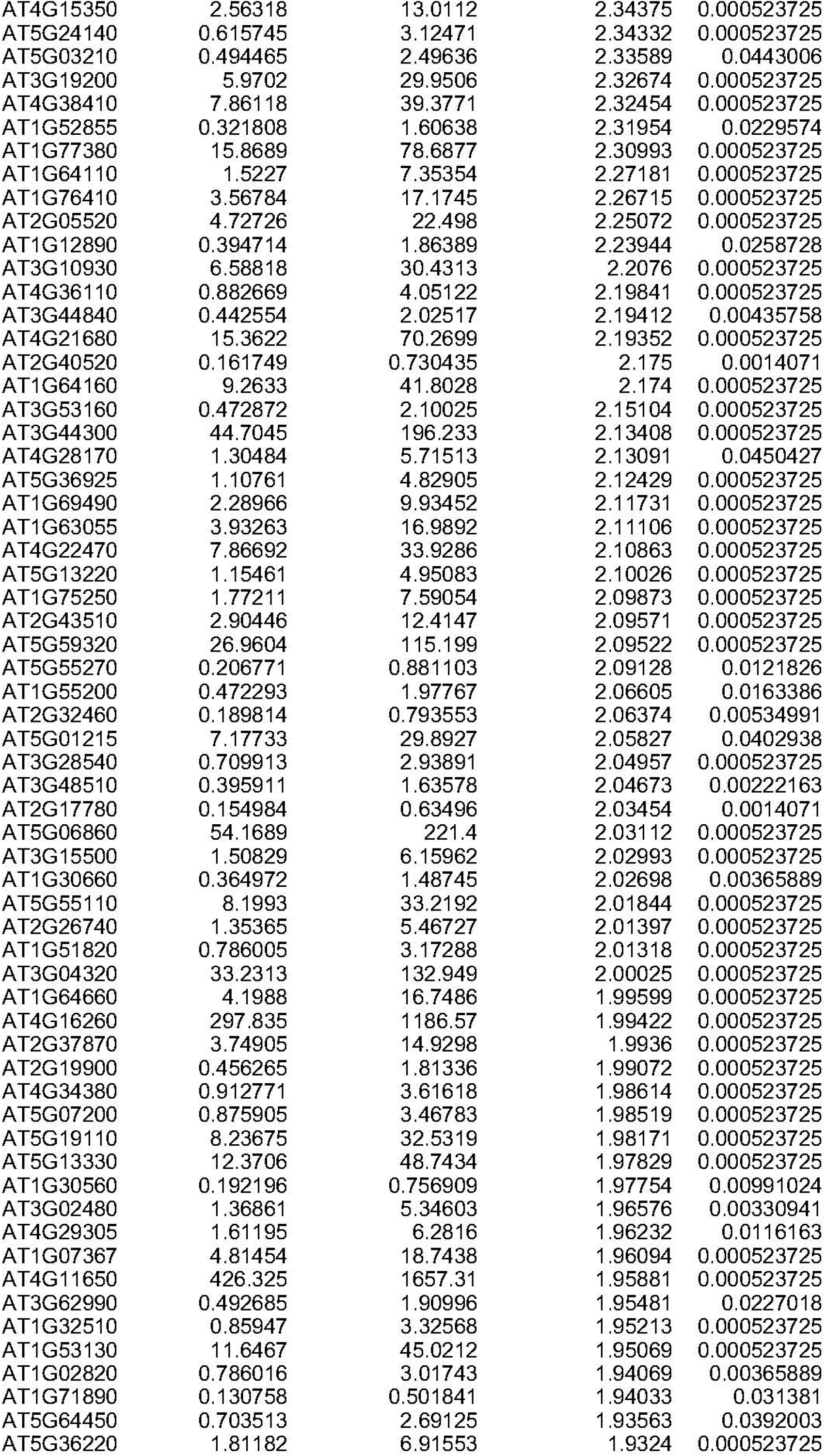
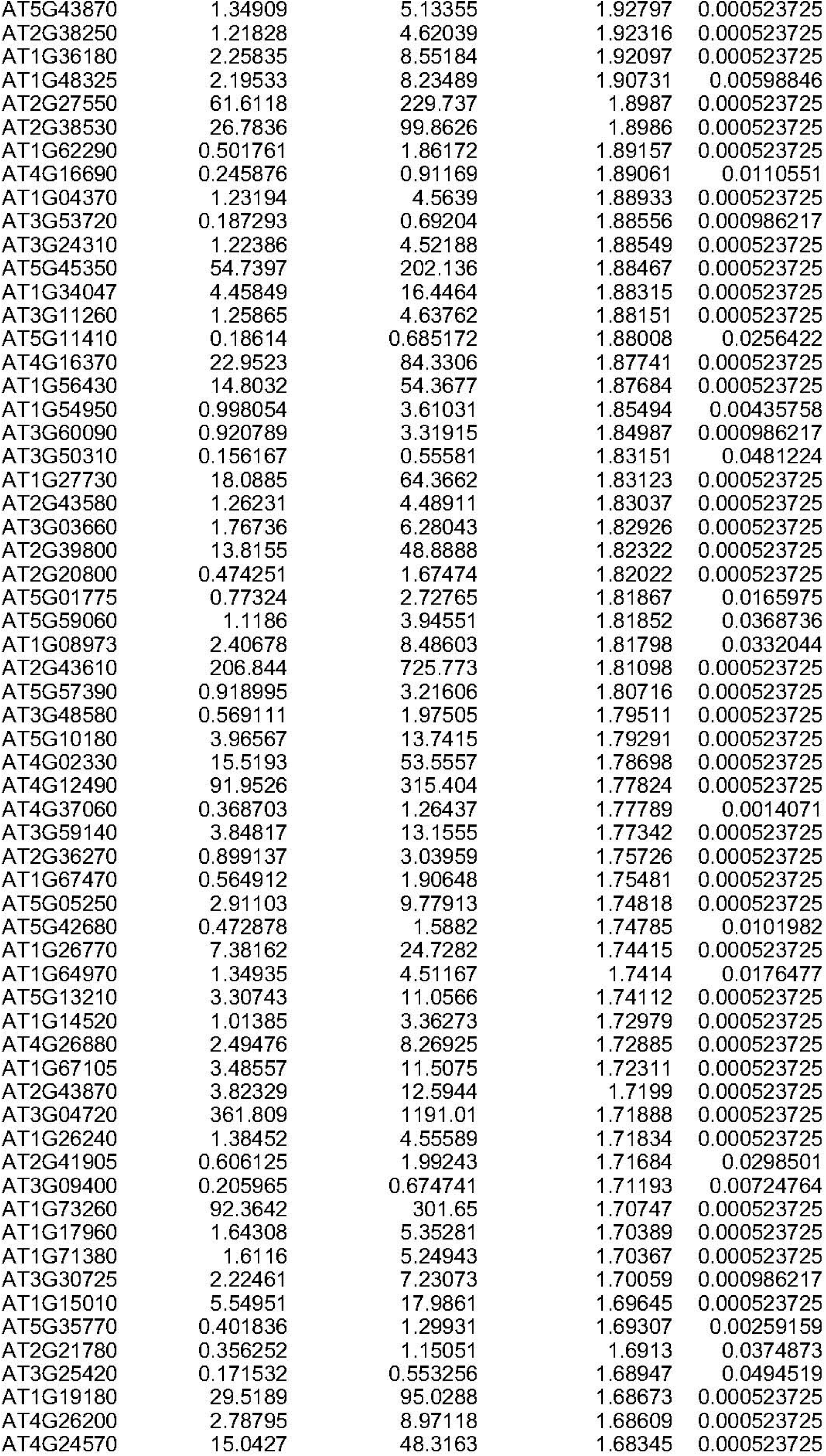
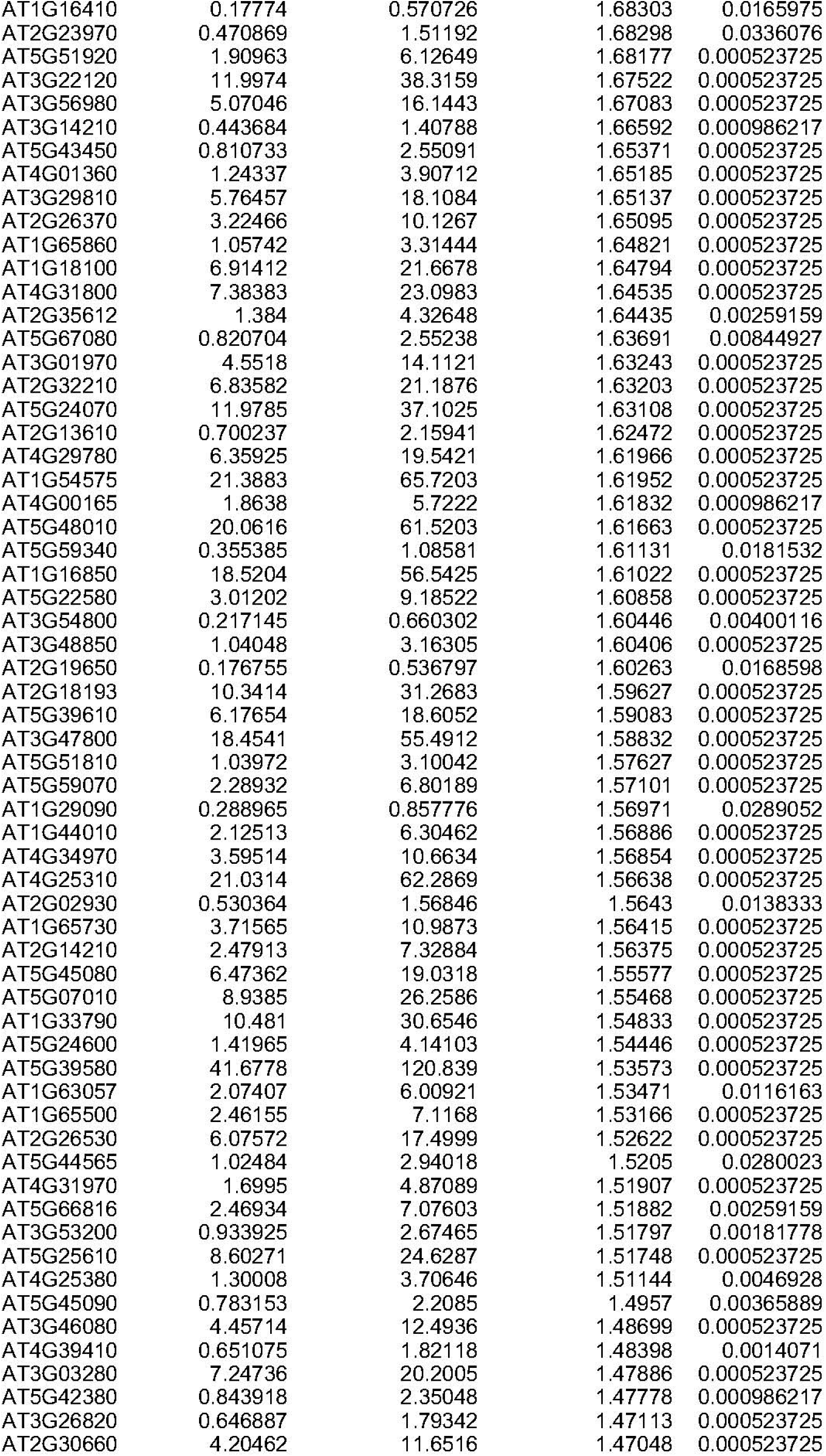
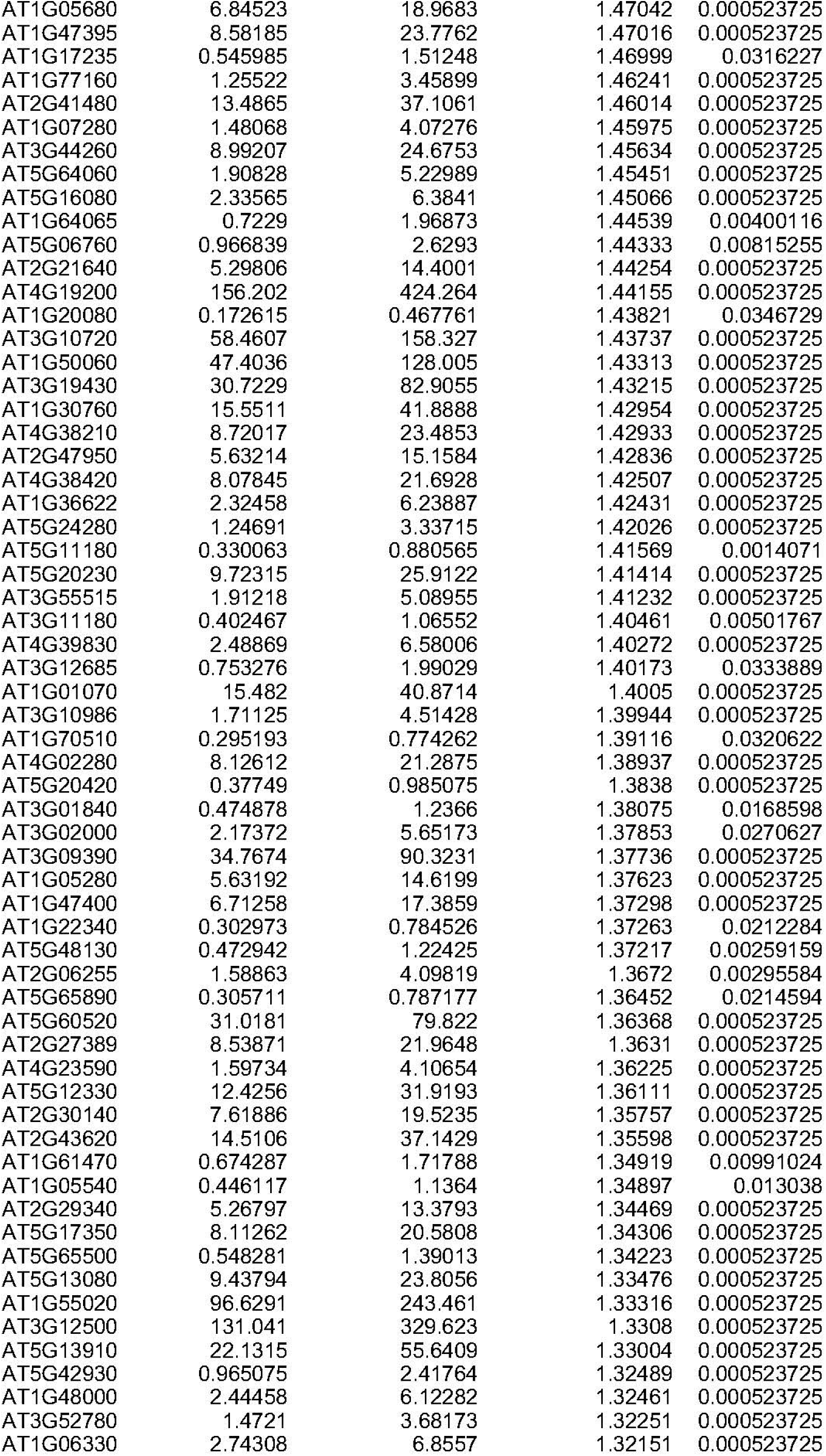
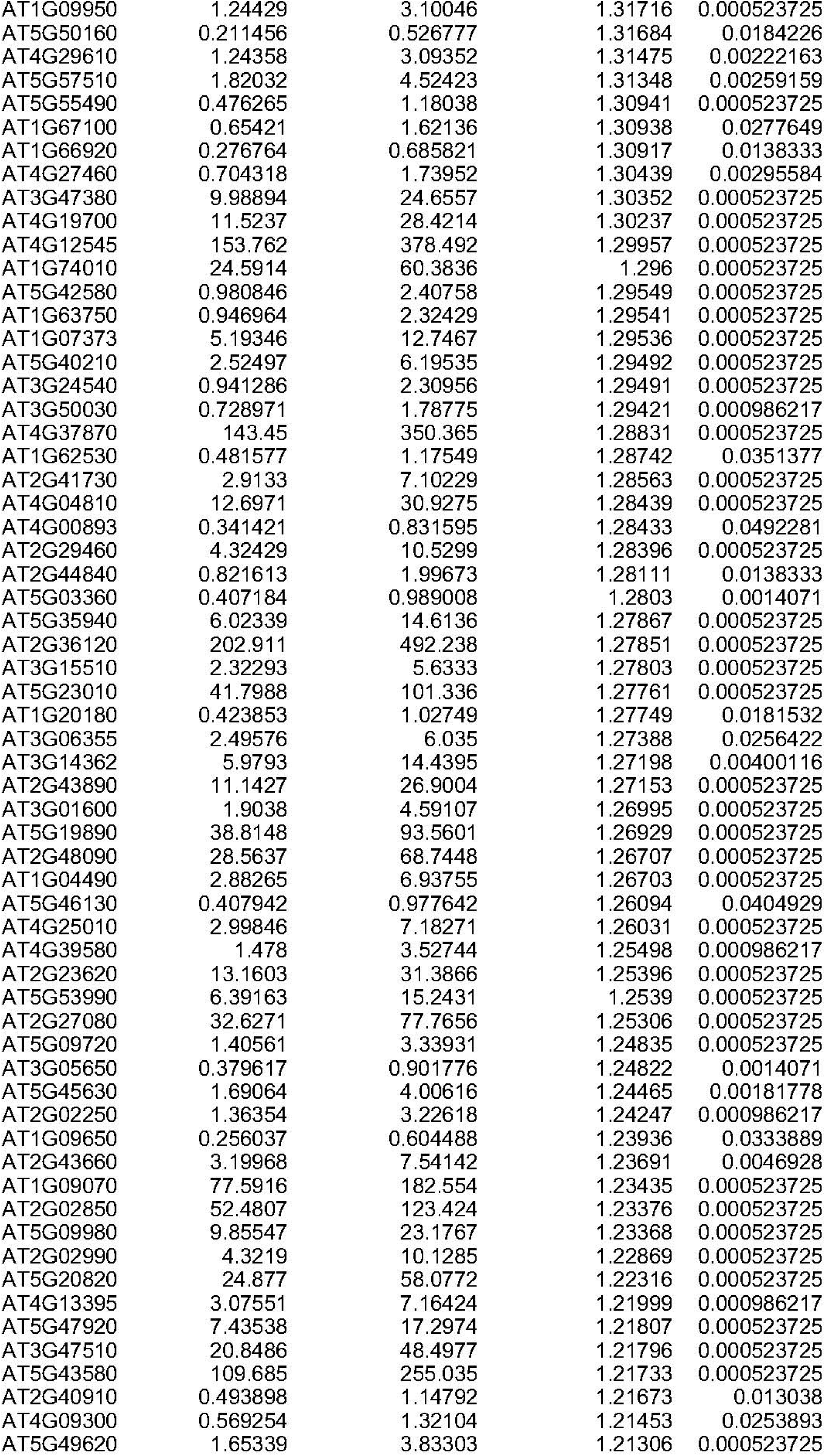
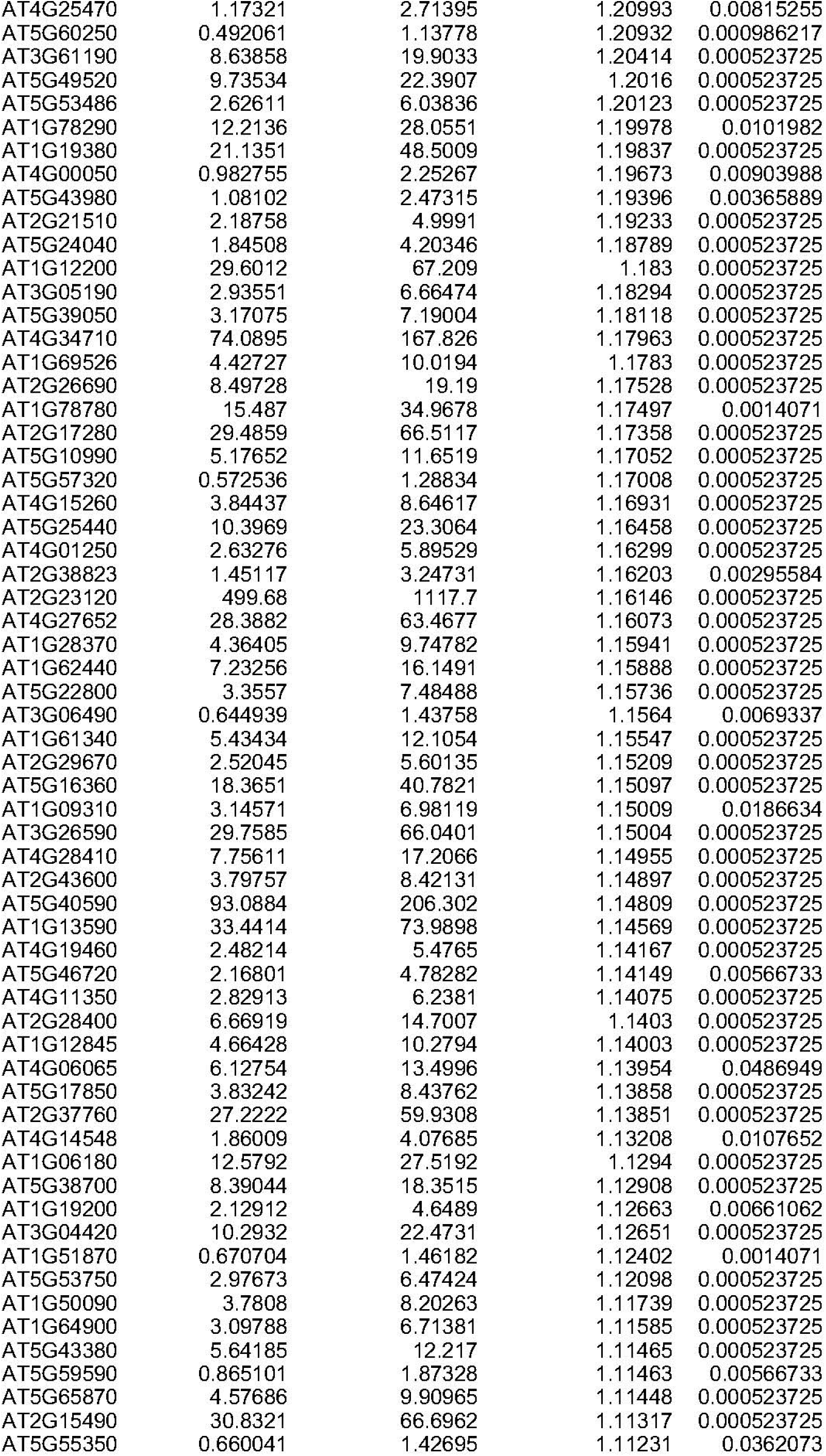
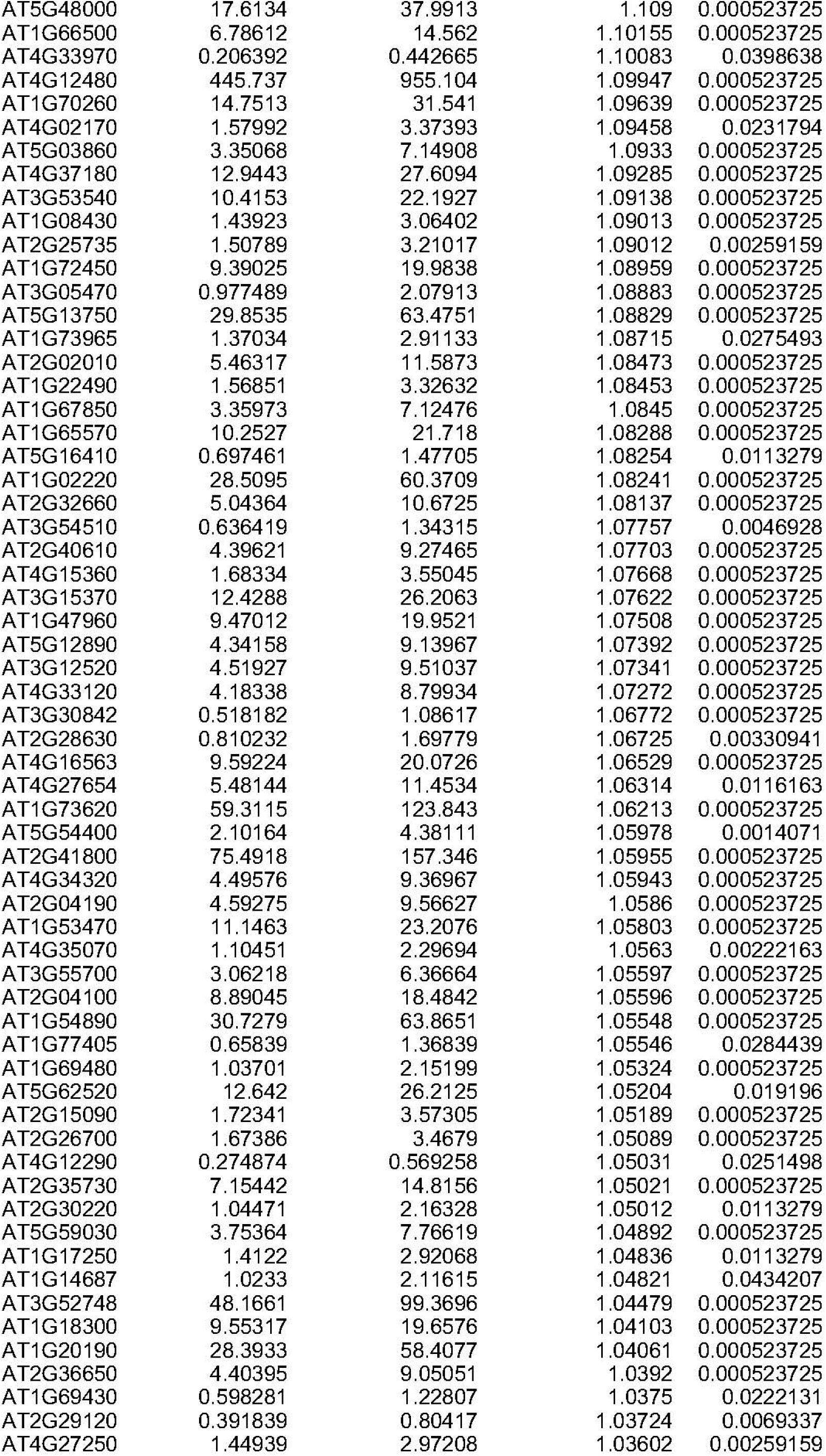
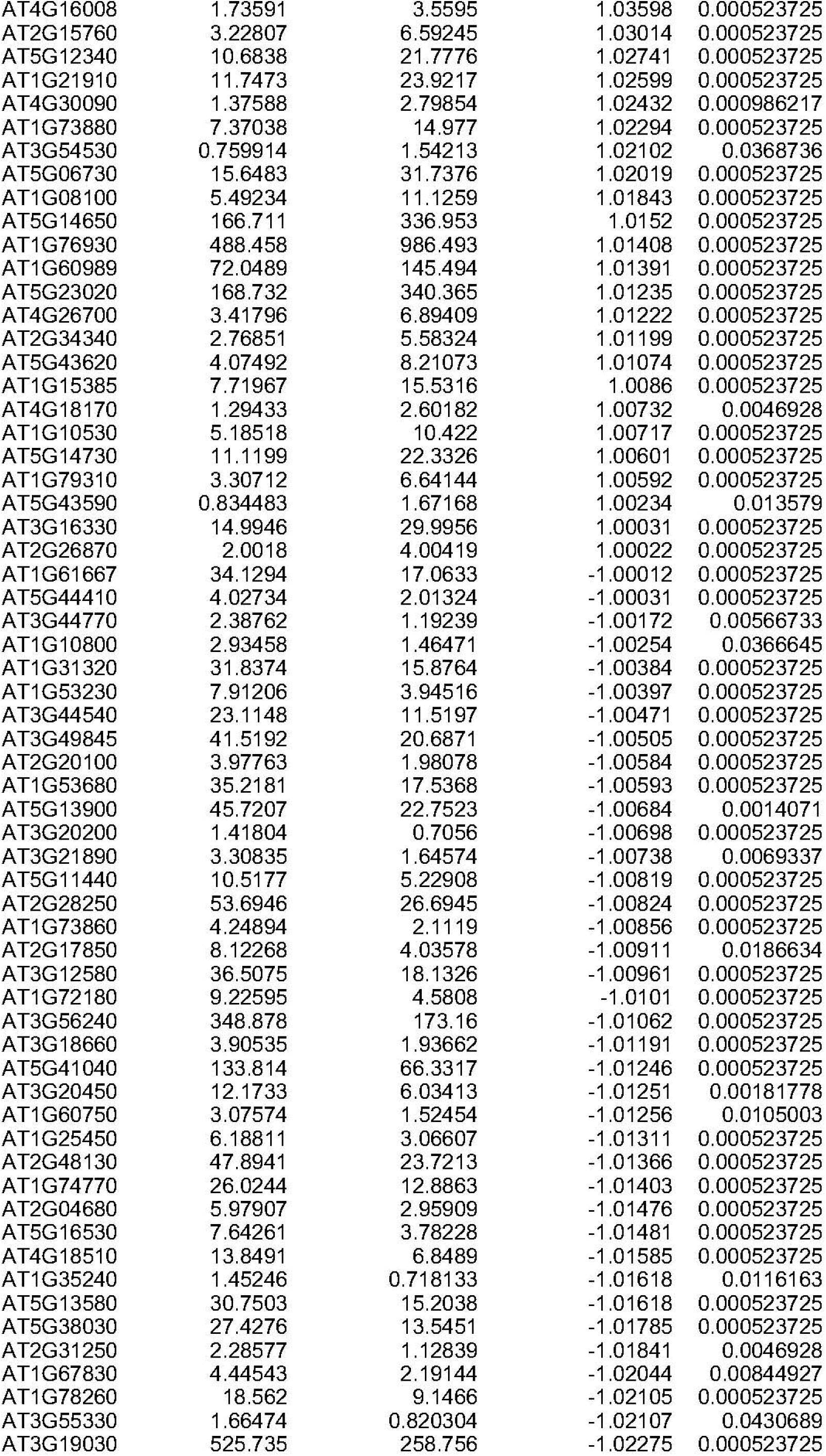
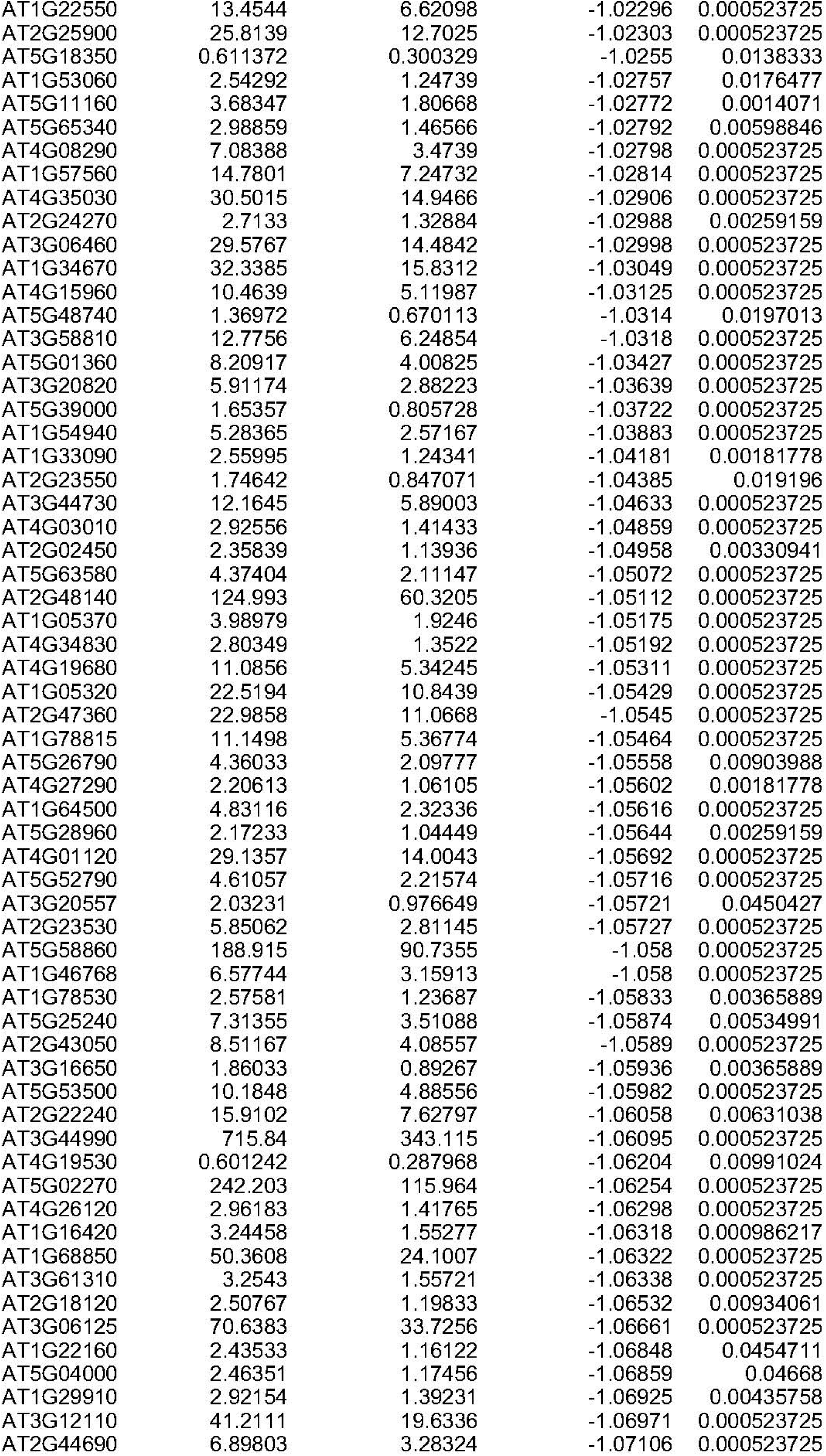
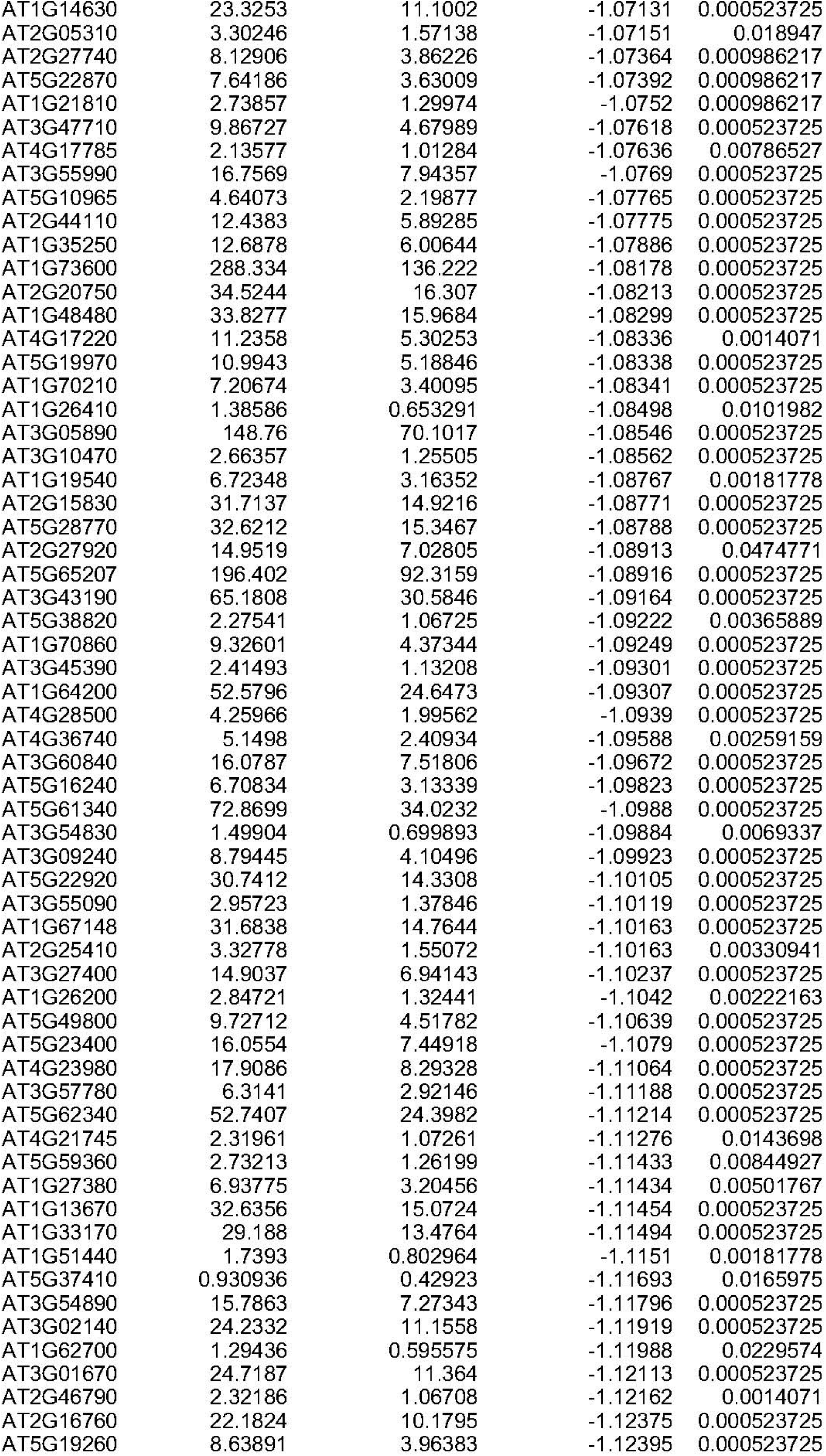
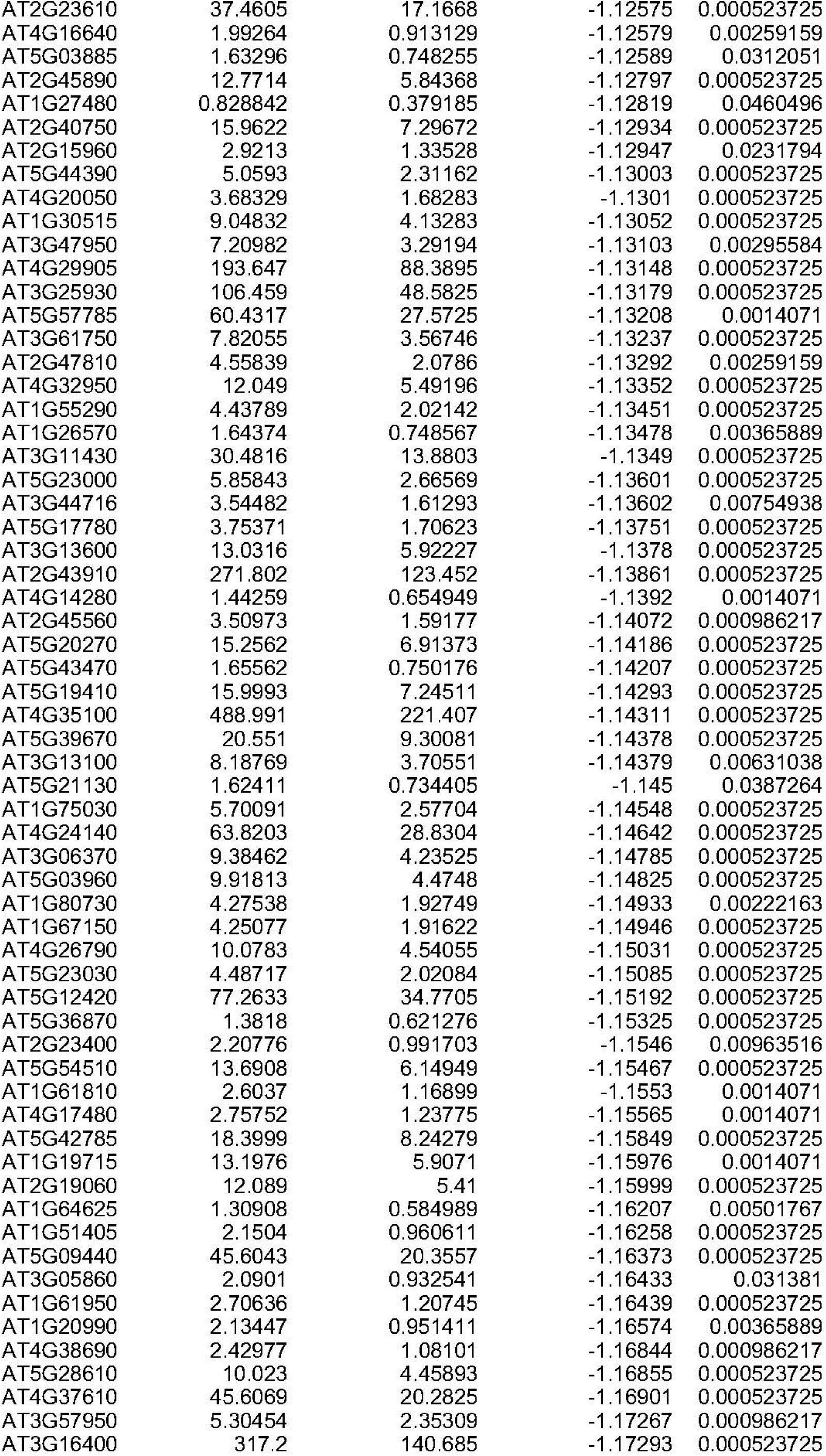
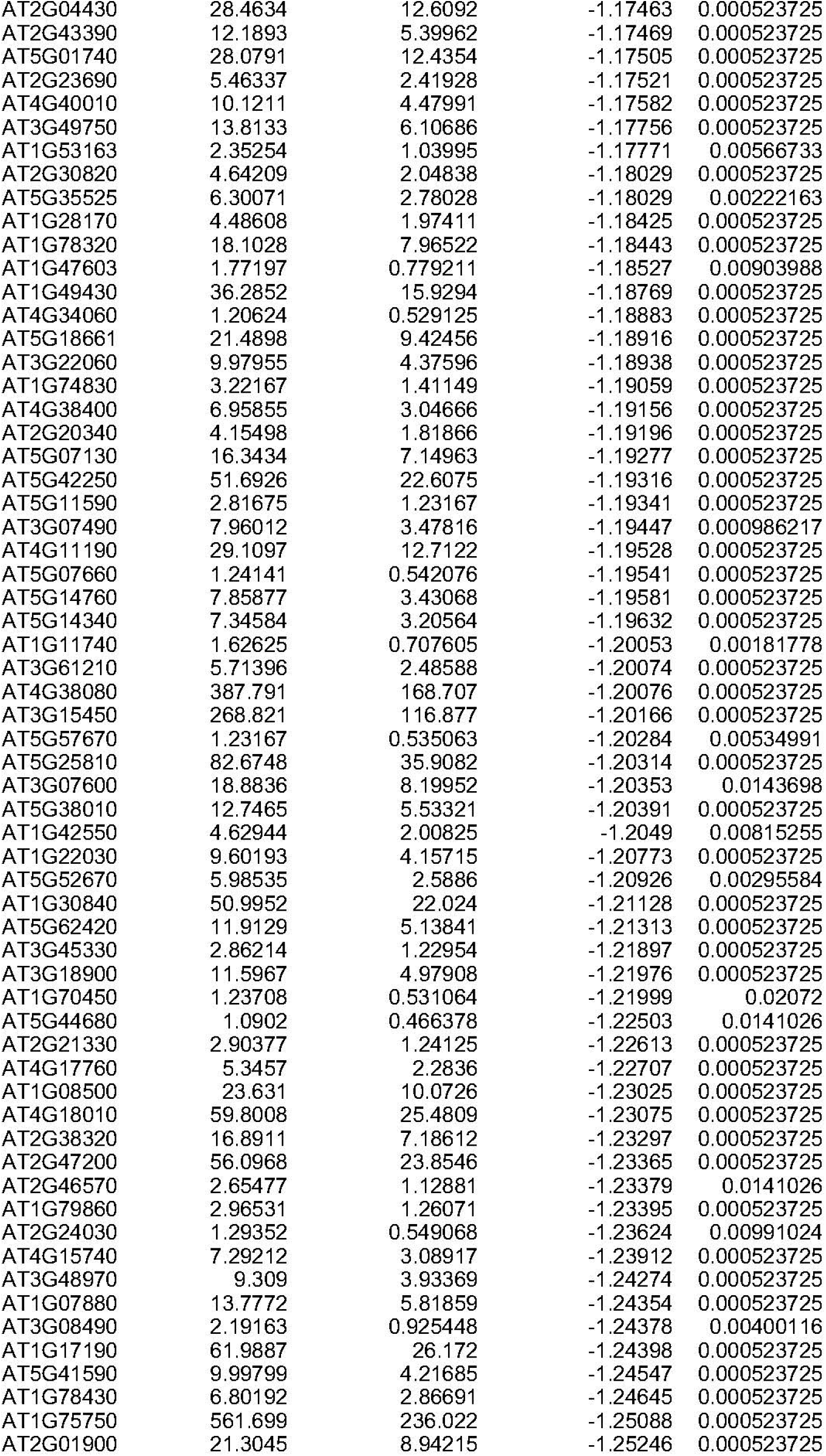
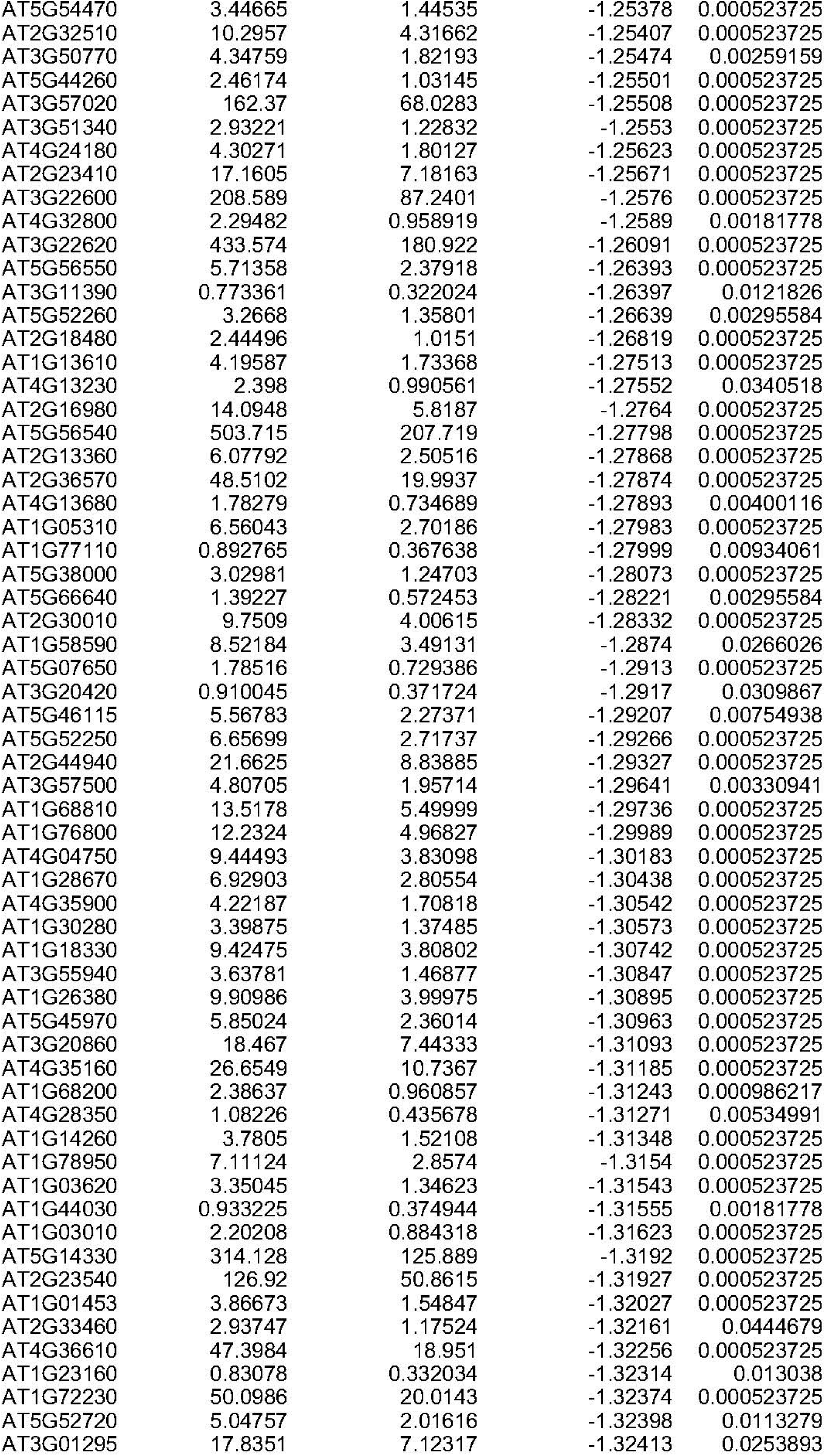
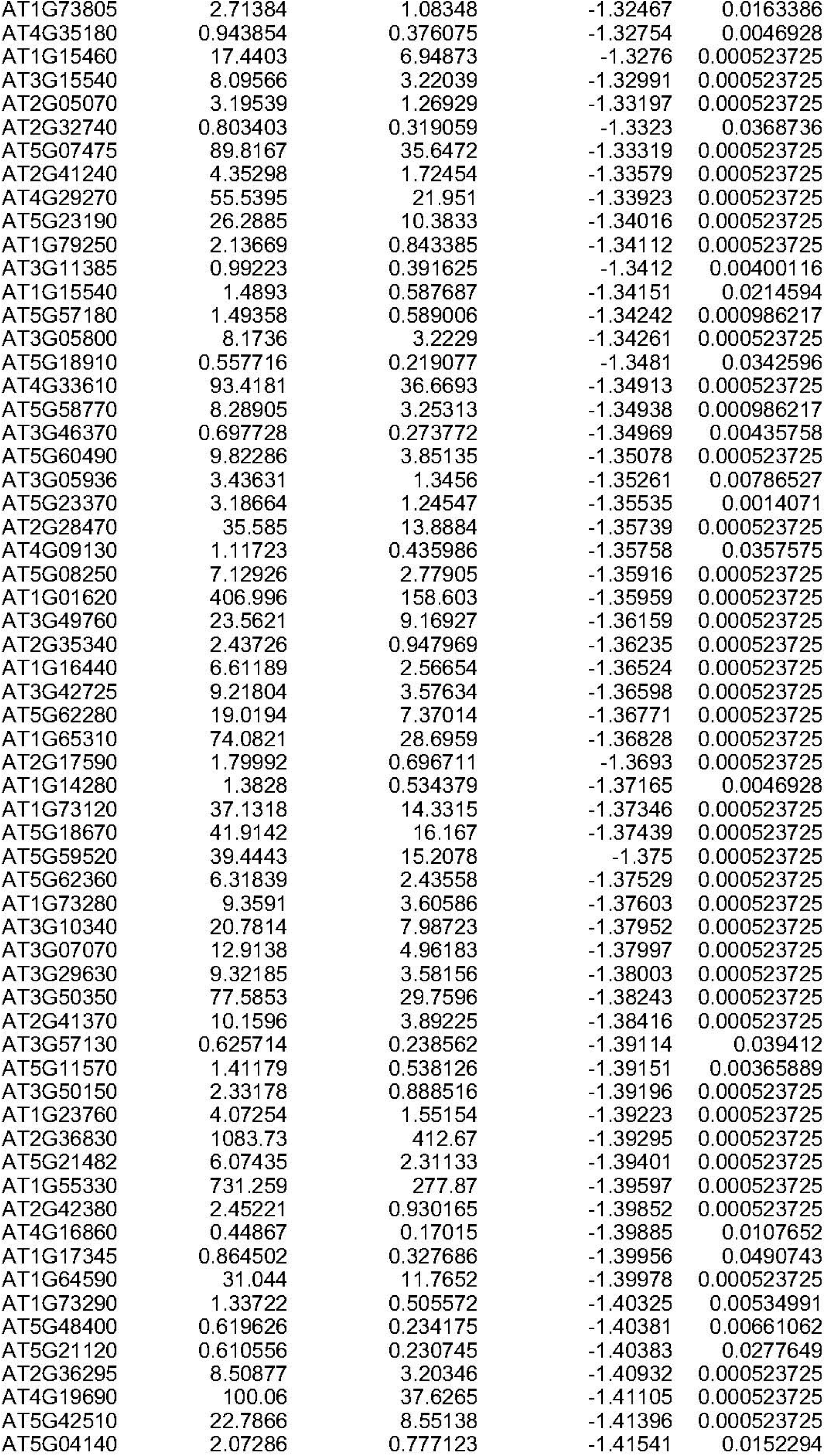
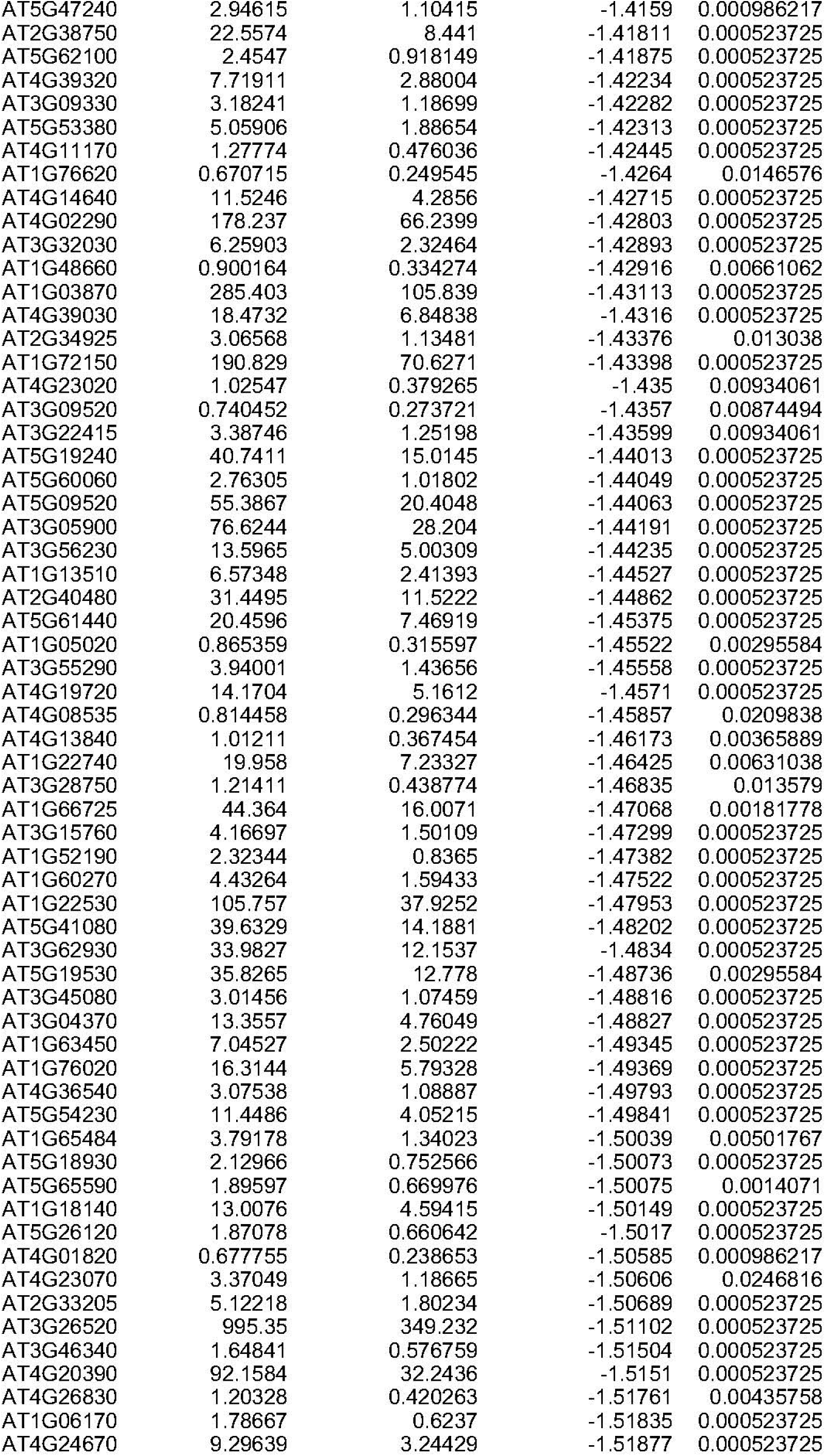
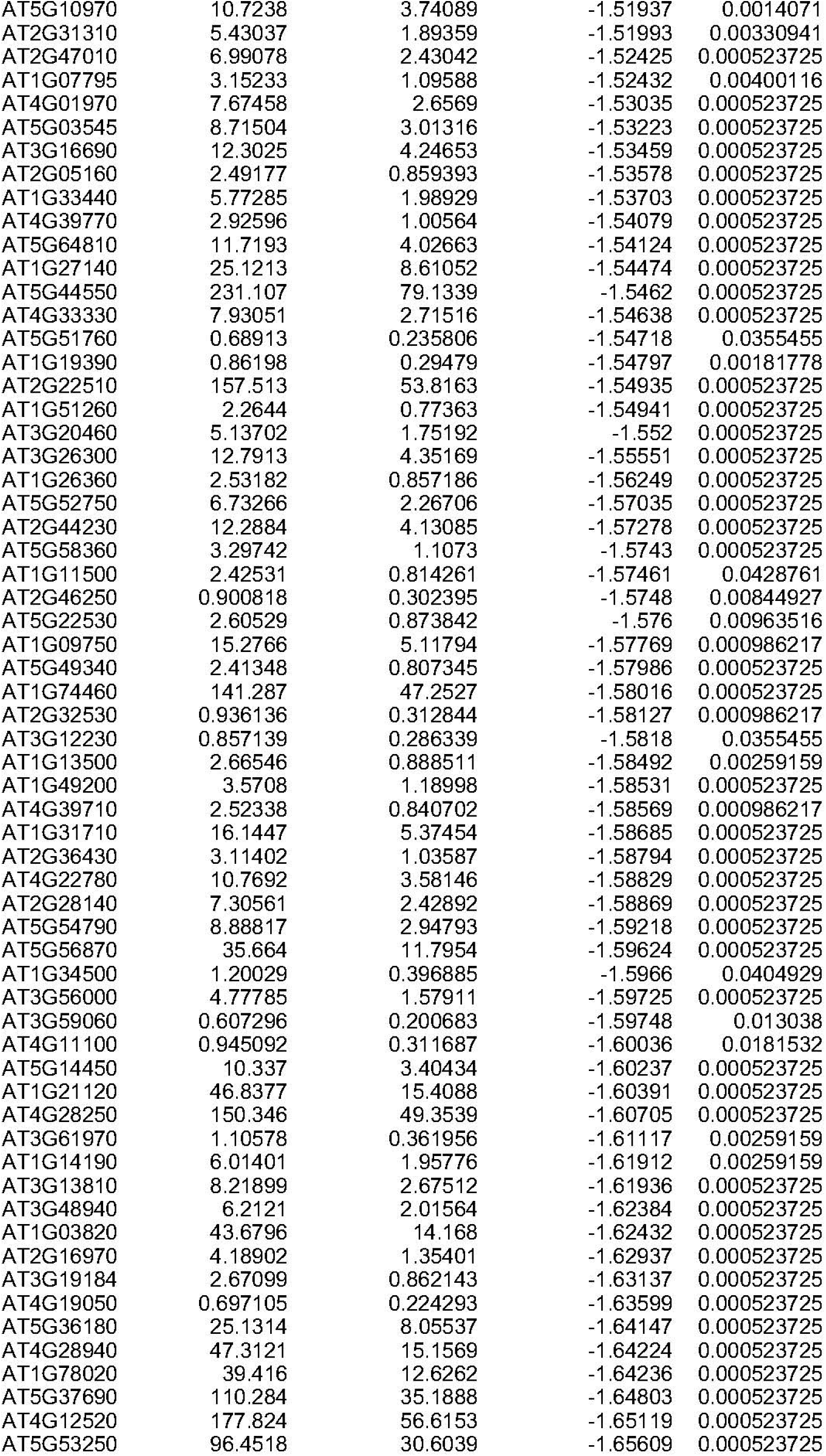
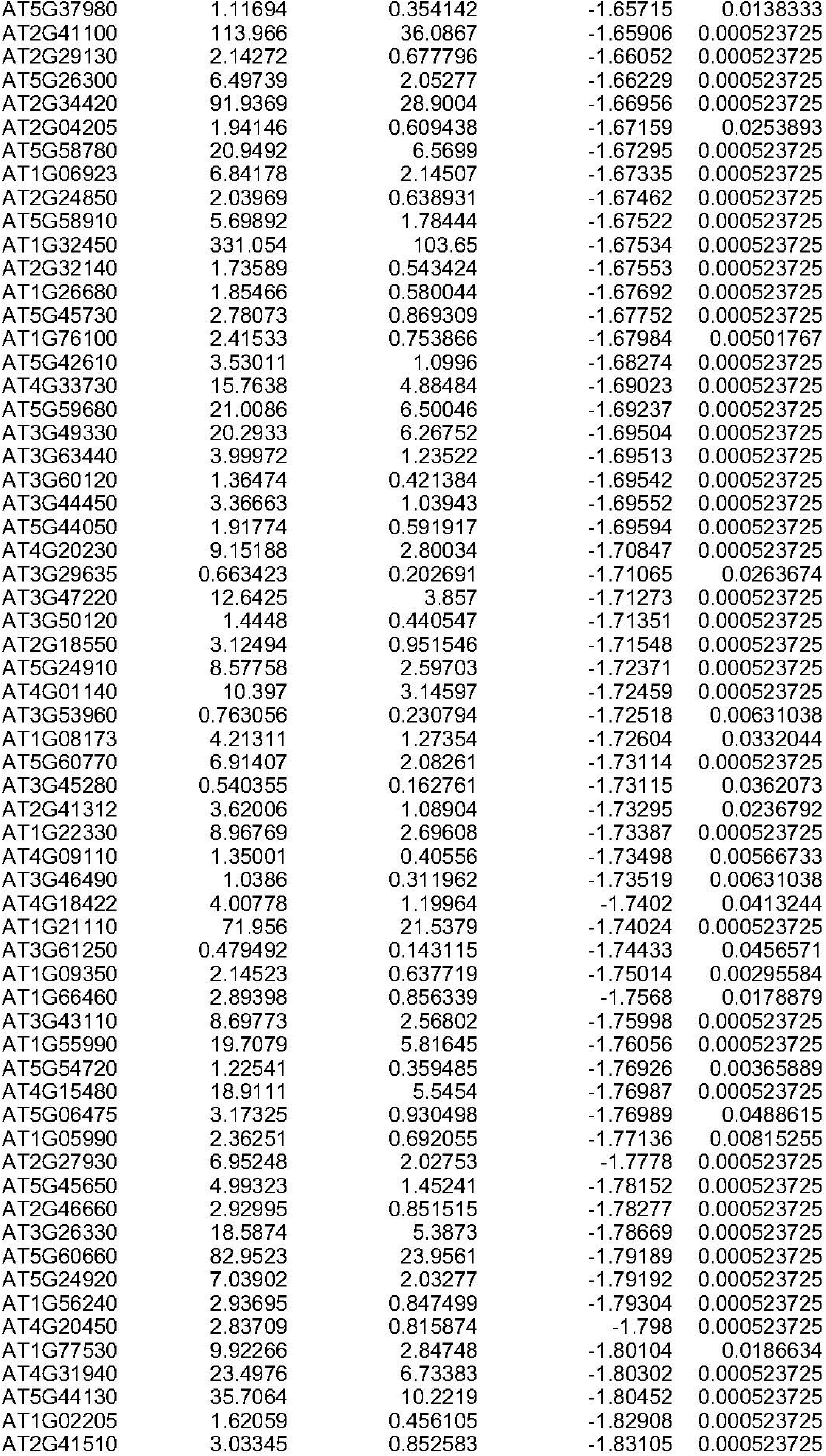
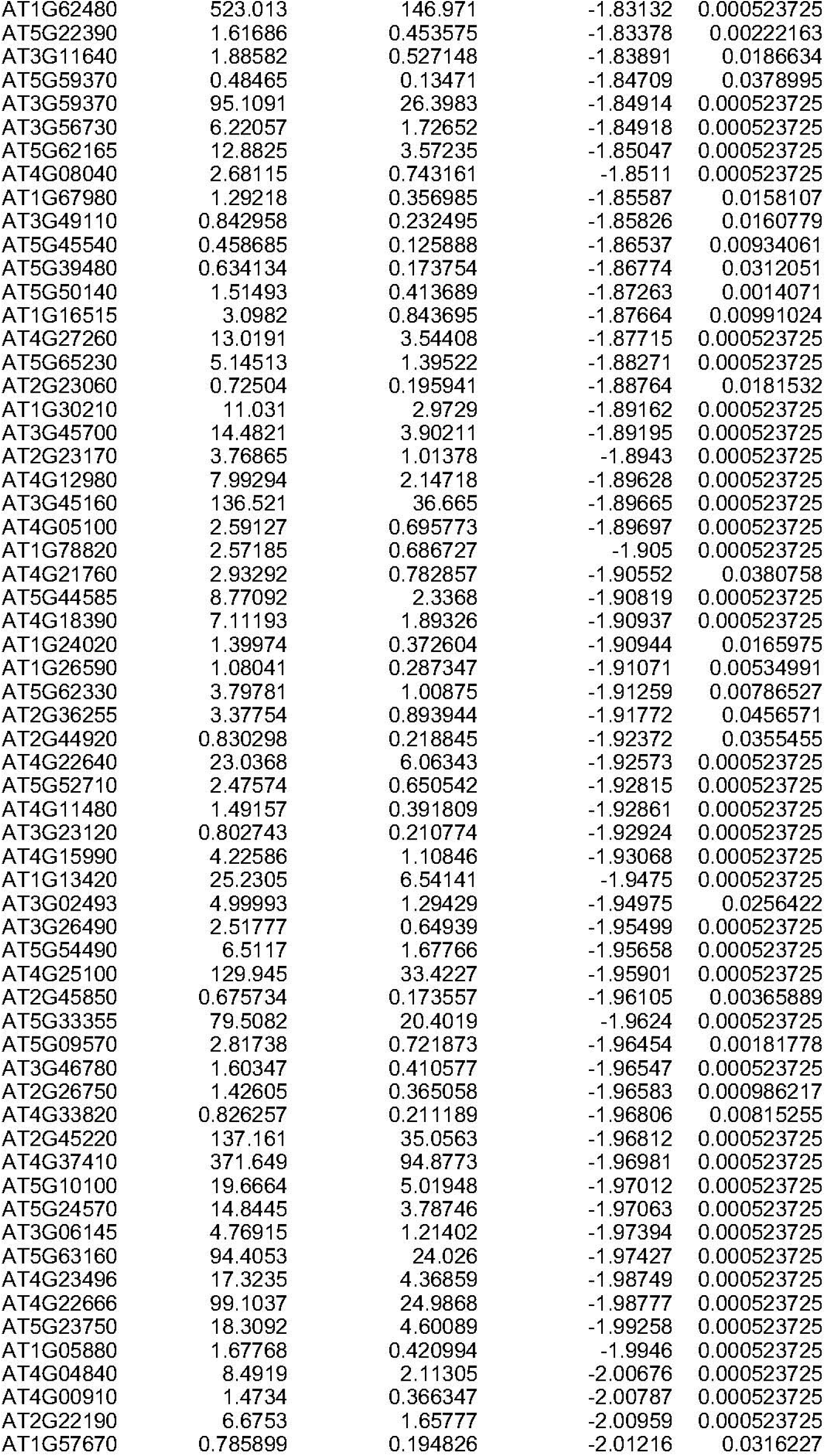
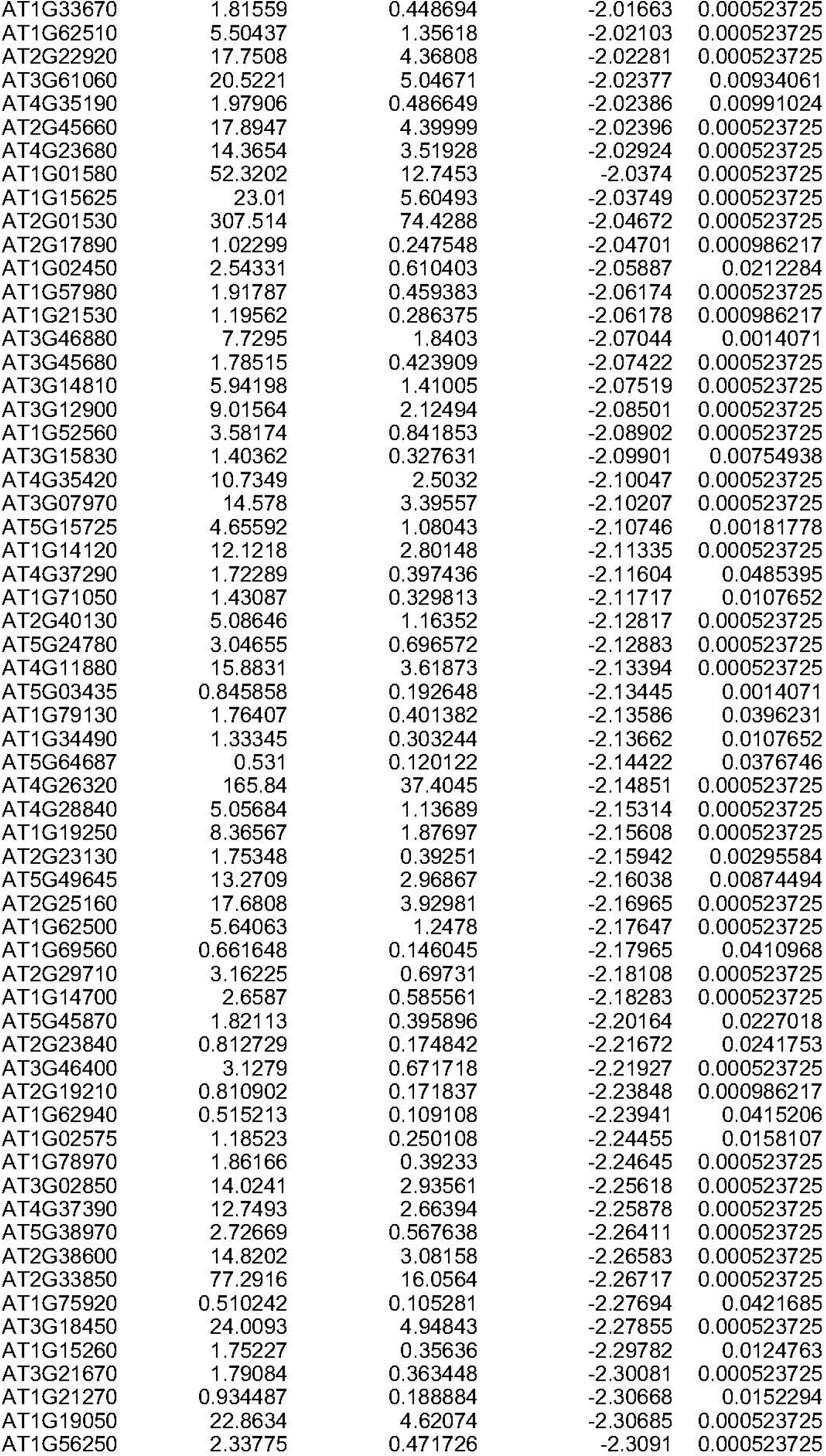
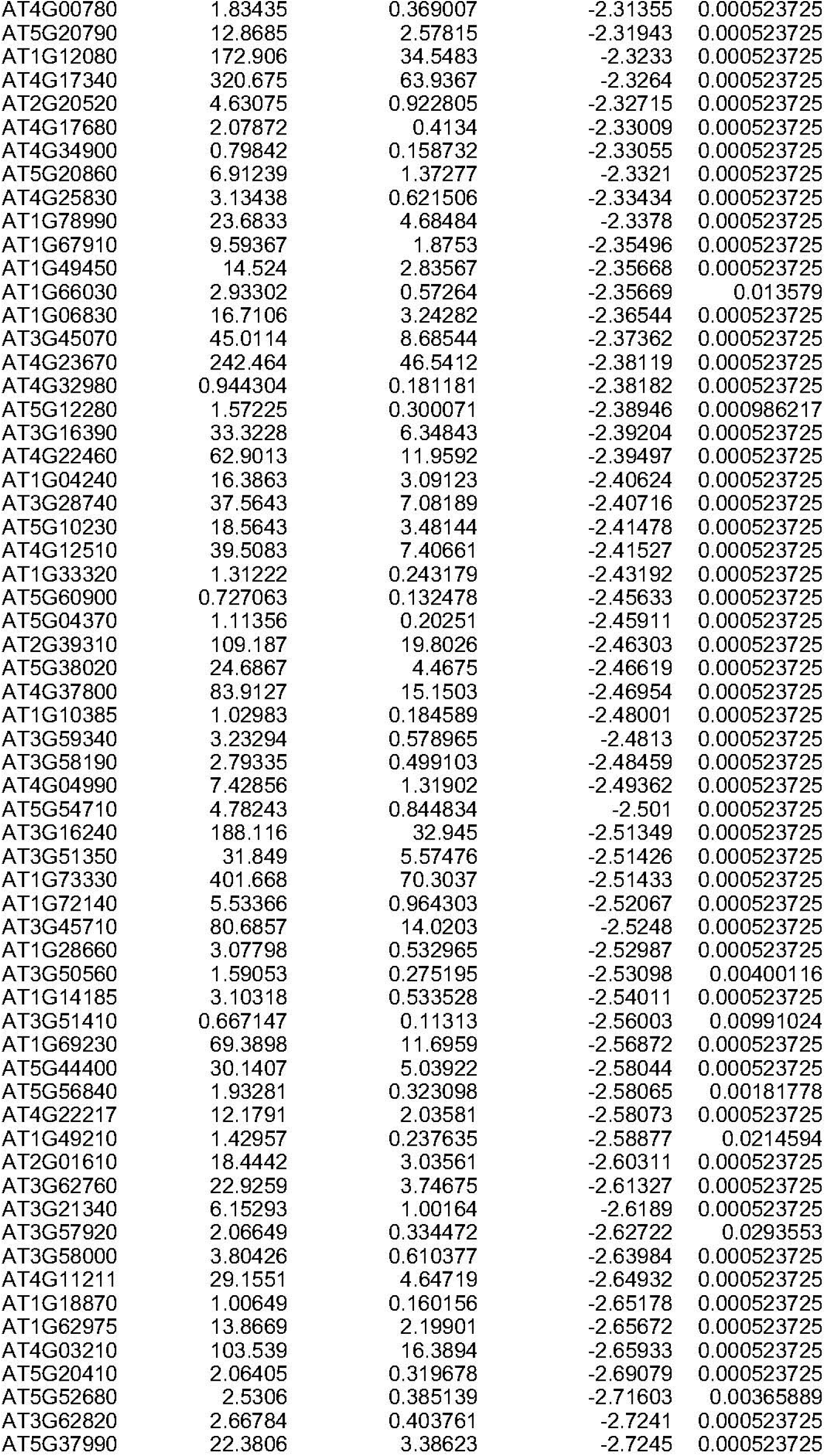
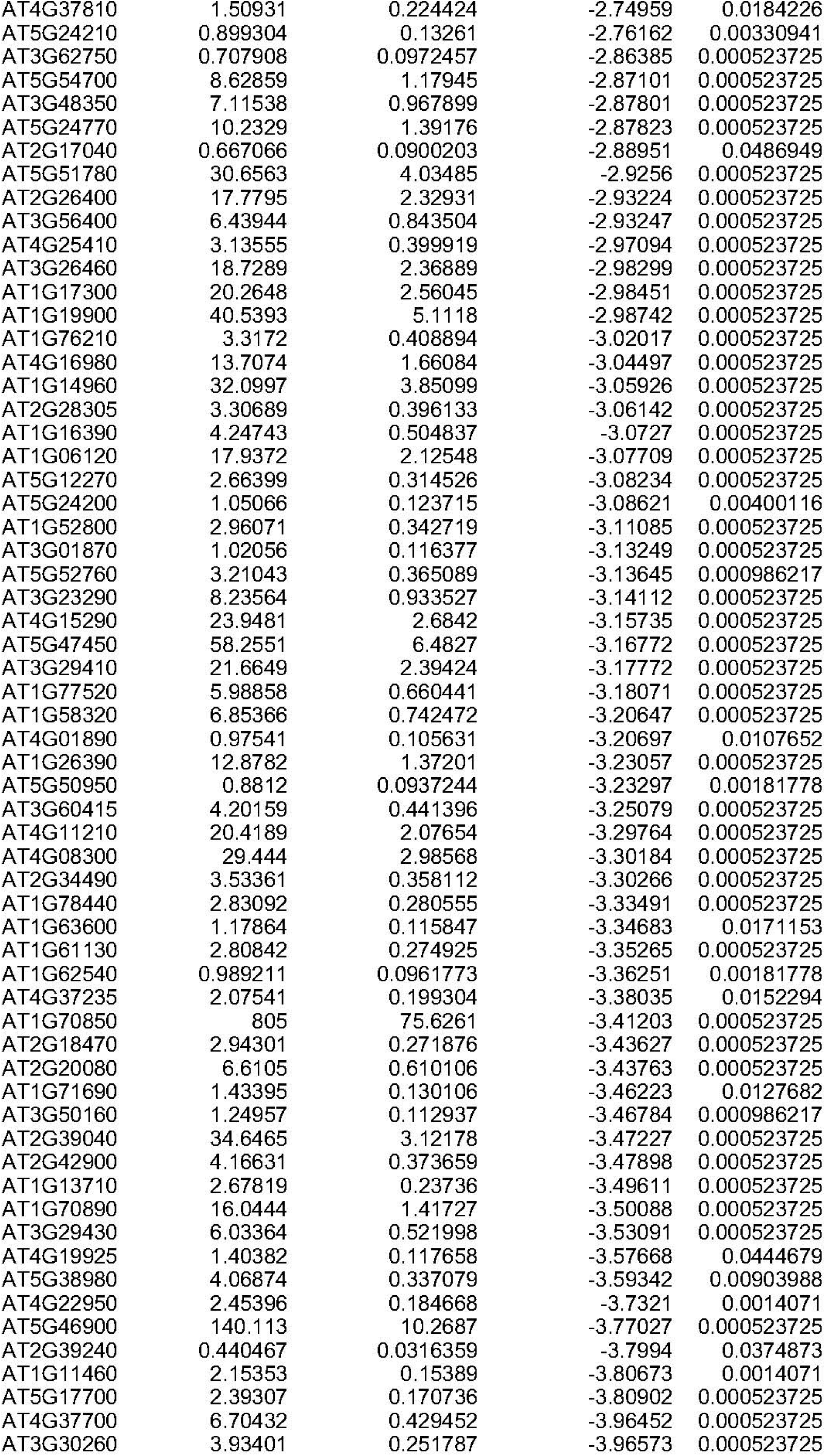
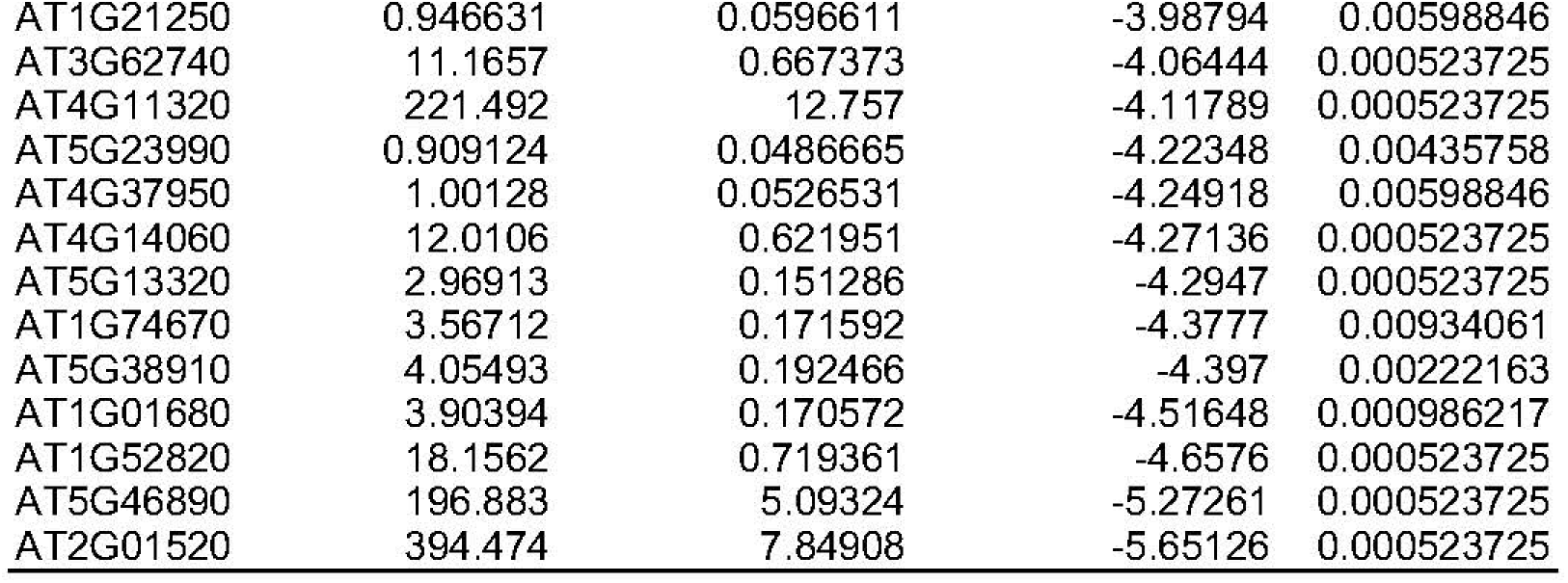
List of salt-responsive differentially expressed genes in the roots of *uri-2* mutant.

## Data availability statement

All data generated or analyzed during this study are included in this published article and online Supplementary data file. The raw data of mRNA-seq presented in this paper have been deposited in the NCBI BioProject database under accession PRJNA1301088.

